# BTP-7, a novel peptide for therapeutic targeting of malignant brain tumors

**DOI:** 10.1101/2020.05.21.106633

**Authors:** Niklas von Spreckelsen, Yarah Ghotmi, Colin M. Fadzen, Justin M. Wolfe, Nina Hartrampf, Charlotte Farquhar, Sonja Bergmann, Mykola Zdioruk, J. Roscoe Wasserburg, Emily Murrell, Fernanda C. Bononi, Leonard G. Luyt, Martine L. M. Lamfers, Keith L. Ligon, E. Antonio Chiocca, Mariano S. Viapiano, Bradley L. Pentelute, Sean E. Lawler, Choi-Fong Cho

**Affiliations:** Harvey Cushing Neuro-Oncology Laboratories, Department of Neurosurgery, Brigham and Women’s Hospital, Harvard Medical School, Boston, MA 02115, United States; Department of Chemistry, Massachusetts Institute of Technology, Cambridge, MA 02139, United States; Department of Neurosurgery, Center for Neurosurgery, Faculty of Medicine and University Hospital, University of Cologne, 50937 Cologne, Germany; Department of Chemistry, University of Zurich, Winterthurerstrasse 190, 8057 Zurich, Switzerland; Department of Chemistry, University of Western Ontario, London, ON N6C 4K3 Canada; Department of Neurosurgery, Brain Tumor Center, Erasmus Medical Center, Rotterdam, the Netherlands; Department of Pathology, Dana-Farber Cancer Institute and Brigham and Women’s Hospital, Harvard Medical School, Boston, MA 02115, United States; Department of Neuroscience and Physiology, State University of New York Upstate Medical University, Syracuse, New York 13210

**Author notes:** Correspondence to Dr. Choi-Fong Cho and Dr. Sean Lawler, 60 Fenwood Rd, Building of Transformative Medicine, Boston, Massachusetts 02115. **Authorship**: Conceptualization: C.-F.C., S.E.L., M.V. and B.L.P. Methodology and investigation: C-F.C., N.v.S., Y.G., C.F., J.W., N.H., C.F., S.B., M.Z., R. W., E. M., and F.B. Sample provision: K.L. and M.L. Manuscript drafting: C-F.C., N.v.S., Manuscript revision and editing: All authors.

**Keywords:** Peptide, Drug targeting, Glioblastoma, Peptide-drug conjugate

## Abstract

**Background:** Targeted therapies for malignant brain cancer that are currently available have little clinical activity, highlighting an urgent need for the development of novel precision medicines. Brevican (Bcan), a central nervous system (CNS)-specific extracellular matrix protein is upregulated in glioma cells. A brevican isoform lacking glycosylation, dg-Bcan, is a unique glioma marker and thus represents a valuable target for anti-cancer therapy. In this study, we aimed to find a versatile dg-Bcan specific ligand to facilitate glioma targeting.

**Methods:** We screened a D-peptide library to identify dg-**B**can-**T**argeting **P**eptide (BTP) candidates, which were characterized extensively through binding kinetic analyses, cell uptake tests and animal studies.

**Results:** The top candidate, BTP-7 binds dg-Bcan with high affinity and specificity, is preferentially internalized by Bcan-expressing glioma cells and can cross the blood-brain barrier *in vitro* and in mice. Functionalization of camptothecin with BTP-7 led to increased drug delivery to intracranial glioblastoma and cytotoxicity in tumor tissues, as well as prolonged survival in tumor-bearing mice.

**Conclusion:** dg-Bcan is an attractive therapeutic target for high-grade gliomas, and BTP-7 represents a promising lead candidate for further development into novel targeted therapeutics.

**Key points:**

1. BTP-7 is a high affinity peptide ligand for the dg-Bcan protein and Bcan-expressing cells.
2. BTP-7 targets human intracranial GBM xenografts in mice.
3. Functionalization of a toxic anti-cancer drug with BTP-7 enables targeted delivery of the therapeutic to intracranial GBM in mice

**Importance of the Study:** Targeted therapies for malignant brain cancer that are currently available have little clinical activity, highlighting an urgent need for the development of novel precision medicines that can selectively recognize and kill high-grade glioma tissues. A protein called dg-Bcan is an ideal target because it is present only in the extracellular matrix of high-grade glioma cells and is absent from normal brain tissues. Here, we describe the discovery of a novel dg-**B**can-**T**argeting **P**eptide, called BTP-7 that can bind specifically to high-grade glioma cells/tissues, and thus serve as a promising drug delivery vehicle.

## Introduction

High-grade gliomas (HGG) are incurable with dismal patient prognosis even after surgery and chemo-radiotherapy.^1^ Invasive tumor cells are protected from most systemic chemotherapeutics by the blood-brain-barrier (BBB)^2^ and potent chemotherapeutics can also have off-target effects leading to toxicity in healthy brain tissues.^3^ These roadblocks limit progress in the field and highlight the critical need for novel therapies with improved brain-delivery and tumor-recognition capabilities that can effectively destroy glioma cells while sparing healthy tissues.

While HGG are well-known for their cellular and genetic heterogeneity, the tumor extracellular matrix (ECM) has less spatial variability. Therefore, strategies to target the tumor ECM may confer advantages over conventional targeted-therapies that are specific for membrane receptors on select tumor cell populations like epidermal growth factor receptor variant III (EGFRvIII).^4,5^ Brevican (Bcan), a major ECM glycoprotein expressed exclusively in the central nervous system (CNS),^6^ is upregulated in HGG,^7,8^ and is associated with increased tumor invasion and aggressiveness.^9^ Although brevican has multiple isoforms in normal brain, a unique underglycosylated isoform (named dg-Bcan) has only been found in human HGG.^7^ Importantly, dg-Bcan is absent in the adult CNS and other neuropathologies.^7^ Thus, dg-Bcan could serve as a promising tumor-specific marker for the development of novel therapeutic targeting strategies for HGG.

To this end, we have chosen to develop peptides with high affinity for dg-Bcan to deliver therapeutic payloads to HGG. Peptides are attractive tools for rationally-designed therapeutics as they are small, cost-effective, scalable, and can be easily modified and tailored to further optimize binding specificity and BBB penetration.^10^ Peptides composed of D-amino acids (a.a.) (mirror image of their native L-form) have improved protease resistance and increased serum half-life.^11^

To identify dg-Bcan-binding peptides, we screened a one-bead-one-compound (OBOC) combinatorial library built on micro-sized polystyrene beads,^12^ with each bead displaying only one unique peptide species composed of 8 D-a.a. Through a combination of high-throughput screening strategies including a magnetic-capture technique and cell-bead binding approach^13–16^ we have identified a novel dg-**B**can-**T**argeting **P**eptide (BTP), referred to as BTP-7 that can specifically bind dg-Bcan and target human glioblastoma (GBM) in an intracranial mouse xenograft model. Furthermore, BTP-7 can cross the BBB in mice and in human BBB organoids. Finally, we demonstrated the utility of BTP-7 to deliver camptothecin (CPT), a topoisomerase I inhibitor with anti-cancer properties^17^ to intracranial GBM established in mice.

## Materials and Methods

### OBOC library screening

An OBOC combinatorial library was synthesized on 1 g of 90 μm TentaGel resin (Tentagel S – NH_2_; 0.33 mmol/g loading) using a “split and pool” strategy so that each bead carried multiple copies of a unique peptide ligand.^13,16,18^ See SI for detailed screening approach.

### Cell uptake analysis

Cells were washed once with PBS and dissociated by treatment with 1 mL of 2.5 mM ethylenediaminetetraacetic acid (EDTA). After centrifugation, PBS supplemented with 10% FBS was added to create a single-cell suspension. All peptide stocks were dissolved in dimethyl sulfoxide (DMSO) and stored at −20 °C in the dark. Cells were incubated with each peptide (1–10 μM) for 3 hrs at 37 °C in the dark under constant rotation. Cells were then washed 3x with PBS/10% FBS (7000 rpm centrifugation for 3 min). Cells were resuspended in 3.7% formaldehyde and washed once with PBS. Peptide uptake (mean fluorescence intensity) was determined by flow cytometry (10,000–20,000 events) or fluorescence microscopy (n_GSCs_=10). For blocking studies, cells were incubated with a 50x molar excess of unlabeled peptide for 10 min on ice, before adding the corresponding fluorescently labeled peptide at a final concentration of 1 μM and incubated for 1.5 hrs on ice. To measure peptide internalization, cells were incubated at 37 °C or 4 °C (to inhibit endocytosis) for 3.5 hrs with Cy5.5-labeled peptide (5 μM). Cells were then washed with cold PBS/10% FBS, fixed and analyzed by flow cytometry or confocal microscopy.

### Octet binding kinetic analyses

The FortéBio OctetRed384 was used to study the binding kinetics of each peptide to recombinant human brevican (in PBS and 0.1 mM EDTA, pH 6.8). The brevican protein was deglycosylated prior to experimental use (see SI for details). All binding kinetics assays were performed using the OctetRed instrument under agitation at 1000 rpm in 0.9% NaCl irrigation with 0.05% Tween (working buffer). Assays were performed at 30°C in solid black 384-well plates (Geiger Bio-One). See SI for details.

### Synthesis of BTP-7-CPT

Disulfide-cleavable CPT prodrug was synthesized as described.^19^ The pyridyldithiol arm of the prodrug allows for conjugation to free thiols via disulfide exchange, enabling CPT to be attached to a cysteine residue on BTP-7. See SI for detailed synthesis.

#### Animal studies

All animal protocols were reviewed and approved by the in-house Institutional Animal Care and Use Committee (IACUC)

### Intracranial GBM implantation and tumor uptake analysis

100,000 GBM-6 GSC resuspended in 2μl PBS were inoculated into the striatum of female 6-8-week-old athymic mice using a stereotactic frame as described.^20^ After two weeks, tumor formation was confirmed via T2-weighted MRI. The following day, mice were injected with either Cy5.5-labeled BTP-7, BTP-8 or BTP-9 (n = 3; 100 μL of 500 μM peptide solution) via the tail vein. A ‘no peptide’ control group (Cy5.5 dye) was also included. After 8 hrs, mice were sacrificed and their brains excised, cryo-sectioned, stained with Hoechst dye and imaged by fluorescence microscopy. The mean Cy5.5 fluorescence intensity of the tumor was quantified using ImageJ and compared with the mean fluorescence intensity of the non-tumor bearing contralateral (left) side of the brain.

#### Survival study

GBM-6 intracranial tumors were established as above. At Day 24 post-implantation, T2-weighted MRI was performed, animals were randomly assigned into groups and injected intraperitoneally with either BTP-7-CPT, Scr-7-CPT or vehicle control (DMSO in 0.9% NaCl) at 10 mg/kg every other day until Day 49. At Day 47, brains were imaged by T2-weighted MRI. The Kaplan-Meier survival graph was plotted using GraphPad Prism.

### Ex vivo immunofluorescence staining of GBM tissue sections

At Day 49, one animal from each treatment group was sacrificed by CO2 asphyxiation and transcardial perfusion. Brains were frozen and cryosectioned into 16 μm sections. and immunostained for phospho-H2AX. For detailed protocol see SI.

### Statistical analysis

Data are presented as means ± SD. Significance was determined via one-way (two-way when applicable) ANOVA and Tukey’s Multiple Comparison test unless stated otherwise in the figure legends. Significance is indicated as follows: (* p < 0.05, ** p < 0.01, *** p < 0.001, **** p < 0.0001)

### Use of human specimens

All human subjects had given informed consent and signed consent forms inspected, approved by the Institutional Review Board (IRB), and provided to the PI in de-identified manner for research.

## Results

### dg-Bcan expression in primary and recurrent glioma cells and tissues

We used the BG1 polyclonal antibody (which detects a deglycosylated epitope (a.a. 535–548) in dg-Bcan) to verify that dg-Bcan was expressed exclusively in HGG specimen, as previously reported.^7,21^ Western blot analysis of dg-Bcan using BG1 as well as a “pan-brevican” antibody confirmed that dg-Bcan lacked normal brevican glycosylation (**Fig. S1a)**. We also confirmed that normally glycosylated brevican was found preferentially in the medium of cultured glioma cells, while the dg-Bcan isoform was found associated with the cell membrane (**Fig. S1a)**, as previously reported.^7^ Immunofluorescence staining and western blot analysis showed that dg-Bcan was expressed in human GBM surgical specimens but not in non-cancerous brain tissues (**Fig. 1a,b)**, consistent with previous findings^7^ and strengthening the evidence of dg-Bcan as a glioma marker. dg-Bcan expression was detected in both primary and recurrent glioma specimens. Furthermore, immunofluorescence staining of GBM biopsy samples isolated from the core and border of each patient’s tumors revealed widespread, albeit variable expression of dg-Bcan throughout the specimen (**Fig. S1b)**.

**Fig. 1.**
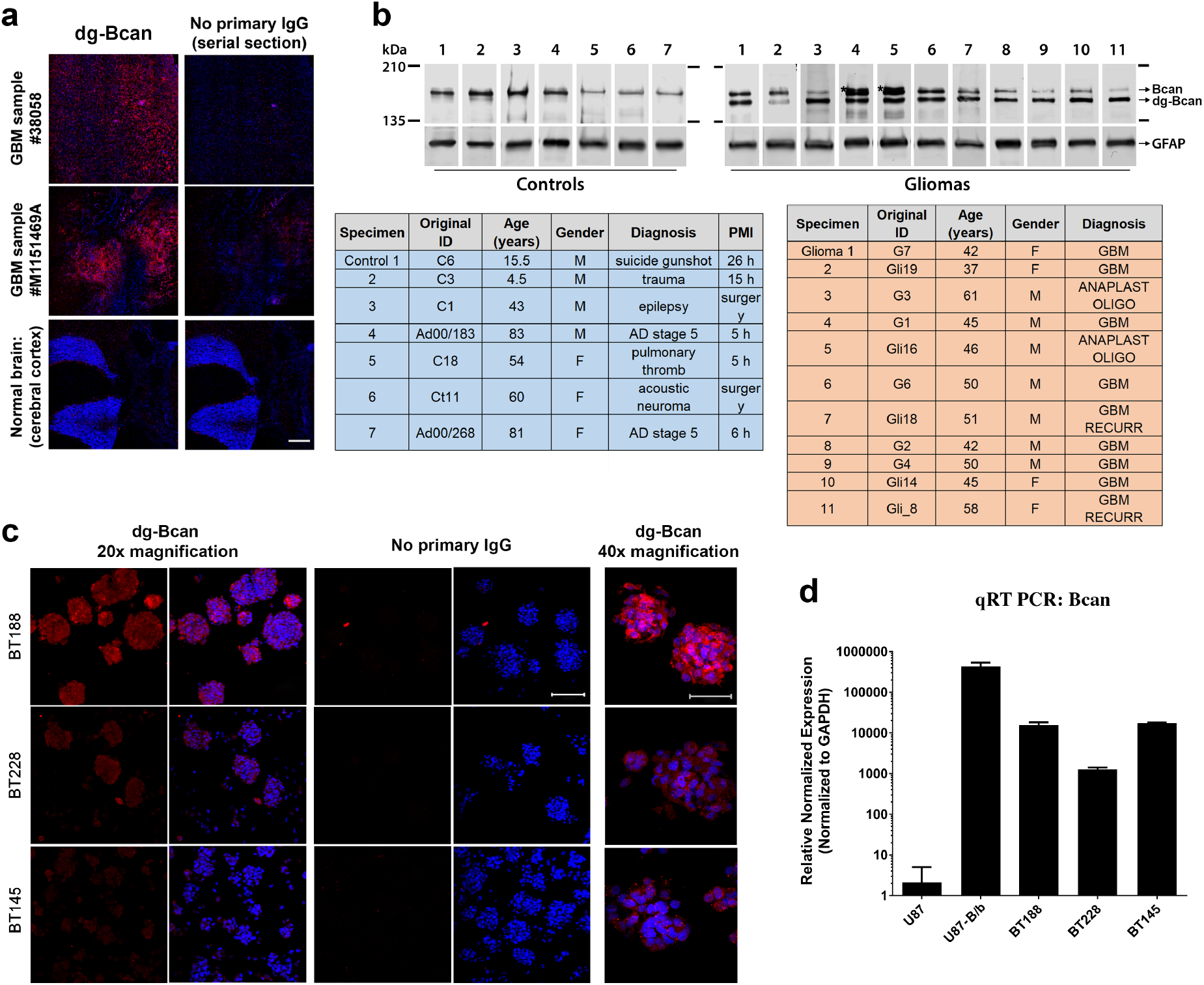
dg-Bcan is expressed in primary and recurrent glioma tissues and cells. **a)** Immunofluorescence images of frozen GBM specimens (using BG1 antibody) showing dg-Bcan (red) and nuclei (blue). Scale bar: 500 μm. **b)** Western blot of HGG (grade 3 or GBM) and control samples (using pan-brevican antibody) with corresponding clinical information (tables). PMI: Post-mortem interval for samples that were recovered after autopsy. **c)** Immunofluorescence images of patient-derived glioma stem cells (GSCs) (using BG1 antibody) showing dg-Bcan (red) and nuclei (blue). (20x scale bar 100μm, 40x scale bar 50μm) **d)** Brevican RNA level in GSCs analyzed by qRT-PCR.

Immunofluorescence staining of patient-derived glioma stem cells (GSC) revealed that dg-Bcan expression is retained in primary cultures,^22^ though expression levels varied, with BT188 showing the highest and BT145 the lowest dg-Bcan expression (**Fig. 1c)**. qRT-PCR analysis showed similar levels of brevican mRNA in BT188 and BT145 despite their different levels of dg-Bcan, suggesting that dg-Bcan levels do not necessarily correlate with brevican expression (**Fig. 1d)**. This phenomenon was also observed at the protein level in GBM tissues (**Fig. 1b)**. We confirmed the expression of dg-Bcan in other patient-derived GSC including a line derived from a recurrent tumor, GSC-401, using western blot analysis. (**Fig. S1c)**. Most established GBM cell lines, including U87 do not express endogenous brevican. Collectively, these data suggest that dg-Bcan could serve as a promising target for the development of novel therapeutic targeting strategies specific for HGG.

### OBOC peptide library screen for dg-Bcan targeting peptides

Our two-stage screening strategy was designed to identify high-affinity dg-Bcan ligands with stringency and low false positives in a high-throughput manner.^16,23^ The fully glycosylated Bcan contains chondroitin sulfate glycosaminoglycan (GAG) chains, as well as *N*-linked and *O*-linked oligosaccharides,^24^ which are not present in dg-Bcan (schematic shown in **Fig. 2a)**. We rationalized that the epitope lacking glycosylation (a.a. 535–548) in dg-Bcan is accessible for targeting for the development of specific dg-Bcan affinity peptides.

**Fig. 2.**
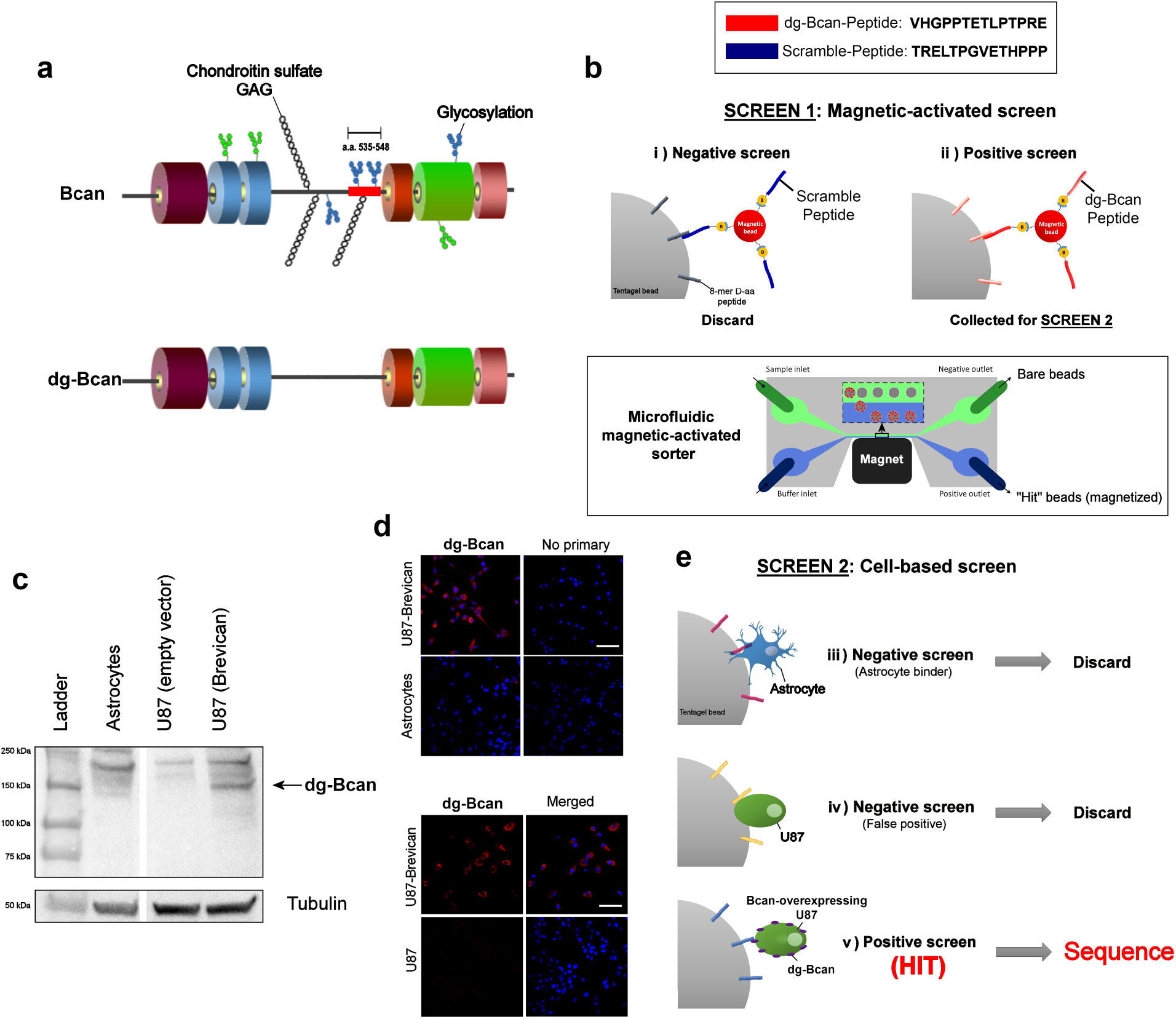
OBOC peptide library screen for dg-Bcan-binding peptides yields 7 candidates. **a)** Schematic of the brevican protein, Bcan (top) and its deglycosylated isoform, dg-Bcan (bottom). The dg-Bcan isoform lacks the vast majority of N- and O-linked carbohydrates (including chondroitin sulfate chains), thus exposing specific epitopes that are blocked in the glycosylated isoform. **b)** Schematic of magnetic-activated OBOC library screen with a microfluidic sorter. **c,d)** Western blot analysis and immunofluorescence images showing astrocytes, U87 (empty vector) and Bcan-overexpressing U87 cells. **e)** Schematic of the secondary cell-based screen.

#### Screen 1

A high-throughput magnetic capture approach was utilized, where OBOC library beads with high affinity for the target protein would be labeled with magnetic “prey” particles (**Fig. 2b)** (see SI for details). First, library beads coated with magnetic particles conjugated to the scramble(dg-Bcan)-peptide were discarded to minimize the incidence of false positives.^15^ The remaining beads that associated with magnetic particles displaying the dg-Bcan-peptide were subjected to screen 2.

#### Screen 2

To ensure the highest chance of identifying peptides that can recognize dg-Bcan in its native conformation on the cell surface, a secondary cell-based screen was conducted using dg-Bcan-overexpressing U87,^25^ as well as dg-Bcan-negative astrocytes and U87 cells (**Fig 2c,d,e**). Beads that did not associate with live astrocytes and U87 cells but had high interaction with dg-Bcan-overexpressing U87 cells were isolated. The peptides on seven hit beads were sequenced by Edman degradation

### Characterization of dg-Bcan-targeting peptides (BTP)

All peptides had a high isoelectric point (pI) and were positively charged (**Fig. 3a)**. We found that BTP-2, 5 and 10 were insoluble in aqueous solution. To characterize dg-Bcan specificity, each peptide was conjugated to fluorescein (FITC) on the C-terminus according to the peptide orientation displayed on the library bead (**Fig. 3b, Fig. S2a)**.

**Fig. 3.**
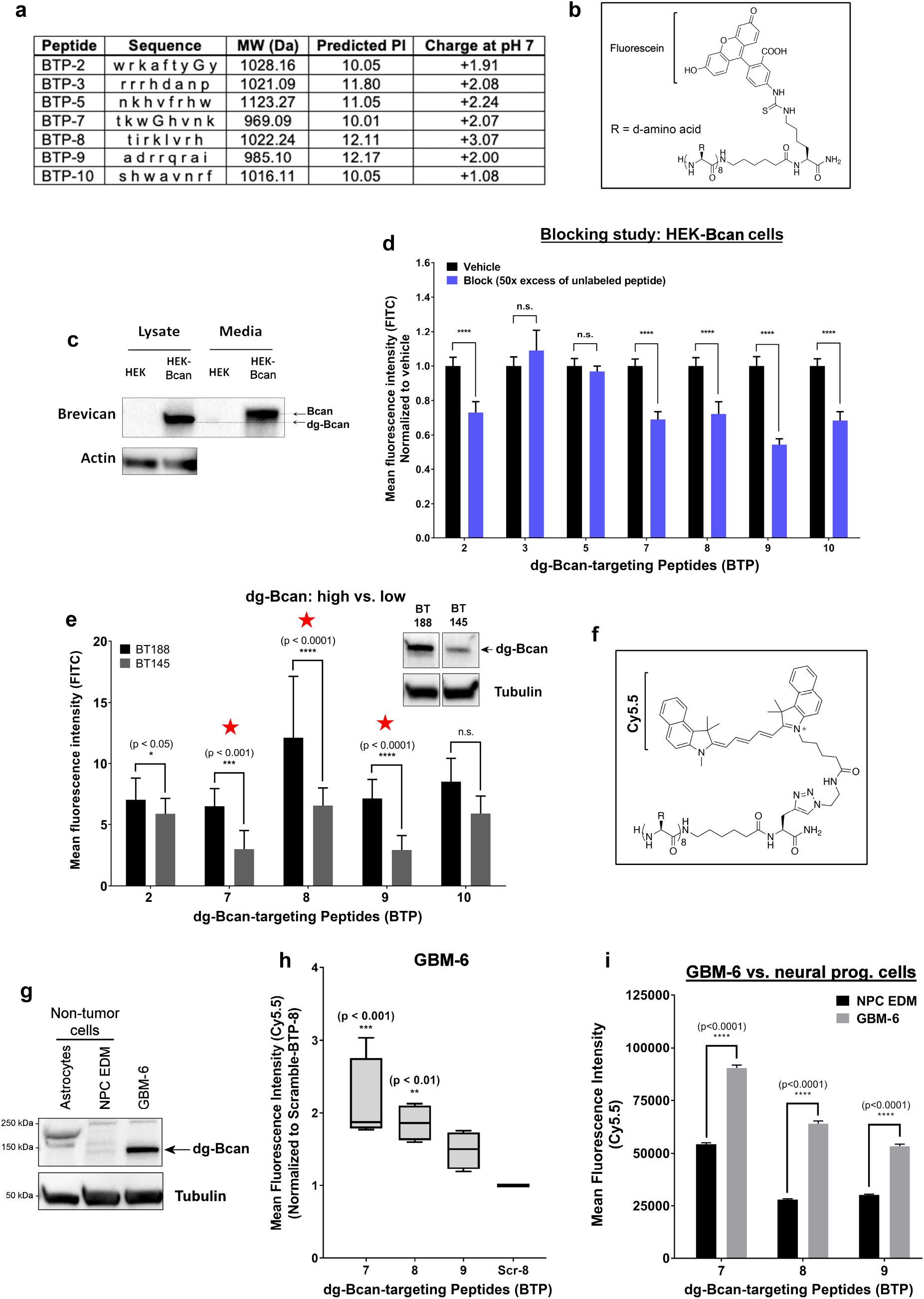
Cell uptake assessments reveal three promising dg-Bcan-targeting peptides (BTP) **a)** Table summarizing the properties of the seven peptides from the OBOC library screen. Each letter abbreviates a D-amino acid. MW = molecular weight. **b)** Chemical structure of the peptide labeled with fluorescein dye. **c)** Western blot of dg-Bcan and Bcan in human embryo kidney (HEK) cells, and Bcan-overexpressing HEK cells (HEK-Bcan). **d)** Flow cytometry analysis showing cell uptake of fluorescein-labeled peptides with or without blocking (50x excess native peptide) in HEK-Bcan cells (n_events_ = 20,000, t-test). **e)** Cell uptake analysis of fluorescein-labeled peptides in BT188 (dg-Bcan-high) or BT145 (dg-Bcan-low) glioma stem cells (GSCs) as measured by fluorescence microscopy. (n_GSCs_ = 10). **f)** Chemical structure the peptide labeled with Cy5.5 dye. **g)** Western blot analysis showing dg-Bcan expression in GBM-6 cells, astrocytes and neural progenitor cells (NPCs). **h)** Flow cytometry analysis showing GBM-6 cell uptake of the top 3 candidates, BTP-7, 8 and 9 (each group normalized to a control scramble peptide (Scr-8) (n_events_ = 20,000, significance measured in comparison to Scr-8). **i)** Flow cytometry analysis showing cell uptake of BTP-7, 8 and 9 in GBM-6 cells and NPCs (n_events_ = 20,000 cells).

The peptides were characterized using HEK293 cells that do not normally express endogenous brevican and engineered HEK293 cells with stable brevican overexpression (HEK-Bcan) (**Fig. 3c)**. Similar to primary GSCs (**Fig. S1a)**, dg-Bcan was found in cell lysates while full-length glycosylated brevican was secreted into the culture medium (**Fig. 3c)**. In a competitive binding study, FITC-labeled BTP candidates, BTP-2, 7, 8, 9 and 10 demonstrated reduced uptake in HEK-Bcan cells in the presence of 50x molar excess of unlabeled (blocking) peptide as analyzed by flow cytometry (**Fig. 3d)**, suggesting specific peptide interaction. Flow cytometry analyses confirmed that these five peptides showed increased uptake in HEK-Bcan cells compared to regular HEK cells (**Fig. S3a)**. Similarly, these peptides exhibited increased cell uptake in endogenous dg-Bcan-expressing BT-188 GSC compared to dg-Bcan-null astrocytes when analyzed by fluorescence microscopy (**Fig. S3b)**. To further narrow down the top candidates, we investigated whether these peptides had differential uptake in dg-Bcan-high BT188 cells compared with dg-Bcan-low BT145 cells (**Fig. 1c and 3e)**. Three peptides, BTP-7, 8 and 9, exhibited the highest uptake in BT188 compared with BT145 cells, and were considered the top candidates (**Fig. 3e)**.

To facilitate *in vivo* characterization, we labeled BTP-7, 8 and 9 with a Cy5.5 dye (**Fig. 3f, Fig. S2c)** that emits light in the near-infrared (NIR) region to enable deeper tissue penetration. The well-characterized GBM-6 patient-derived xenoline (PDX)^26^ was used due to its stable expression of dg-Bcan (**Fig. 3g)**, and its ability to form aggressive intracranial tumors in nude mice. Flow cytometry analysis showed that Cy5.5-labeled BTP-7 displayed the highest uptake in GBM-6 cells (**Fig. 3h**). The Cy5.5-labeled peptides exhibited greater uptake in GBM-6 cells compared with dg-Bcan-null astrocytes (**Fig. S3c,d,e**). Additionally, GBM-6 cells showed significantly higher peptide uptake than neural progenitor cells (NPC), which do not express dg-Bcan (**Fig. 3g,i**). Flow cytometry analysis also confirmed increased uptake of Cy5.5-labeled peptides in HEK-Bcan cells compared to regular HEK cells (**Fig. S3f**). Incubation of GBM-6 cells with each peptide at 4°C resulted in a dramatic reduction in peptide uptake compared to 37°C (**Fig. S3g**), suggesting ligand-receptor internalization. This was confirmed by confocal microscopy (**Fig. S3h**).

### Affinity of BTP-7 for dg-Bcan protein

To assess the binding of BTP-7, 8 and 9 to purified recombinant dg-Bcan protein, binding kinetics analyses using the Octet RED platform were performed. This revealed that native BTP-7 bound dg-Bcan protein with the highest affinity (K_D_ = 0.26 μM) compared to BTP-8 and BTP-9 (K_D_ = 1.57 μM and 1.50 μM, respectively) (**Fig. 4a,b and c**), thus making BTP-7 the top candidate for dg-Bcan targeting. These peptides did not bind the normally glycosylated form of brevican (**Fig. 4a,b and c**), highlighting their specificity for the dg-Bcan isoform. A scrambled sequence of the BTP-7 peptide (Scr-7) exhibited dg-Bcan protein binding (**Fig. 4a)**. Fluorescein-conjugated BTP-7 retained its specificity for dg-Bcan and did not bind to glycosylated brevican, although the binding affinity (K_D_ = 3.80 μM) was more than 10x weaker than the unlabeled BTP-7 (**Fig. S4a)**. Conversely, BTP-7 conjugated to a Cy5.5 dye was found to have a slightly higher affinity for dg-Bcan (K_D_ = 0.19 μM) than the native peptide (**Fig. S4b)**.

**Fig. 4.**
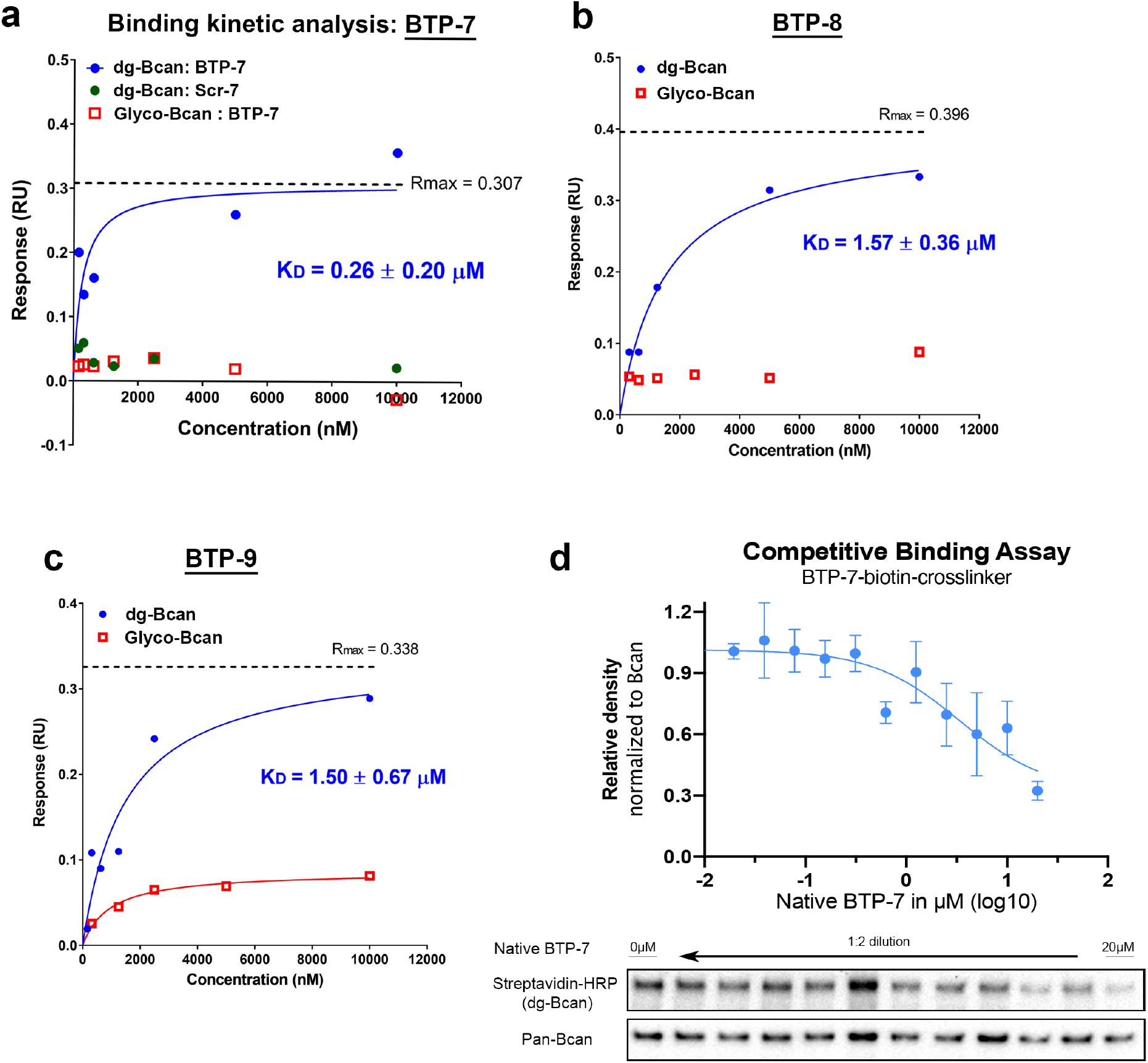
Binding affinity and specificity of top 3 peptides to recombinant dg-Bcan protein. **a,b,c)** Steady state binding kinetics analysis of BTP-7, Scr-7 (scramble BTP-7; green), BTP-8 and BTP-9 to recombinant dg-Bcan (blue) or the fully glycosylated Bcan (red) protein using the Octet (ForteBio) platform. K_D_ = dissociation constant, R_max_ = maximal response in response unit. K_D_, R_max_ and curves were calculated using a non-linear regression (one site specific binding) fit in Graphpad Prism. **d)** Competitive binding assay using biotinylated BTP-7 functionalized with a UV-crosslinker. Peptide binding/crosslinking to recombinant dg-Bcan (in the presence of native BTP-7 blocking peptide) is detected by western blot using streptavidin-HRP (n = 3).

The specificity of BTP-7 binding to dg-Bcan was further confirmed using BTP-7 functionalized with an ultraviolet (UV)-sensitive crosslinker and biotin (**Fig. S2b)**. Upon interaction, the functionalized BTP-7 can be crosslinked to dg-Bcan by exposure to UV light, and detected by western blot using streptavidin-HRP. Competitive binding studies showed that the presence of native BTP-7 (blocking) peptide decreased the level of crosslinking in a dose-dependent manner (**Fig.4d)**. This effect was not observed when blocking with the scramble peptide, Scr-7 (**Fig. S5a)**.

Next, we investigated whether BTP-7 would interact specifically with the brevican sequence at a.a. 535-548. Analysis by circular dichroism (CD) spectrometry indicated that the BTP-7 structure was altered in the presence of the dg-Bcan-peptide, and this effect was negligible in the presence of the scramble(dg-Bcan) peptide (**Fig. S5b)**. Meanwhile, the BTP-8 structure appeared to be unaffected in the presence of the dg-Bcan or scramble(dg-Bcan) peptide (**Fig. S5c)**. Increasing dg-Bcan-peptide concentration resulted in alteration of BTP-7 structure a dosedependent manner (**Fig. S5d)**. This was not observed when BTP-7 was mixed with the scramble(dg-Bcan) peptide (**Fig. S5d)**, highlighting that BTP-7 interacted with the dg-Bcan-peptide specifically.

### BTP-7 targets intracranial gliomas

We showed that BTP-7 is relatively stable in human serum, with approximately 50% of peptide detected after 12 hours at 37°C (**Fig. S6a)**. In contrast, when each residue in the BTP-7 sequence was substituted with the corresponding L-a.a., the peptide was rapidly degraded within an hour (**Fig. S6b)**.

For *in vivo* studies, patient-derived GBM-6 GSC were implanted intracranially into the right frontal lobe of nude mice and tumor formation was confirmed by MRI after two weeks. (**Fig S6c)**. Mice were then injected with either Cy5.5-labeled BTP-7, 8, 9 or Cy5.5 dye (without peptide) as a control via the tail vein. Fluorescence imaging of cryo-sections revealed that BTP-7-Cy5.5 had significantly higher tumor uptake (more than 10x) compared to other peptides 7 hours post-injection (**Fig. 5a,b)**, reaffirming BTP-7 as the prime GBM-targeting peptide. We also found that tumor tissues in the BTP-7-Cy5.5 group exhibited 4x higher fluorescence intensity compared to normal brain, underlining tumor specificity (**Fig. 5c)**. IVIS fluorescence imaging showed that BTP-7-Cy5.5 had the highest accumulation in the liver and kidneys four hours post-intravenous administration (**Fig. S6d,e**), suggesting that the major routes of peptide clearance occurred through these organs. The brain showed low peptide uptake compared to other organs (**Fig. S6e**).

**Fig. 5.**
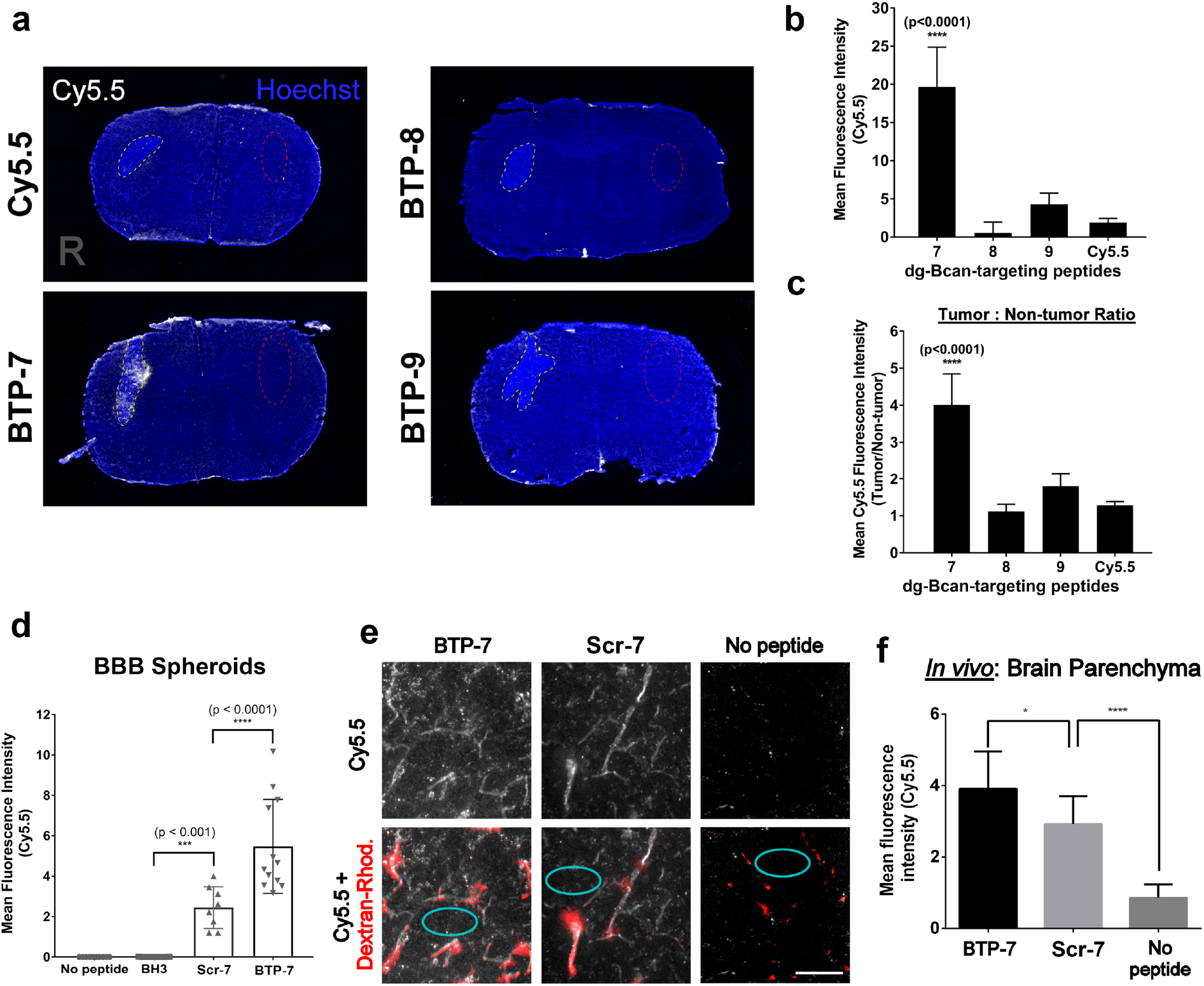
Functional characterization of BTP-7 in vitro and in vivo. **a)** Fluorescence images showing brain cryo-sections 7 hrs post intravenous injection of Cy5.5-labeled BTP-7, BTP-8 or BTP-9 (100 μL of 500 μM peptide solution, n = 3) at Day 15 post-tumor implantation. Cy5.5 dye (no peptide) was injected as a negative control. Tumor regions (right (R) brain hemisphere) are marked with yellow dotted lines. **b)** Quantification of the mean Cy5.5 intensity in the tumor (n_sections_ = 15). **c)** Quantification of tumor:non-tumor ratio (from a); non-tumor regions (left hemisphere) are marked with red dotted lines. Significance was compared with the Cy5.5 control group. **d)** Permeability of Cy5.5-BTP-7 and Scr-7 in BBB organoids (BH3 peptide used as a non-permeable control) (n_spheres_ = 4-6, c_peptide_ = 10 μM, t = 3 hrs). **e)** Fluorescence images of brain cryo-sections (30 μm slices) showing the distribution of Cy5.5-BTP-7 and Cy5.5-Scr-7 (white) in the frontal lobe of non-tumor bearing mice upon intravenous administration. TRITC-dextran (155 kDa; red) was injected 4 hrs later intravenously, and the mice were sacrificed 30 min later. Red areas indicate regions of high perfusion. Scale bar: 100 microns. **f)** BBB permeability was quantified by measuring the mean Cy5.5 intensity within the brain parenchyma outside of highly perfused areas (cyan circles in panel **e)** (n_tissues_ = 2; n_areas_ = 10).

### BBB penetration

We examined whether BTP-7 could cross the BBB using an *in vitro* human BBB organoid model.^27^ BTP-7-Cy5.5 exhibited a dramatic influx into the organoids, highlighting potential BBB-penetrating properties (**Fig. 5d)**. An increased influx was also observed in the organoids with the scramble variant, Scr-7-Cy5.5 (compared to the control non-penetrating BH3 peptide^27^), though the level of penetration was significantly lower than that observed with BTP-7 (**Fig. 5d)**. In the presence of BTP-7-Cy5.5, the organoids remained relatively impermeable to high molecular weight dextran molecules (4.4 kDa) (**Fig. S7**), showing that BTP-7 did not damage the BBB spheroid barrier. To confirm this observation *in vivo*, BTP-7-Cy5.5 or Scr-7-Cy5.5 was administered intravenously into healthy non-tumor bearing mice. After 4 hours, TRITC-dextran (155 kDa) was administered intravenously so that perfused blood vessels could be visualized. Confocal microscopy revealed that both BTP-7 and Scr-7 accumulated within the brain parenchyma (outside of regions labeled by TRITC-dextran), though the level of BTP-7 was higher than that of Scr-7 (**Fig. 5e,f**). Overall, these results demonstrated the ability of BTP-7 to cross the BBB *in vivo* and in human BBB organoids.

### BTP-7 as a vehicle to deliver toxic payload to HGG

GBM-6 proliferation and invasion in collagen was not affected in the presence of BTP-7 **(Fig. S8a,b,c,d)**. We hypothesized that BTP-7 could serve as a vehicle to target a therapeutic payload directly to tumor tissues. As a proof-of-principle, we conjugated camptothecin (CPT) to BTP-7 (BTP-7-CPT). CPT is a potent topoisomerase I inhibitor, and has shown promising results in preclinical models but failed to translate well in the clinic in large part due to its insolubility.^17,28^ Here, we functionalized BTP-7 with CPT (BTP-7-CPT) via a disulfide linker at the C-terminus that is cleaved upon cell internalization to release the active CPT drug metabolite (**Fig. S9a,b)**.^29^ A scrambled peptide-drug conjugate, Scr-7-CPT was synthesized as a control. Binding kinetic analysis using the Octet platform showed that BTP-7-CPT binds recombinant dg-Bcan protein with a K_D_ of 10.25 μM (**Fig. S9c)**, demonstrating that CPT conjugation reduced the peptide’s affinity for dg-Bcan (K_D_ of BTP-7 = 0.26 μM) (**Fig. 4a)**. BTP-7-CPT had a relatively short halflife and was no longer detectable in human serum within 2 hours (**Fig. S9d)**. Nonetheless, a cell proliferation assay showed that BTP-7-CPT was more toxic to dg-Bcan**-**overexpressing HEK cells than regular HEK cells (**Fig. 6a)**. This difference was not seen when the cells were treated with Scr-7-CPT (**Fig. 6b)**. BTP-7-CPT was also cytotoxic to GBM-6 cells in a dose-dependent manner with an IC_50_ of 4.98 μM (**Fig. 6c, Fig. S9e**).

**Fig. 6.**
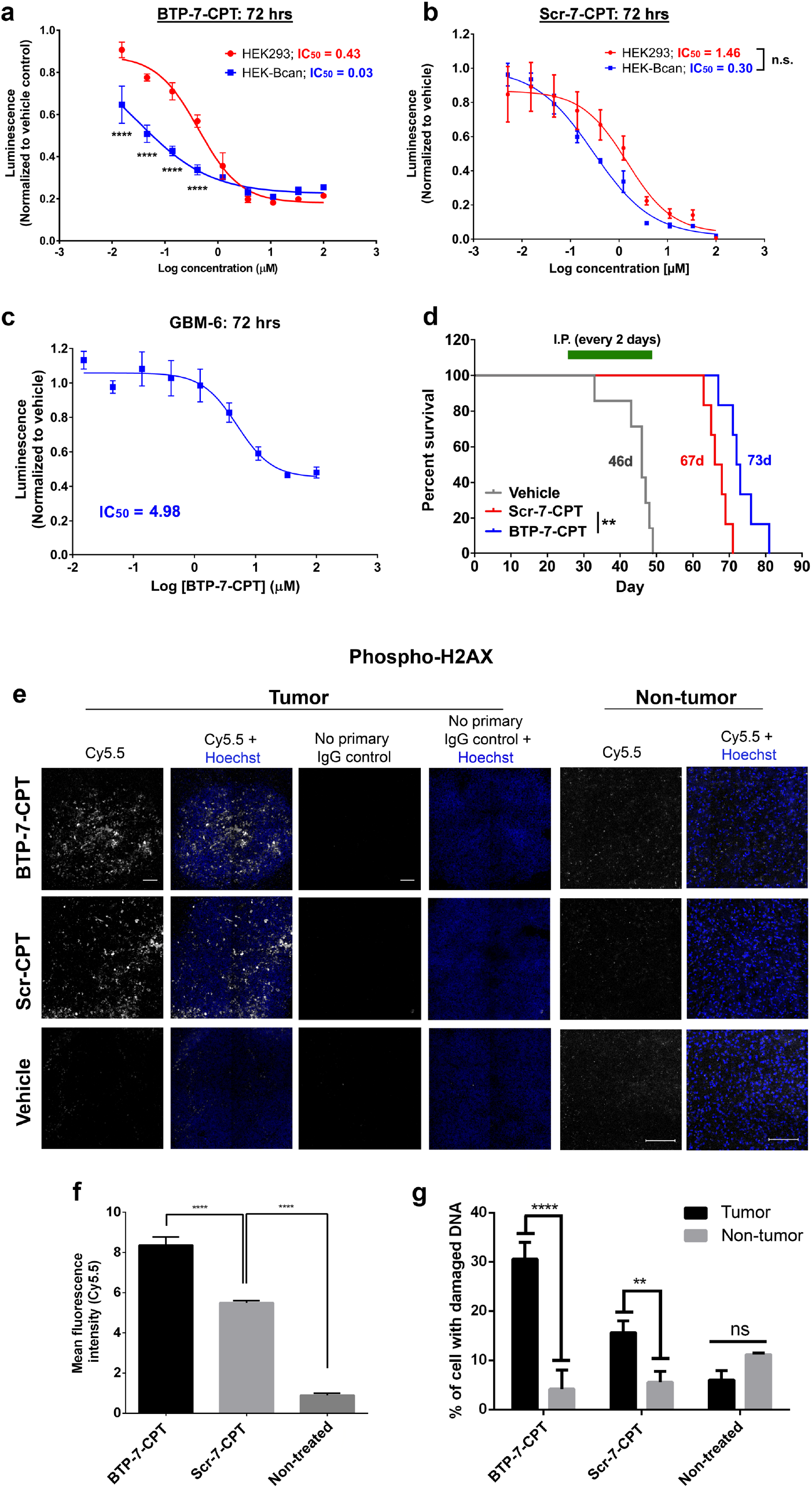
Camptothecin (CPT) functionalized with BTP-7 preferentially inhibits proliferation of Bcan-expressing cells and prolongs survival in mouse model of GBM. **a,b)** Luminescent cell viability (CellTitre Glo) assay of Bcan-overexpressing HEK (HEK-Bcan) cells (blue) and control HEK cells (red) in the presence of BTP-7-CPT or Scr-7-CPT after 72 hrs. **c)** CellTitre Glo assay of GBM-6 cells treated with BTP-7-CPT (n_wells_ = 3). IC_50_ values were measured through the non-linear ‘log(inhibitor) vs. response - variable slope (four parameters)’ fit. **d)** Kaplan-Meier survival plot of mice bearing GBM-6 tumors. Mice were injected intraperitoneally (10 mg/kg) with BTP-7-CPT (blue; median survival = 73 days), Scr-7-CPT (red; median survival = 67 days) or vehicle (gray; median survival = 46 days). Treatment was performed every 2 days starting from Day 25 to Day 49 post tumor implantation. A significant difference (p < 0.01) is observed between the BTP-7-CPT and Scr-7-CPT group, and between the treated groups and vehicle group, as determined by the Log-rank (Mantel-Cox) test. **e)** Immunofluorescence staining of tissue cryo-sections for phospho-H2AX from mice brains harvested at day 49 after 13 treatments (nuclei counterstained with Hoechst dye (blue)). Scale bar: 100 microns. **f)** Quantification of phospho-H2AX signal within the tumor area. (n_tissue_ = 3). **g)** Quantification of number of nuclei with positive phospho-H2AX signal in tumor and nontumor tissues (n_tissue_ = 3).

To determine potential toxicity, healthy nude mice were treated with a single intraperitoneal dose of 25, 50 or 100mg/kg of BTP-7-CPT. The animals did not exhibit any signs of morbidity or weight loss over 14 days, demonstrating the safety of BTP-7-CPT in mice even at high doses. To investigate the efficacy of BTP-7-CPT in an intracranial GBM mouse model, GBM-6 tumors were established and tumor formation was confirmed via MRI (**Fig. S9f).** Mice were randomized into three treatment groups: vehicle (control), Scr-7-CPT, and BTP-7-CPT. We observed a substantial decrease in tumor size in mice treated with BTP-7-CPT or Scr-7-CPT in comparison to vehicle (**Fig. S9f)**. While treatment with either of the CPT conjugates extended survival compared to the control animals, mice treated with BTP-7-CPT had a significant survival benefit over mice treated with Scr-7-CPT (median survival = 73 vs. 67 days, p < 0.01) (**Fig. 6d)**. We observed similar results in a second survival study with a prolonged treatment regimen (25 vs. 50 days) (median survival = 82 vs. 88 days, p < 0.05) (**Fig. S9g).**

*Ex vivo* analysis of brain cryo-sections showed that tumor tissues in both treatment groups displayed significantly higher phospho-H2AX levels, indicating greater DNA damage compared to controls (**Fig. 6e,f**). Tumor tissues from the BTP-7-CPT group had significantly higher level of phospho-H2AX than the Scr-7-CPT group, highlighting the ability of BTP-7 to improve drug targeting to GBM (**Fig. 6e,f,g**). Furthermore, for both treated groups, there was no evidence of DNA damage in non-cancerous brain tissues (**Fig. 6e,g**). Overall, these results show that BTP-7-CPT preferentially targeted dg-Bcan-expressing gliomas without harming healthy brain tissues.

## Discussion

Tumor-specific targeting agents represent an attractive approach to developing precision therapeutics. To overcome tumor genetic heterogeneity,^30^ we have focused our efforts on an ECM protein, brevican, that has a unique glioma-specific isoform, dg-Bcan. The tumor specificity and ubiquity of dg-Bcan underscore its potential as a novel glioma-specific marker for the development of targeted therapies.

Brevican is a ECM proteoglycan that is highly upregulated in HGG and implicated in tumor invasion and progression.^25,31^ It has multiple isoforms produced by glycosylation, cleavage, and alternative splicing but only one of these isoforms, dg-Bcan is uniquely expressed in human gliomas.^32^ Brevican activates EGFR/mitogen-activated protein kinase (MAPK) signaling and fibronectin production in glioma cells, increasing their migratory and invasive properties.^33^ The specific function of dg-Bcan remains unknown, although it provides an excellent and accessible target that is glioma-specific. Primary patient-derived GSC in their undifferentiated state retain dg-Bcan expression, even after being passaged in mice, and can be used for both *in vitro* and *in vivo* GBM modeling and drug targeting analyses.

Here, we have performed a stringent peptide library screen to discover BTP-7, a novel D-a.a. peptide that is stable in serum, binds recombinant human dg-Bcan with nanomolar affinity with little to no cross-reactivity with the major full-length glycosylated brevican isoform, labels cultured cells in a dg-Bcan-dependent manner and homes to human GBM xenografts established intracranially in mice. To demonstrate therapeutic targeting, we have conjugated BTP-7 to CPT, which is known to have limited clinical utilization due to its poor solubility.^34^ For our animal studies, we chose a treatment dose of 10 mg/kg to ensure minimal risk of toxicity over continuous administrations. We observed that BTP-7 enables *in vivo* drug targeting to the tumor, leading to increased tumor cell death and prolonged animal survival.

BTP-7 is the first dg-Bcan-binding peptide described, and we have demonstrated its ability to target HGG. In the future, we will perform rational peptide modifications to develop second-generation peptides with increased dg-Bcan affinity and BBB permeability, investigate stable cleavable linkers such as acid or enzyme-cleavable linkers^35^ and explore BTP-7 conjugation to a wide array of other anti-cancer therapeutics to further promote therapeutic efficacy. Additionally, BTP-7 could provide a robust platform for the development of novel cytotoxic radionuclides for targeted radiotherapy, as well as targeted non-invasive molecular imaging agents, such as BTP-7-conjugated radiotracers (for positron emission tomography (PET) imaging) and gadolinium or iron oxide nanoparticles (for MRI).

Taken together, this work shows that targeting dg-Bcan is a novel approach with potential translatability for GBM. This study opens the door for further development of BTP-7 and testing in combination with standard of care.

## Supplementary Materials

### Supplementary Materials and Methods

*Fig S1. Expression of dg-Bcan in high-grade glioma tumor samples and glioma stem cell lines*

*Fig. S2. Workflow for the synthesis of labeled peptides*

*Fig. S3. Characterization of peptides for dg-Bcan specificity using cell uptake studies*

*Fig. S4. Binding affinity of functionalized BTP-7 to dg-Bcan protein*

*Fig. S5. Binding of BTP-7 to dg-Bcan and the dg-Bcan-derived peptide*

*Fig. S6. In vivo analysis of BTP-7*

*Fig. S7. BBB permeability of BTP-7 in vitro*

*Fig. S8. No effect of BTP-7 on GBM cell proliferation and invasion*

Fig. S9. Characteristics and efficacy of Camptothecin (CPT) functionalized with BTP-7 in vitro and in vivo

*Supplementary Notes: Sequences, synthesis and chromatograms*

