## Supplementary Info for "BTP-7, a novel peptide for therapeutic targeting of malignant brain tumors": SI von Spreckelsen et al_Neuro_onc_main.pdf

**Affiliations:**

#### Supplementary Materials

**Supplementary Materials and Methods**

**Materials:**

For peptide synthesis, N$^{\alpha}$-Fmoc protected D-amino acids (a.a., Fmoc-D-Ala-OH, Fmoc-D-Arg(Pbf)-OH; Fmoc-D-Asn(Trt)-OH; Fmoc-D-Asp-(O*t*-Bu)-OH; Fmoc-D-Cys(Trt)-OH; Fmoc-D-Gln(Trt)-OH; Fmoc-D-Glu(O*t*-Bu)-OH; Fmoc-D-His(Boc)-OH; Fmoc-D-Ile-OH; Fmoc-D-Leu-OH; Fmoc-D-Lys(Boc)-OH; Fmoc-D-Lys(alloc)-OH, Fmoc-D-Met-OH; Fmoc-D-Phe-OH; Fmoc-D-Pro-OH; Fmoc-D-Ser(*t*-But)-OH; Fmoc-D-Thr(*t*-Bu)-OH; Fmoc-D-Trp(Boc)-OH; Fmoc-D-Tyr(*t*-Bu)-OH; Fmoc-D-Val-OH) and Fmox-L-Lys(biotin)-OH were purchased through Advanced ChemTech (Louisville, KY). Chem-Impex (Wood Dale, IL) and Peptides International (Louisville, KY). N$^{\alpha}$-Fmoc protected L-amino acids (Fmoc-L-Ala-OH, Fmoc-L-Arg(Pbf)-OH; Fmoc-L-Asn(Trt)-OH; Fmoc-L-Asp-(O*t*-Bu)-OH; Fmoc-L-Cys(Trt)-OH; Fmoc-L-Gln(Trt)-OH; Fmoc-L-Glu(O*t*-Bu)-OH; Fmoc-Gly-OH, Fmoc-L-Ile-OH; Fmoc-L-Leu-OH; Fmoc-L-Lys(Boc)-OH; Fmoc-L-Met-OH; Fmoc-L-Phe-OH; Fmoc-L-Pro-OH; Fmoc-L-Ser(*t*-But)-OH; Fmoc-L-Thr(*t*-Bu)-OH; Fmoc-L-Trp(Boc)-OH; Fmoc-L-Tyr(*t*-Bu)-OH; Fmoc-L-Val-OH;) were purchased from the Novabiochem-line through Millipore Sigma (Darmstadt, Germany). Fmoc-L-His(Boc)-OH was bought from CEM. Succinimidyl 4,4'-azipentanoate (NHS-Diazirine) was purchased from Thermo Fisher Scientific (Waltham, MA) (#26167). Amino acids in peptide sequences are abbreviated with one letter code; capitalized letters refer to L-amino acids, lowercase letters refer to D-amino acids.

H-Rink Amide-ChemMatrix resin was obtained from PCAS BioMatrix Inc. (St-Jean-sur-Richelieu, Quebec, Canada). 4-pentynoic acid, Fmoc-L-propargylglycine, 2-(1H-benzotriazol-1-yl)-1,1,3,3-tetramethyluronium hexafluorophosphate (HBTU), and 2-(7-aza-1H-benzotriazole-1-yl)-1,1,3,3-tetramethyluronium hexafluorophosphate (HATU) were purchased from Chem Impex (Wood Dale, IL) and P3 Biosystems (Louisville, KY). (7-Azabenzotriazol-1-

yloxy)tripyrrolidinophosphonium hexafluorophosphate (PyAOP) was purchased from P3 Biosystems (Louisville, KY). AldraAmine trapping agents (for 1000–4000 mL DMF), N-methyl pyrrolidinone (NMP), triisopropylsilane (TIPS), *t*-butylmethyl ether (TBME), Diisopropylethylamine (DIEA; 99.5%, biotech grade), piperidine ( ACS reagent, ≥99.0%), trifluoroacetic acid (HPLC grade, ≥99.0%), triisopropylsilane (≥98.0%), formic acid (FA, ≥95.0%), dimethyl sulfoxide (DMSO, HPLC grade, ≥99.7%) were purchased from Sigma-Aldrich. N,N-Dimethylformamide (DMF), dichloromethane (DCM), and HPLC-grade acetonitrile were from EMD Millipore (Billerica, MA). All solvents used for HPLC-MS were purchased from EMD and Fluka (Darmstadt, Germany). Cy5.5-azide was obtained from Lumiprobe (Hallandale Beach, FL). All other chemicals and reagents were purchased from Sigma-Aldrich (St. Louis, MO). Water was deionized using a Milli-Q Reference water purification system (EMD Millipore, Billerica, MA). Nylon 0.22 µm syringe filters were TISCH brand SPEC17984.

The following supplies and reagents were used in our experimental studies: Recombinant human brevican (R&D Systems Minneapolis, MN; Cat. # 4009-BC-050), streptavidin-coated pink-fluorescent magnetic particles (2.0–2.9 µm, Spherotech, Lake Forest, IL; Cat. # FSVM-2058-2), *O*-glycosidase (Roche; 11347101001), Neuraminidase (Sialidase) (Sigma-Aldrich, St. Louis, MO); Cat. # 10269611001), PNGase F (New England Biolabs, Ipswich, MA; Cat. # P0704S), Octet Ni-NTA Biosensors (Forte Bio, Fremont, CA; Cat. # 18-5101), CellTiter-Glo 3D cell viability assay (Promega, Madison, WI; Cat. # G983), and PureCol purified bovine collagen solution (Advanced Biomatrix, San Diego, CA; Cat. # 5005-B). The dg-Bcan-peptide and scramble(dg-Bcan)-peptide with and without functionalization with biotin-Ahx (Ahx: aminohexanoic acid) on the N-terminus were synthesized by GenScript USA Inc.,Piscataway, NJ. The OBOC library and the FITC-labeled peptides were synthesized in the Luyt laboratory. All other peptides (including the native peptides, Cy5.5-labeled and peptide-drug conjugates) were synthesized in the Pentelute lab using established protocols.[1]

**Instruments:**

For fluorescence imaging, the Zeiss LSM 710 laser scanning confocal microscope or the Nikon Eclipse Ti epi-fluorescence microscope equipped with a QIClick camera was used. For wide-field time-lapse imaging for cell invasion/migration assays, the Nikon Eclipse TE2000-U epi-fluorescence microscope was used. For flow cytometry analysis, either the Becton Dickinson SORP LSR II or Beckman Coulter Gallios (from the Beth Israel Deaconess Medical Center Flow Cytometry Core) was used. For binding kinetic analyses, the Octet RED384 Platform (Forte Bio) and the Jasco J-815 Circular Dichroism (CD) Spectropolarimeter (from the Harvard University Center for Molecular Interactions) were used. For UV crosslinking, we used the Fisherbrand UV crosslinker. For magnetic resonance imaging (MRI), we used a high-field (7 Tesla) unit MR Biospec 70/20 + cryoprobe (Bruker) for small animal imaging at the Brigham and Women's Hospital animal facility. Fluorescence imaging of whole organs was conducted using the *In Vivo* Imaging System (IVIS) Lumina III (Perkin Elmer) connected to isoflurane vaporizer.

**Liquid chromatography–mass spectrometry (LC-MS)**

For mass spectrometry analysis, the filtered peptide solution (10 µL of a 1mg/mL solution) was diluted in 50% acetonitrile in water with 0.1% TFA (90 µL) to a final concentration of approximately 0.1 mg/mL. LC-MS chromatograms and associated high resolution mass spectra were acquired using an Agilent 6520 Accurate-Mass Q-TOF LCMS (abbreviated as 6520) or an Agilent 6550 iFunnel Q-TOF LCMS system (abbreviated as 6550). Solvent compositions used in the LC-MS are water with 0.1% formic acid additive (solvent A) and acetonitrile with 0.1% formic acid additive (solvent B).

**Mass-directed reversed-phase high performance liquid chromatography (RP-HPLC)**

For RP-HPLC purification, the crude lyophilized peptides were dissolved in water with 0.1% TFA additive containing a minimal amount of acetonitrile for solubility (e.g. 5% acetonitrile). All samples were filtrated through a Nylon 0.22 µm syringe filter prior to purification. For all HPLC purifications, a gradient of acetonitrile with 0.1 % TFA additive (solvent B) and water with a 0.1% TFA additive (solvent A) was used unless otherwise noted. Specific purification conditions such as column and gradient are specified for each case.

**NMR Spectroscopy.** Proton nuclear magnetic resonance ($^{1}$H NMR) spectra were recorded in 5 mm tubes on a Bruker Avance Neo spectrometer in deuterated solvents at room temperature. Chemical shifts (□ scale) are expressed in parts per million (ppm) and are calibrated using residual protic solvent as an internal reference (DMSO: □ = 2.50 ppm). Data for $^{1}$H NMR spectra are reported as follows: chemical shift (□ ppm) (multiplicity, coupling constants (Hz), integration). Couplings are expressed as: s = singlet, d = doublet, t = triplet, q = quartet, m = multiplet or combinations thereof. Carbon chemical shifts (□ scale) are also expressed in parts per million (ppm) and are referenced to the central carbon resonance of the solvent (DMSO: □ = 39.52 ppm). In order to assign the $^{1}$H and $^{13}$C NMR spectra, a range of 2D NMR experiments (COSY, HSQC, HMBC, NOESY) were used as appropriate.

**Infrared spectroscopy (IR).** Infrared spectra (IR) were recorded on a Bruker Alpha II FTIR. IR data is reported in frequency of absorption ($cm^{-1}$). The IR bands are characterized as: w = weak, m = medium, s = strong, br = broad, or combinations thereof.

**Human malignant glioma tissue specimens**

Biopsy or surgical specimens were obtained as frozen tissues (stored at -80°C) from BWH Neuropathology, Erasmus Medical Center (Rotterdam, Netherlands), and Upstate Medical Center, State University of New York (SUNY). All pathology reports were provided in a de-identified manner, and details are included in this article.

**Western blotting**

Western blotting was performed as previously described.(27) Two rabbit polyclonal antibodies were produced against synthetic peptides corresponding to the chondroitin sulfate attachment region of brevican: The pan-brevican (Bcan) antibody (targeting a.a. 506-529 of rat brevican) detected both glycosylated and deglycosylated isoforms of the protein, whereas the antibody BG1 (targeting a.a. 537–548 of human brevican) specifically recognized the deglycosylated isoform (verified in **Fig. S1).** Horseradish peroxidase (HRP)-conjugated anti-rabbit IgG (GE Healthcare; Cat. # NA934V) was used as a secondary antibody, and SuperSignal West Femto Maximum Sensitivity chemiluminescent substrate (Thermo Fisher Scientific, Waltham, MA; Cat. # 34096) was used for enhanced chemiluminescence (ECL) detection.

**Immunofluorescence staining**

The following antibodies were used for immunofluorescence staining: BG1 (see above) and rabbit anti-phospho-H2AX (Cell Signaling Technology, Danvers, MA; Cat. # 9718). Anti-rabbit Alexa-Fluor 647 secondary antibody (Invitrogen, Carlsbad, CA, Carlsbad, CA) and Hoechst dye (Life Technologies, Carlsbad, CA; Cat. # H3570) were used for detection.

*Human tissue*

The slides were fixed with 3.7% formaldehyde for 10 min at room temperature. Tissues were permeabilized with phosphate buffered saline (PBS) containing 0.1% Tween-20 (v/v) for 30 min. Blocking was performed in 5% normal chicken serum diluted in PBS containing 0.025% Tween-20 (v/v) for 1 hr at room temperature. Then, the tissues were incubated with the BG1 anti-dg-Bcan antibody (1:100 dilution) overnight at 4 $^{o}$C. Tissues were washed three times with PBS containing 0.025% Tween-20 in a Coplin jar, and then incubated with anti-rabbit Alexa Fluor 633 antibody (Invitrogen, Carlsbad, CA) (1:1000 dilution) and Hoechst dye (1:1000 dilution) in blocking solution for 1 hr at room temperature. Tissues were then washed three times with PBS containing 0.025% Tween-20. Mounting media was applied onto each tissue before a coverslip was mounted onto the slide. To prevent dehydration of tissues for long-term storage, all edges of the coverslip were sealed onto the tissue-containing slide using clear nail polish.

*Xenograft mouse brain/tumor tissue*

Animals were sacrificed by $CO_2$ asphyxiation. Transcardial perfusion was performed using 50 mL PBS, followed by 50 mL of formalin. The brains were excised, submerged in 15% sucrose, followed by 30% sucrose (overnight at 4$^{o}$C) under constant rotation. The brains were frozen and cryosectioned into 16 μm sections.

Brain tissue sections were fixed with methanol for 1 min. The slides were then washed twice with PBS + 0.025% Triton X-100. Tissue sections were blocked with blocking solution (10% normal goat serum diluted in PBS + 0.025% Triton X-100). Primary antibody against phospho-H2AX was added to the blocking solution (at 1:100 dilution) and the tissues were incubated overnight at 4 $^{o}$C in a dark humidified box. The slides were then washed three times (2 min each) with PBS + 0.025% Triton X-100. Tissues were then incubated with Alexa Fluor secondary antibody and Hoechst 33342 (both at 1:1000 dilution) in blocking solution. Then, the slides were washed three times (2 min each) with PBS + 0.025% Triton X-100. Vectashield mounting medium for fluorescence were applied onto the tissues, and a coverslip was mounted onto the slide. The edges of the coverslip were sealed with clear nail polish, and the tissues were imaged using an epifluorescence microscope under a 20x objective.

**Cell lines and culture conditions**

All glioma stem cell lines and patient-derived glioma stem cells (obtained from BWH Pathology or Erasmus Medical Center) were cultured as neurospheres in Neurobasal medium (Invitrogen, Carlsbad, CA) supplemented with 2% B27 (Invitrogen, Carlsbad, CA), 1% glutamine (Invitrogen, Carlsbad, CA), epidermal growth factor (EGF) (20 ng/ml; PeproTech, Rocky Hill, NJ), and fibroblast growth factor-2 (FGF) (20 ng/ml; PeproTech, Rocky Hill, NJ). The GBM-6 cells used in our studies had been passaged once in a mouse brain, where the cells were implanted intracranially and allowed to form a solid tumor over 30 days. Then, the GBM-6 tumor was excised, dissociated in a tissue culture flask and cultured in Neurobasal medium as above. Human neural progenitor cells (NPC) were purchased from EMD Millipore, Billerica, MA and cultured in neurobasal media. U87 and human embryonic kidney (HEK) cells were cultured in DMEM (Invitrogen, Carlsbad, CA) supplemented with 10% FBS. Cells were grown in T25 or T75 vented-cap tissue culture flasks (Sarstedt AG and Co). HEK-Bcan cells were generated by cloning hBCAN cDNA into the pcDNA3.1 vector and stably transfected in HEK293 cells as previously described.(10) Primary human astrocytes were purchased from Lonza Bioscience, Basel, Switzerland and cultured in Astrocyte Growth Medium (AGM; Lonza Bioscience, Basel, Switzerland) (consisting of astrocyte basal medium supplemented with hEGF (human epidermal growth factor), insulin, ascorbic acid, GA-1000 (Gentamicin, Amphotericin-B), L-glutamine and 1% fetal bovine serum (FBS)). Astrocytes were grown in T75 Cell+ vented-cap tissue culture flasks (Sarstedt AG and Co, Nümbrecht Germany). Human brain microvascular pericytes (HBVP) (ScienCell Research Laboratories, Carlsbad, CA) were cultured in Pericyte Medium (ScienCell Research Laboratories, Carlsbad, CA) containing 2% FBS, pericyte growth supplement and penicillin-streptomycin. Immortalized human cerebral microvascular endothelial cells (hCMEC/D3) (Cedarlane Laboratories, Burlington, Canada) were maintained in culture in Endothelial Growth Medium (EGM-2) containing hEGF, hydrocortisone, GA-1000, FBS, VEGF, hFGF-B, $R^3$-IGF-1, ascorbic acid and heparin (Lonza Bioscience, Basel, Switzerland). For BBB organoid formation in low-attachment co-culture condition and functional assays, the spheroids were maintained in EGM-2 (Lonza Bioscience, Basel, Switzerland) supplemented with 2% human serum (Valley Biomedical, Winchester, VA; Cat. # HS1021) and with the elimination of VEGF supplementation (this media formulation will henceforth be known as 'BBB working media'). All cells were cultured in a humidified incubator at 37 $^oC$ with 5% $CO_2$, and 95% natural air. For cell dissociation, StemPro Accutase Cell Dissociation Reagent (Thermo Fisher Scientific, Waltham, MA; Cat. # A1110501) was used on glioma stem cells cultured as neurospheres, and Trypsin-EDTA (0.05% v/v, with phenol red) (Thermo Fisher Scientific, Waltham, MA; Cat. # 25300054) was used for detaching adherent cells. All cell lines were regularly tested for mycoplasma contamination.

**Evaluating brevican expression by qRT-PCR**

Glioma stem cells were washed once with PBS by centrifugation at 300 rcf for 5 min. RNA was extracted by resuspending the cells in 100 µL of Trizol reagent in a 1.5 mL microcentrifuge tube. After 5 min at room temperature, 20 µL of chloroform was added and vortexed rigorously for 15 s. After 2-3 mins of incubation at room temperature, the tube was centrifuged at 12,000 rcf for 15 min at 4 °C. The upper aqueous phase (colorless) was collected using a micropipette and placed inside a new microcentrifuge tube. Then, 50 µL of isopropanol was added to precipitate the RNA. After incubating for 10 min at room temperature, the tube was centrifuged at 12,000 rcf for 10 min at 4 °C. The supernatant was removed, and the RNA was washed once with 500 µL of 70% ethanol. The tube was centrifuged again at 12,000 rcf for 5 min at 4 °C. The ethanol was removed, and the pellet allowed to air dry for 10 min. Once dried, the RNA was resuspended in 10 µL of RNA-ase free $H_2O$, and the concentration of the RNA solution was determined using the NanoDrop spectrophotometer (Thermo Fisher Scientific, Waltham, MA). RNA was reverse-transcribed using iScript cDNA Synthesis Kit (BioRad, Hercules, CA) and quantitative real-time PCR was performed using SYBR Green Master Mix (Applied Biosystem, Foster City, CA) qRT-PCR was performed using the following primer sequences:

| | | |
|---|---|---|
| BCAN Forward | – | 5'-GCTCCTGCAGCTTTAGCAG-3', |
| BCAN Reverse | – | 5'-AGGTAGTGGACGTGGCAAG-3' |
| GAPDH Forward | – | 5'-AGTCCATGCCATCACTGCCAC-3' |
| GAPDH Reverse | – | 5'-ATGACCTTGCCCACAGCCTTG-3' |

The following conditions were used: 70 °C for 3 min, 4 °C for 5 min, 42 °C for 1 hr, 70 °C for 10 min, and then stored at 4 °C. All datasets were normalized to the corresponding GAPDH values.

**OBOC library synthesis**

An octapeptide OBOC library composed of D-amino acids (D-a.a.) was constructed on 90 μm Tentagel beads where each position was one of 18 of the conventional 20 a.a. (excluding cysteine and methionine to prevent the formation of oxidation products) using a standard split-mix synthesis approach,[2] to yield a library containing a theoretical diversity of more than $10^{16}$ different peptides, with each bead displaying $10^{13}$ copies of a unique peptide:

A photolabile linker, 3-amino-3-(2-nitrophenyl) propionic acid (ANP), was manually added to Tentagel resin beads using standard solid phase peptide synthesis (SPPS) using a fluorenylmethyloxycarbonyl (Fmoc) protecting group startegy. The remaining synthesis process was conducted in an automated peptide synthesizer (Biotage Syro Wave, Charlotte, NC). The resin was kept in the dark throughout the synthesis. Briefly, 20% piperidine in *N*,*N*-dimethylmethanamide (DMF, 800 μL/well, x2) was used for Fmoc deprotection. A different D-a.a. (3 eq.) was coupled in each well with *O*-(1*H*-6-chlorobenzotriazole-1-yl)-1,1,3,3-tetramethyluronium hexafluorophosphate (HCTU, 3 eq.) and *N*,*N*-diisopropylethylamine (DIPEA, 6 eq.) in DMF. A total of 18 wells (18 a.a.) were used, one for each of the common D-isomer a.a., excluding Cys and Met to avoid oxidation products. After each coupling step, the resin was rinsed with DMF and dichloromethane (DCM) several times and recombined in a peptide vessel and mixed thoroughly before being split again into the synthesizer wells for another round of Fmoc deprotection and coupling. The process of deprotection and coupling cycle was repeated until the library reached the desired length of eight a.a.

**OBOC library screening**

Library beads (500 mg) were washed 2x using DMF, 2x with methanol, 1x with 5% DIPEA in DMF, 3x in pure DMF, 3x in DCM, and then finally in 50% DMF in water to completely remove all unbound reagents. In the end, ethanol (70% in water) was added to the library beads to remove traces of organic solvents, and the beads were then re-suspended in phosphate buffer saline (PBS). The screening strategy (outlined in **Fig. 2**) employed a magnetic capture technique that we have previously described.[4,5] The a.a. 535–548 peptide sequence of dg-Bcan (epitope of BG1, and referred to as "dg-Bcan-peptide") as well as a corresponding scramble (scramble(dg-Bcan)) peptide was synthesized with a biotin on the N-terminus, and then conjugated onto small streptavidin-coated magnetic particles (2 µm size). The magnetic particles displaying the a.a. 535–548 peptide sequence were then used to capture high-affinity peptide ligands displayed on the OBOC library beads (Screen 1). Specifically, a 'negative screen' was first conducted to minimize the incidence of false positives by incubating the OBOC library with magnetic particles coated with the scramble(dg-Bcan) peptide. OBOC beads that exhibited high association with the 'scramble-particles' (likely through non-specific interactions) became highly magnetized, and the magnetized beads were sorted from the non-magnetic population using a magnetic-activated microfluidics device as previously described.[6] The magnetized beads isolated at this stage were discarded, and the non-magnetic population was subjected to the 'positive screen' using the 'dg-Bcan-particles', enabling high-throughput magnetic labeling of OBOC beads which presented peptide candidates with high affinity for the dg-Bcan-peptide (**Fig. 2b)**. The magnetized 'hit' beads were isolated using the microfluidic device (described above), and washed extensively with ethanol and PBS to remove the bound magnetic particles. Next, a secondary cell-based assay (Screen 2) was performed using dg-Bcan-expressing and non-expressing cells (**Fig. 2d,e,f**) to increase the likelihood of attaining affinity peptides that will not only bind purified dg-Bcan protein, but also target dg-Bcan-expressing cells. To ensure specificity, 'hit' beads from Screen 1 were first incubated with a confluent plate (10 cm) of live adherent astrocytes (dg-Bcan-negative) overnight at 37ºC, and beads (a few hundred) that did not associate with any cells were collected using a micropipette under an inverted dissecting microscope. These beads were then washed with ethanol, followed by a few washes in

Dulbecco's Modified Eagle Medium (DMEM) supplemented with 10% fetal bovine serum (FBS), and then incubated with a confluent plate (10-cm) of adherent U87 cells that do not express dg-Bcan overnight at 37°C. Beads that did not associate with the U87 cells (approx. 100 beads) were collected (using a micropipette under a dissecting microscope), and then incubated in a similar manner with dg-Bcan-overexpressing U87 (U87-Bcan) cells. 'Hit' beads with the highest interaction with U87-Bcan cells (n = 7) were isolated, washed extensively with ethanol and water, and then sequenced by Edman degradation.

**FITC-labeling**

The synthesis workflow is demonstrated in **Fig S2**. The peptide sequences were synthesized using standard Fmoc-solid phase peptide synthesis conditions on a Biotage Syrowave automated microwave peptide synthesizer. Fmoc-protected Rink Amide MBHA resin was used as a solid support (0.37 mmol/g). After swelling the resin in DCM, Fmoc removal was achieved with two subsequent treatments of 40% piperidine in DMF (1.2 mL) for 30 sec and 12 min, respectively. Coupling conditions for each amino acid were Fmoc-protected amino acid (4 eq.), HCTU (4 eq.) and 8 equivalents of DIEA (8 eq.9 in DMF/NMP over 1 hr). After completion of the peptide sequence on the peptide synthesizer, Mtt was manually removed with 2 mL of deprotection solution (93% DCM, 2% trifluoroacetic acid (TFA), 5% triisopropylsilane (TIPS)) at room temperature (10x2min). The resin was neutralized with a wash of 5% DIEA in DMF. Subsequently, fluorescein isothiocyanate (FITC) was conjugated to the lysine side-chain with FITC (2 eq.) and DIEA (6 eq.) in 2 mL of DMF for 18 hrs. Final N-terminal Fmoc removal was performed through two cycles of 20% piperidine in DMF (3 min and 15 min). The peptide was subjected to simultaneous global side-chain deprotection and cleavage from resin by treatment with (95% (v/v) TFA, 2.5% (v/v) TIPS, 2.5% (v/v) water) at room temperature over 5 hrs. The peptide was precipitated in cold TBME and centrifuged at 1700 rcf for 15 min. The TBME was decanted, the peptide dissolved in water, frozen at -78 °C and lyophilized. The peptide was then purified by preparative RP-HPLC using an Agilent Zorbax PrepHT SB-C18 column (21.2x150 mm, 5 µm particle size) at a flow rate of 20 mL/min with various gradients acetonitrile (0.1%TFA) in water (0.1% TFA). After purification, the collected fractions were frozen at -78 °C and lyophilized. Peptides were characterized by ESI+ mass spectrometry using a Waters Micromass Quatro Micro Mass Spectrometer.

**Cy5.5-labeling**

The synthesis workflow is demonstrated in **Fig. S4**. The peptides were synthesized using an automated flow-based synthesizer using Fmoc chemistry and H-Rink Amide-ChemMatrix resin (200 mg) as previously reported.[3] The C-terminus of each peptide was coupled to F-moc-6-aminohexanoic acid, followed by F-moc-L-propargylglycine to functionalize the C-terminus of the peptide with an alkyne. The peptide was then cleaved from the resin and purified using RP-HPLC on a Zorbax $C_3$ or the $C_{18}$ column (Agilent) as detailed below. Cy5.5-azide was conjugated using copper-catalyzed "click" chemistry. Briefly, 4 µmol of peptide-alkyne and 4 µmol of Cy5.5-azide was dissolved in a 50:50 mixture of $H_2O$/t-butanol. Then, 100 µL of 500 mM Tris and 50 µL of 100 mM $CuSO_4$ (both dissolved in $H_2O$) were added to the peptide-dye mixture. Then, 10 µL of tris(benzyltriazolylmethyl)amine (TBTA; dissolved in DMSO), 10µL of tris(2-carboxyethyl)phosphine (TCEP; dissolved in $H_2O$) and 100 µL of ascorbic acid were added. 480 µL of a 50:50 mixture of $H_2O$/*t*-butanol were added in the end to generate a final volume of 1 mL. The combined mixture was allowed to incubate at room temperature for at least 3 hrs. The labeled peptide was separated from unlabeled by RP-HPLC on a Zorbax $C_3$ or the $C_{18}$ column using the parameters below.

The following liquid chromatography (LC) method was used:

A = water 0.1% TFA; B = acetonitrile 0.1% TFA; flow rate = 0.8 mL/min

0–2 min 5% B,

2–11 min linear gradient 5–65% B

11–12 min 65% B

Column: Zorbax 300SB $C_3$ or the $C_{18}$ column (9.4 × 250 mm, 5 µm), 40 ºC

**Deglycosylation of recombinant brevican protein**

The brevican protein was deglycosylated in water + 0.05% tween, sialidase, *O*-glycosidase, *N*-glycosidase in sodium acetate [200mM], NaCl [250mM], Tris-HCl [200mM], pH 7.5.) in the presence of protease inhibitor overnight at 4ºC. This entire composition is referred to as 'deglycosylation buffer' throughout the manuscript.

**Octet binding kinetic analyses**

The FortéBio OctetRed384 was used to study the binding kinetics of each peptide to recombinant human brevican (in PBS and 0.1 mM EDTA, pH 6.8). The brevican protein was deglycosylated prior to experimental use as detailed above. All binding kinetics assays were performed within the OctetRed instrument under agitation at 1000 rpm in 0.9% NaCl irrigation with 0.05% Tween (working buffer). Assays were performed at 30°C in solid black 384-well plates (Geiger Bio-One). The final volume for all the solutions was 80 µl/well. Firstly, Ni-NTA biosensors were soaked for 10 min in working buffer. Before loading the protein onto each biosensor, a baseline was established in working buffer for 60 s. Deglycosylated brevican, dg-Bcan (50 µg/ml; His-tagged) was loaded on the surface of each biosensor for 180 s. Typical capture levels varied slightly between 0.5 and 2 nm, and variability within run did not exceed 0.1 nm. Reference biosensors were exposed to the 'de-glycosylation buffer' lacking brevican protein during the loading step as internal controls. A 60 s biosensor washing step was applied. Then, biosensors were exposed to the analyte (peptide) in working buffer (ranging between 0–10 μM BTP) for 300 s during the 'association' step. Finally, the biosensors were exposed to working buffer (without peptide) during the 'dissociation' step for 600 s. Binding affinity of each peptide was assessed through steady state analysis, where the response unit was plotted over peptide concentration. All data was plotted and curves were fitted using the non-linear 'one-site specific binding' fit, and the dissociation constant ($K_D$) value was calculated using the GraphPad Prism software.

**UV-Crosslinker-labeling**

The synthesis workflow is demonstrated in **Fig. S3**. H-Rink Amide-ChemMatrix resin (resin loading: 0.49 mmol/g, 0.10 g, 49 μmol) was pre-coupled to Fmoc-L-Lys(biotin)-OH. BTP-7 containing an Fmoc-L-Lys(alloc)-OH was synthesized using an automated flow-based synthesizer using Fmoc chemistry as previously reported.[1] After synthesis, the peptidyl resins were washed with DCM (3x5 mL) and dried under reduced pressure. To each resin was added a mixture of di-*tert*-butyl dicarbonate ($Boc_2O$, 33 mg, 0.15 mmol, 14 eq.) and DIEA (0.12 mL, 0.68 μmol, 14 eq.) in DCM (1.6 mL) and the reaction proceeded at room temperature for 45 min. Afterwards, the resin was washed with DCM (3x5 mL). Then, a mixture of tetrakis(triphenylphosphine)palladium(0) (30 mg, 26 μmol, 0.5 eq.) and piperidine (0.25 mL, 0.29 mg, 3.4 mmol, 70 eq.) in DCM (1.5 mL) was added to each peptidyl resin at room temperature. After 30 min, the resins were washed with DCM (3x5 mL) and dried under reduced pressure.

NHS-diazirine (38 mg, 0.17 mmol, 7 eq.) and DIEA (98 uL, 0.57 mmol, 23 eq.) in DMF (2.5 mL) were then added to half of each peptidyl resin (~25 μmol) at room temperature and the mixture was kept in the dark for 60 min. The resins were washed with DCM (3x5 mL) and dried under reduced pressure.

The resin was transferred into a 50 mL conical polypropylene tube covered in aluminum foil and 3 mL of cleavage solution (82.5% TFA, 5% water, 5% phenol, 5% thioanisole, 2.5% EDT) was added. The tube was kept on a nutating mixer at room temperature for 4 hrs. Ice cold diethyl ether (45 mL) was added to the cleavage mixture and the precipitate was collected by centrifugation (1700 rcf, 2 min, 4 °C) and triturated twice more with cold diethyl ether (45 mL). The supernatant was discarded. Residual ether was allowed to evaporate and the peptide was dissolved in 50% acetonitrile in water with 0.1% TFA. The peptide solutions were filtrated with a Nylon 0.22 µm syringe filter, frozen and lyophilized in the dark.

Purification by RP-HPLC on a semipreparative Agilent Zorbax 300SB-C3 column (9.4 mm×250 mm, 5 µm particle size) at a flow rate of 4 mL/min with an acetonitrile gradient (1–31% over 60 min) gave BTP-7-diazirine (2.1 mg, 0.94 mol, 4%). Purity of each RP-HPLC fraction was confirmed by LC-MS.

**Competitive binding assay**

BTP-7 was functionalized with a C-terminal diazirine and biotin to allow for UV-activated crosslinking and western blot detection with streptavidin-horseradish peroxidase (HRP) (see SI for chemical structure and synthesis). For the dose dependent blocking study a 50 µM stock of native BTP-7 was diluted 1:2 with 0.9% NaCl and 15 µl of each dilution were plated in a 96-well plate. From this point on light exposure was limited and the experiment performed under dim lighting conditions. dg-Bcan protein and the BTP-7-biotin-crosslinker were added into each well of the 96-well plate (final concentrations: dg-Bcan (0.041µM), BTP-7-biotin-crosslinker (4.2 µM), native BTP-7 (20-0µM)). The plate was incubated at room temperature (RT) for 1 hr and placed on ice in a UV crosslinker (Fisherbrand, 13-245-221) at 368 nm wavelength for 20 min. The samples were resolved by sodium dodecyl sulfate polyacrylamide gel electrophoresis (SDS-PAGE) and transferred to a nitrocellulose membrane. Membranes were blocked with 5% BSA in TBST for 1 hr at RT. To measure the overall dg-Bcan level, membranes were probed with the primary pan-Bcan antibody (1:20,000 dilution), followed by HRP-conjugated secondary antibody. To detect the crosslinked BTP-7-dg-Bcan entity, membranes were stripped with 6 M guanidine hydrochloride, washed thoroughly with TBST, and then probed with streptavidin-HRP (1:10,000). The streptavidin-HRP signal was quantified and normalized against the corresponding dg-Bcan protein level using ImageJ.

**Circular dichroism**

The dg-Bcan peptide (or control dg-Bcan(Scramble) peptide), BTP-7-FITC peptide, or a mixture of both (dissolved in Milli-Q $H_2O$ to a final concentration of 50 μM for each peptide) was loaded into a quartz cuvette (1 mm pathlength; Hellma Analytics, Müllheim, Germany, Catalog # 110-1-40). The cuvette was inserted into the Jasco J-815 circular dichroism spectropolarimeter, and the spectral scan measurement was conducted using the Spectrum Manager 2 software. The cuvette was rinsed with Milli-Q $H_2O$ in between each run. A blank (water) was run to establish the baseline for all samples. Scans were measured from 190 nm to 260 nm in a continuous mode (50–100 nm/min) at 20 °C. To measure the BTP-7 structural change in the presence of either the dg-Bcan (or control scramble) peptide, spectrum of the dg-Bcan (or scramble) peptide alone was subtracted from the spectrum of the mixture, and then overlaid with the spectrum of BTP-7 alone for comparison. To investigate the structural changes in BTP-7 in a dose-dependent manner, 50 μM of BTP-7 peptide was loaded together with varying concentrations of either the dg-Bcan (or control scramble) peptide. Similarly, the spectrum of each dg-Bcan (or control scramble) peptide at the concentration was subtracted from the spectrum of the corresponding mixture, and then overlaid with the spectrum of BTP-7 alone for comparison.

**Serum stability Assay**

To test for serum stability, a 10 mM stock of each compound was prepared in DMSO. 1.4 µL of each stock solution was transferred into a microcentrifuge tube containing 200 µL of PBS with 25% human serum and incubated at 37 °C. At timepoints 0, 0.25, 0.5, 1, 3, 6, 12 and 24 hrs, 10 µL of each sample were removed from the microcentrifuge tube and transferred to a different microcentrifuge tube containing 20 µL of a solution of 6 M GuHCl and 1 M EDTA. Then, 10 µl of these time-point samples were purified via solid-phase extraction with Millipore $C_{18}$ 10 µL ziptips. Samples were eluted with 10 µL of 70% acetonitrile in water containing 0.1% TFA into an LC-MS vial containing 20 µL of water with 0.1% TFA additive and analyzed *via* LC-MS. Remaining intact compound was determined through an extracted ion current (EIC) chromatogram using MassHunter software. Each timepoint was performed in triplicate.

**Cell invasion assay**

GBM-6 neurospheres (GSC) were plated on a 96-well plate in 25 µL volume per well in the center of the well. Once plated, spheres were allowed to adhere to the plate for approximately 2–3 hrs. Collagen solution PureCol Collagen (Cat. # 5015) was prepared using 5x DMEM basal media, and supplements (EGF, FGF and B27 in the appropriate proporation) were added to the 5x DMEM. Then, the pH was neutralized to 7.5 using 1 N NaOH. BTP-7 (at 50 µM final concentration) was added into the collagen solution. Then, 50 µL of the collagen mix was added into each well containing the GBM-6 neurospheres, and the solution allowed to polymerize at 37 °C for 1 hr. Media containing 50 µM of BTP-7 was added into each well (100 µL volume) on top of the polymerized collagen. Live imaging of the neurospheres was performed under an inverted microscope over 48 hrs (snapshots were captured at every 20-min interval) within a 37 °C chamber supplied with $CO_2$ (using a 4x objective).

**Cell proliferation assay**

The CellTitre-Glo® luminescent cell viability assay was used. GBM-6 glioma stem cells [7] (cultured in Neurobasal growth media), or HEK cells (cultured in supplemented DMEM media) were washed once with PBS, dissociated to form single cell suspension using either StemPro Accutase for GBM-6 neurospheres, or 0.05% (v/v) Trypsin-EDTA for HEK cells, resuspended in the appropriate growth media, and then counted using a hemocytometer. Cells were seeded at the desired number (typically 20,000–30,000 GBM-6 cells per well; 5,000–20,000 HEK cells per well) into clear-bottom black-well 96-well plates (Greiner BIO-ONE, Monroe NC; Cat. # 655090) in triplicates. A serial dilution of the camptothecin (CPT)-conjugated to BTP-7 (BTP-7-CPT) or scramble (Scr-7-CPT) (dissolved/diluted in DMSO) was prepared so that the following final working concentrations on cells were achieved: 100, 33.3, 11.1, 3.70, 1.24, 0.412, 0.137, 0.046, 0.015, 0.005 µM. The drug at each dilution was added onto cells in triplicate (final volume of media is 100 µM). Vehicle (DMSO with no drug) was added as a control. The plates were returned into a 37 °C tissue culture incubator. At every 24 hrs after incubation, a plate was removed from the incubator and 100 µL of CellTiter-Glo working reagent was added to each well using a multi-channel pipette. The plate was allowed to incubate (in the dark) for 10 min (while shaking under low speed), and then analyzed in a luminescence plate reader (POLARstar Omega, BMG Labtech, Offenburg, Germany). Each dataset was normalized to the vehicle (control), and plotted using GraphPad Prism. $IC_{50}$ values were obtained using a non-linear fit (log(inhibitor) vs. response - variable slope (four parameters)).

**BBB organoid assay**

Immortalized hCMEC/D3, primary HBVP and primary human astrocytes were used for generation of BBB organoids as previously described.[8,9] Briefly, agarose (0.5 g) was dissolved in 50 mL of MilliQ water or PBS in a conical flask and the solution was heated in a microwave for 1–2 min until completely dissolved. Melted agarose was transferred into a sterile tissue culture hood and immediately poured into a 50-mL sterile basin. 50 μL of melted agarose was dispensed into each well in a 96-well plate using a multichannel pipette. The plate was set aside to allow agarose to solidify completely inside the wells for at least 15 min. Then, each cell type (hCMEC/D3, HBVP and astrocytes) was released from the flask using 1–1.5 mL of Trypsin-EDTA by incubating for 1–3 min at 37 °C in an incubator. Once cells were detached, 9 mL of 'BBB working media' was used to neutralize the trypsin and resuspend the cells. Each cell type was counted using a hemocytometer, and 1,500 of each cell type were seeded into each well containing solidified agarose. Cell density was verified under an inverted microscope after each seeding step. BBB working medium was used to adjust the final volume to 200 μL. The plate was placed in at 37 °C incubator, and the organoids allowed to form in culture for 48 hrs.

For peptide permeability analyses, after 48 hrs, organoids that formed successfully were pooled together into a 0.5-mL microcentrifuge tube in BBB working medium. The test peptide (Cy5.5-labeled) was added to the tube containing the organoids (at a final concentration of 10 μM) in 500 μL of fresh BBB working medium. To test the integrity of the organoid surface, the organoids were co-incubated with the peptide together with fluorescent dextran (rhodamine-dextran (4,400 Da)) at a final concentration of 10 μg/mL for 3 hrs at 37 °C in an incubator under constant rotation. The organoids were then washed 3x with 500 μL of BBB working medium. For nuclei staining, Hoechst dye (1:1000) was incubated for 1 min at room temperature, and the organoids were washed 3x with 500 μL of BBB working medium. The organoids were then transferred onto a thin-well chambered cover glass and imaged by confocal fluorescence microscopy. Quantification of peptide permeability was performed using the ImageJ software, where the mean fluorescence intensity of the core of each spheroid (at a depth between 50–90 μm) was measured, and then plotted on GraphPad Prism.

**Biodistribution studies**

Lyophilized BTP-7-Cy5.5 powder was dissolved to a final concentration of 10 mM in DMSO by mass. The peptide was then diluted to a concentration of 100 μM in a solution of 50:50 polyethylene glycol (PEG)-300:0.9% sodium chloride (v/v) irrigation solution. A 100 μL dose of each peptide solution was administered intravenously *via* the tail vein into healthy 8-week old female nude mice. After 4 hrs, 100 μL of 50 mg/mL tetramethylrhodamine isothiocyanate (TRITC)-labeled dextran (155 kDa) in 0.9% sodium chloride was injected via the tail vein. Mice were sacrificed 30 min later by cervical dislocation. The brain, heart, lungs, kidneys, spleen, and liver were excised from each mouse, frozen on dry ice and imaged using an *In Vivo* Imaging System (Perkin Elmer, Waltham, MA) at an excitation of 640 nm. Using Living Image software, regions of interest were drawn around each organ and the total radiant efficiency of Cy5.5 from each organ was quantified.

**BBB permeability in vivo**

To measure BBB penetration, naïve nude mice were injected intravenously *via* the tail vein with either Cy5.5-labeled BTP-7 or a scramble (Cy5.5-Scr-7) peptide (100 μL of 100 μM peptide solution). 4 hrs later, mice were injected with 100 μL of 50 mg/mL TRITC-dextran (155 kDa) *via* the tail vein. Mice were euthanized after 30 min, and their brains excised, frozen and cryo-sectioned into 16 μm slices. Tissue sections from the frontal lobe were imaged by confocal microscopy using a 20x objective. Areas with high TRITC (red) signal indicate regions of high perfusion (*i.e.* blood vessels). Cy5.5 (peptide) intensity was measured in areas outside of visible

TRITC signal to ensure that only peptide intensity in the brain parenchyma (peptide that has successfully extravasated from the vessels and accumulated in the brain tissues) was quantified.

#### Supplementary Figures

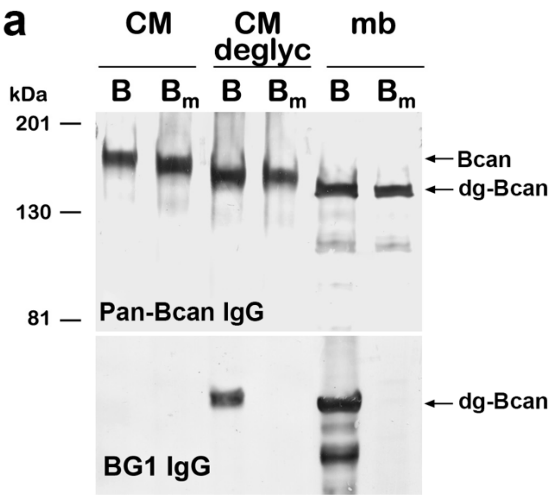


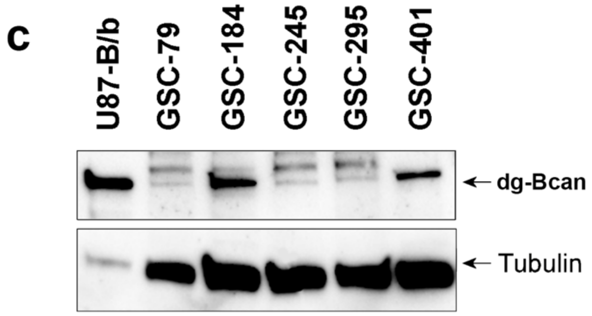


| Sample ID | Age | Gender | Grade | Recurrent | MGMT Methylation |
|---|---|---|---|---|---|
| GS79 | 74 | F | 4 | Newly Diagnosed | Negative |
| GS184* | 50 | M | 4 | Newly Diagnosed | Methylated |
| GS245 | 68 | M | 4 | Newly Diagnosed | Negative |
| GS295 | 50 | M | 4 | Newly Diagnosed | Mixed |
| GS401 | 52 | M | 4 | *Recurrent of GS184 | Methylated |

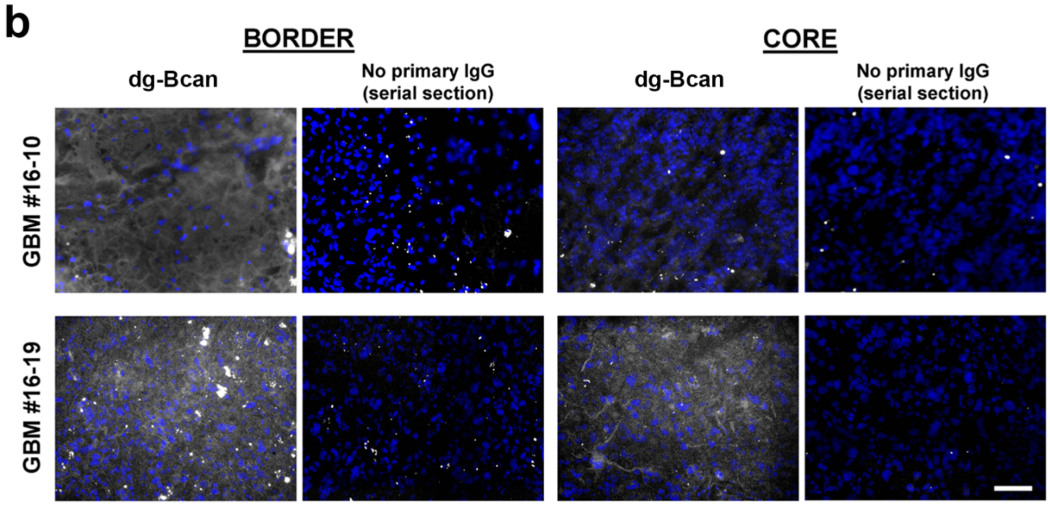


**Fig S1. Expression of dg-Bcan in high-grade glioma tumor samples and glioma stem cells**
**a)** Western blot analysis of culture medium (CM), culture medium + deglycosylases (CM deglyc) and cell membranes (mb) of U87 cells overexpressing either Bcan or a mutated version of Bcan ($B_m$). In Bm cells three threonine residues within the a.a. 535–548 were replaced with alanines. The pan-Bcan polyclonal antibody recognizes all Bcan isoforms (top), while BG1 antibody recognizes only the dg-Bcan isoform and not the glycosylated or mutated Bcan (bottom). **b)** Immunofluorescence images showing high level of dg-Bcan expression (white) in frozen biopsy GBM specimens obtained from the SUNY Upstate Medical Center. Biopsy sample from the tumor border or core isolated from the same patient were stained with a dg-Bcan-specific primary antibody. Scale bar: 100 microns (20x objective). **c)** Western blot analysis of dg-Bcan expression in glioma stem cells (cultured as neurospheres in Neurobasal medium) derived from patients with IDH-wildtype GBM (from the Erasmus Medical Center, Netherlands). Additional pathology information is listed in the table below.

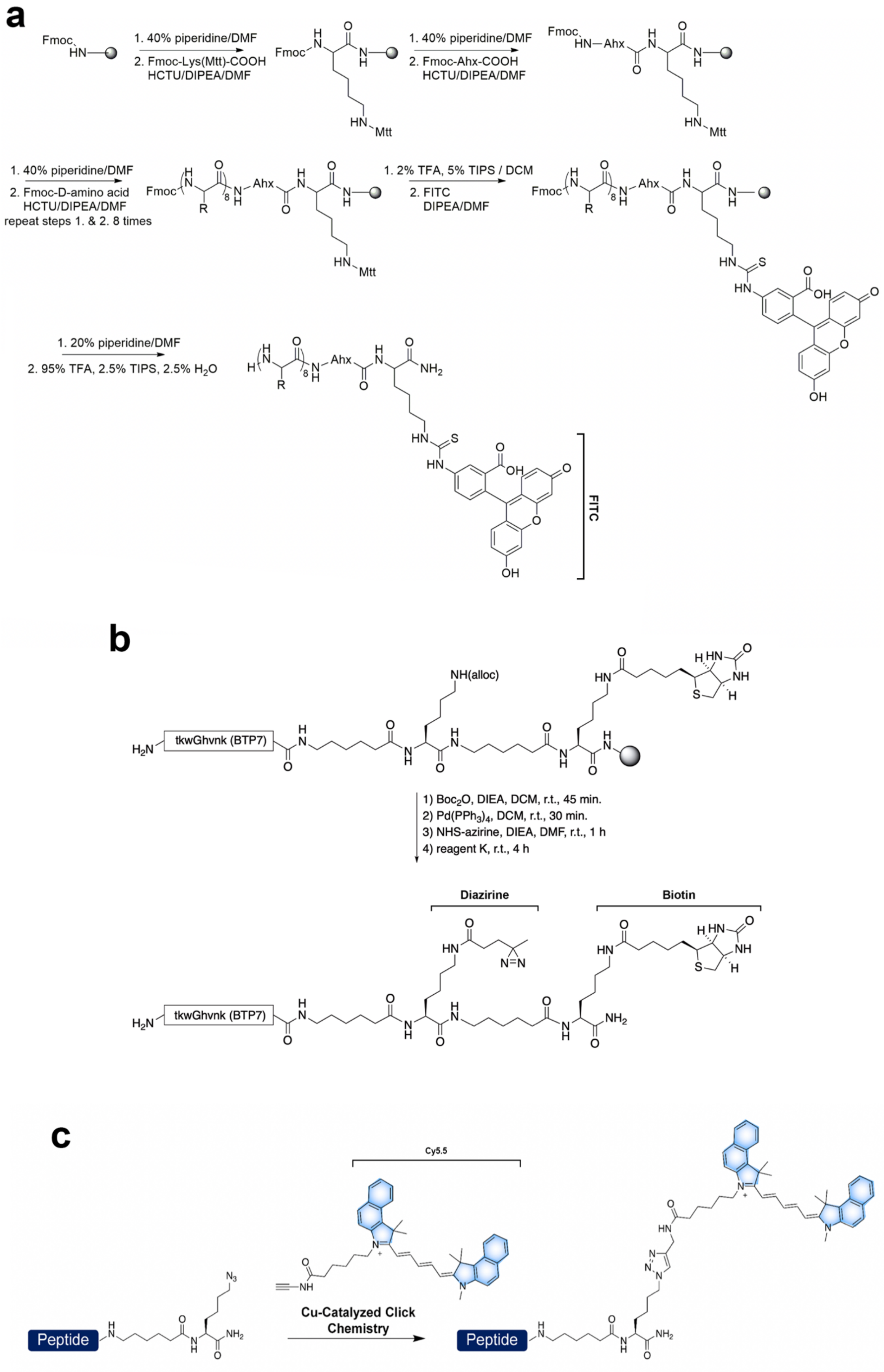
a
1. 40% piperidine/DMF
2. Fmoc-Lys(Mtt)-COOH HCTU/DIPEA/DMF
1. 40% piperidine/DMF
2. Fmoc-Ahx-COOH HCTU/DIPEA/DMF
1. 40% piperidine/DMF
2. Fmoc-D-amino acid HCTU/DIPEA/DMF
repeat steps 1. & 2. 8 times
1. 2% TFA, 5% TIPS / DCM
2. FITC DIPEA/DMF
1. 20% piperidine/DMF
2. 95% TFA, 2.5% TIPS, 2.5% H2O
FITC
b
tkwGhvnk (BTP7)
NH(alloc)
1) Boc2O, DIEA, DCM, r.t., 45 min.
2) Pd(PPh3)4, DCM, r.t., 30 min.
3) NHS-azirine, DIEA, DMF, r.t., 1 h
4) reagent K, r.t., 4 h
Diazirine
Biotin
c
Cy5.5
Cu-Catalyzed Click Chemistry
Peptide

**Fig. S2. Workflow for the synthesis of fluorescein (FITC)-labeled peptide (a), peptides for UV-Crosslinker labeling (b) and Cy5.5-labeled peptide (c)** (See methods)

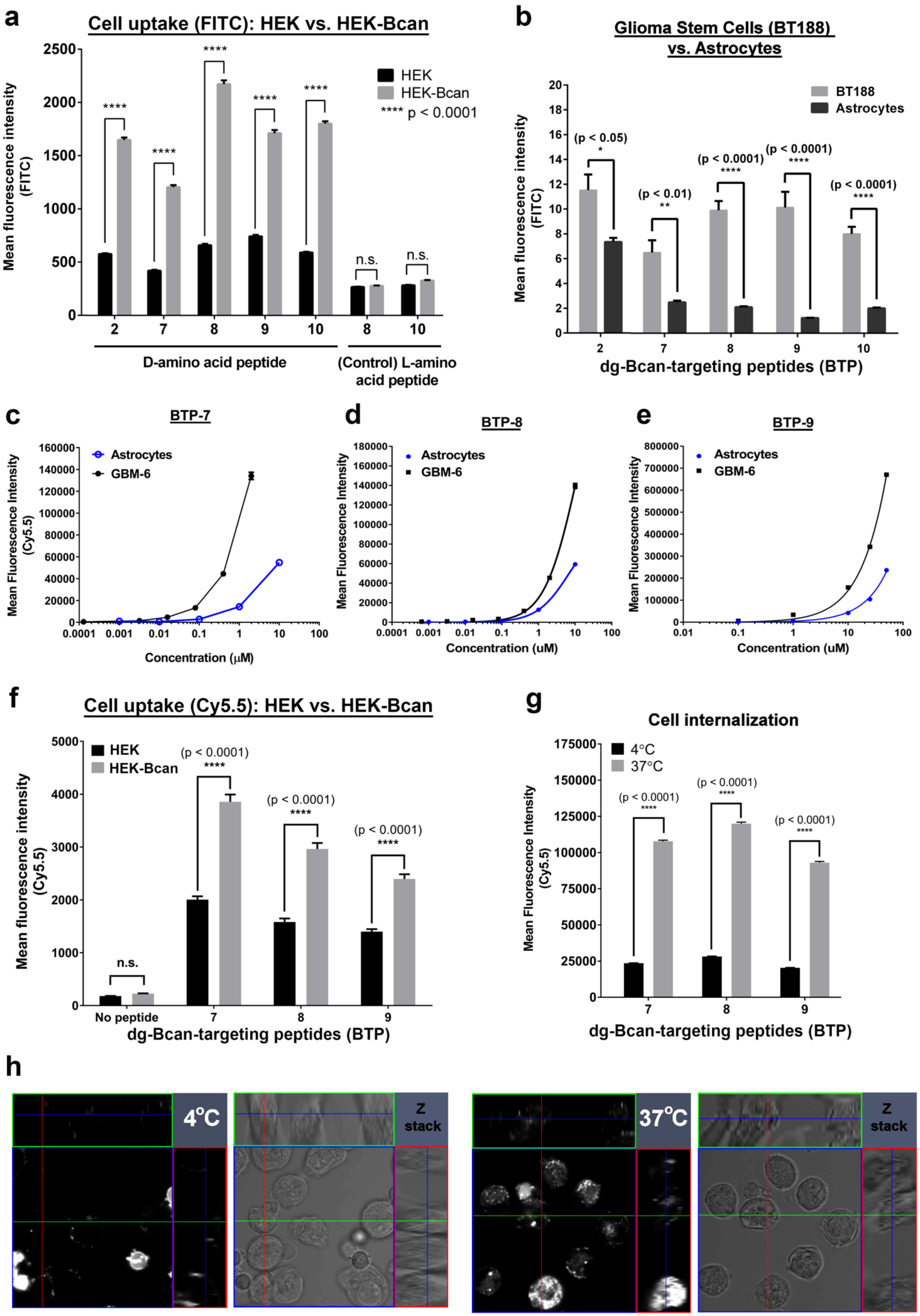
a
Cell uptake (FITC): HEK vs. HEK-Bcan
Mean fluorescence intensity (FITC)
HEK
HEK-Bcan
**** p < 0.0001
n.s.
D-amino acid peptide
(Control) L-amino acid peptide
b
Glioma Stem Cells (BT188) vs. Astrocytes
BT188
Astrocytes
(p < 0.05)
(p < 0.01)
(p < 0.0001)
dg-Bcan-targeting peptides (BTP)
c
BTP-7
d
BTP-8
e
BTP-9
Astrocytes
GBM-6
Mean Fluorescence Intensity (Cy5.5)
Concentration (μM)
Concentration (uM)
f
Cell uptake (Cy5.5): HEK vs. HEK-Bcan
Mean fluorescence intensity (Cy5.5)
No peptide
dg-Bcan-targeting peptides (BTP)
g
Cell internalization
4°C
37°C
Mean Fluorescence Intensity (Cy5.5)
dg-Bcan-targeting peptides (BTP)
h
4°C
Z stack
37°C
Z stack

**Fig. S3. Characterization of peptides for dg-Bcan specificity using cell uptake studies**

**a)** Cell uptake of fluorescein-conjugated BTP candidates in HEK (dg-Bcan-null) or HEK-Bcan cells by flow cytometry analysis. When BTP-8 and BTP-10 were mutated to L-amino acids, the increased peptide uptake in HEK-Bcan cells was abrogated ($n_{events}$ = 5,000, c = 10μM, Two-way ANOVA and Sidak's Multiple Comparison test). **b)** Uptake of fluorescein-labeled BTP candidates in BT188 (dg-Bcan-high) glioma stem cells (GSCs) or astrocytes (dg-Bcan-null) (c = 2μM, $n_{GSCs}$ = 20, confocal microscopy, two-way ANOVA and Sidak's Multiple Comparison test). **c**,**d**,**e**) Flow cytometry uptake analysis of Cy5.5-conjugated-peptides BTP-7,8 and 9 in GBM-6 GSCs (dg-Bcan-positive) in comparison to astrocytes ($n_{events}$ = 20,000). **f)** Flow cytometry uptake analysis of Cy5.5-conjugated BTP-7, 8 and 9 in HEK (dg-Bcan-null) or HEK-Bcan cells ($n_{events}$ = 20,000, c = 5 μM). **g)** Cell uptake of Cy5.5-BTP-7, BTP-8 and BTP-9 in GBM-6 cells at 4 °C (inhibition of endocytosis) and 37 °C by flow cytometry analysis ($n_{events}$ = 20,000, c = 5 μM). **h)** Representative fluorescence images acquired using confocal microscopy depicting higher BTP-7 uptake/internalization into GBM-6 cells at 37 °C compared to 4 °C.

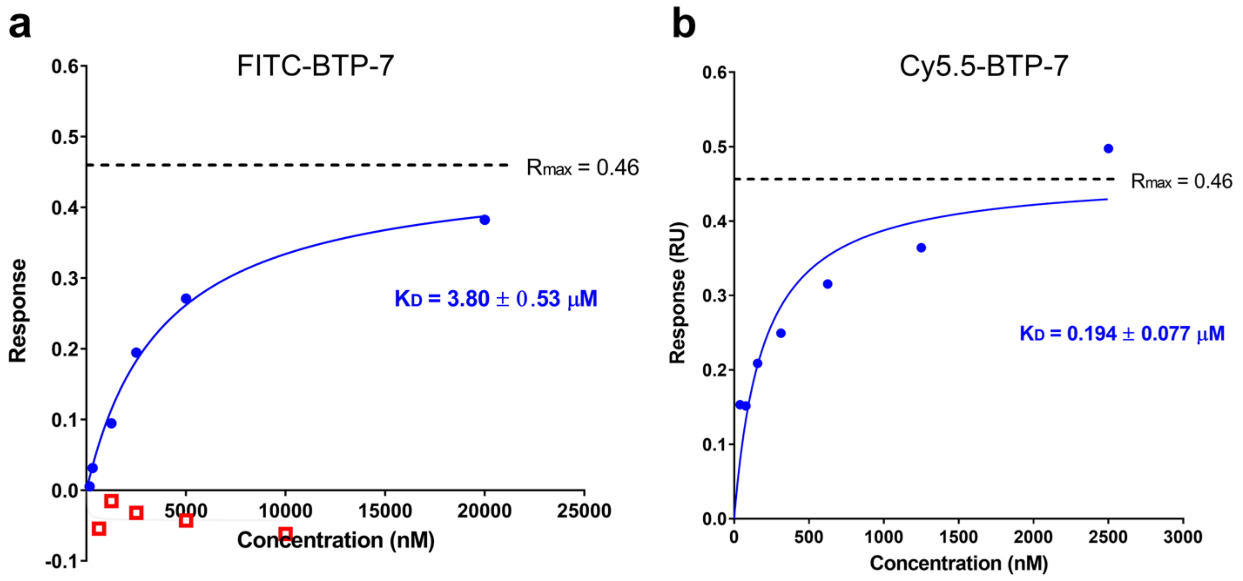


**Fig. S4. Binding affinity of functionalized BTP-7 to dg-Bcan protein**

**a)** Steady state analysis using the Octet platform (Forte Bio) measuring binding affinity of fluorescein-labeled BTP-7 to recombinant dg-Bcan (blue) or the fully glycosylated Bcan (red) protein. **b)** Steady state analysis measuring binding affinity of Cy5.5-labeled BTP-7 to recombinant dg-Bcan. The $K_D$ (dissociation constant) of each peptide to dg-Bcan is indicated. All steady state curves were plotted on the GraphPad Prism software (version 7.03), and the $K_D$ and $R_{max}$ values were measured through a non-linear regression (one site specific binding) fit.

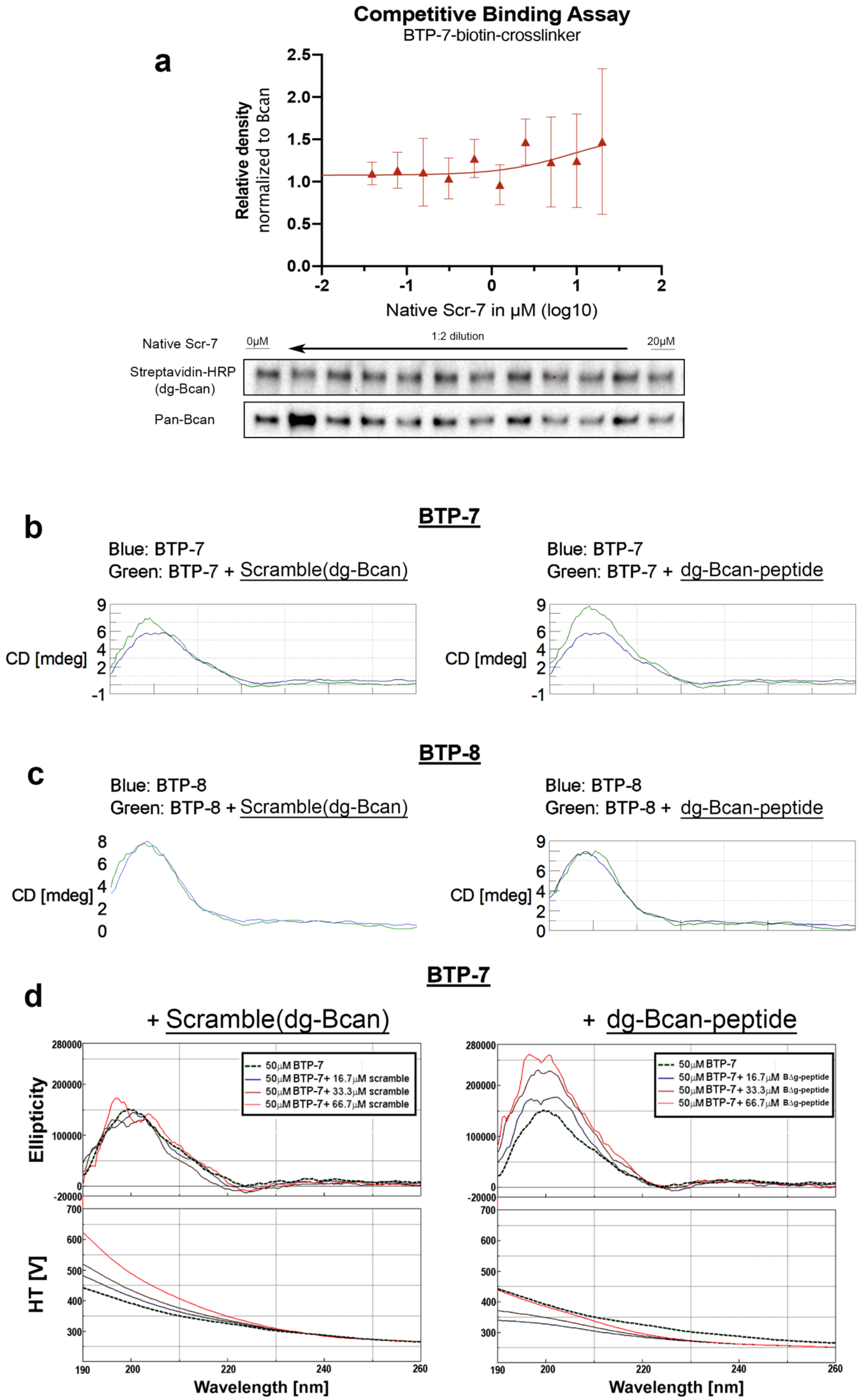
a
Competitive Binding Assay
BTP-7-biotin-crosslinker
Relative density
normalized to Bcan
Native Scr-7 in μM (log10)
Native Scr-7
0μM
1:2 dilution
20μM
Streptavidin-HRP
(dg-Bcan)
Pan-Bcan
b
BTP-7
Blue: BTP-7
Green: BTP-7 + Scramble(dg-Bcan)
Blue: BTP-7
Green: BTP-7 + dg-Bcan-peptide
CD [mdeg]
c
BTP-8
Blue: BTP-8
Green: BTP-8 + Scramble(dg-Bcan)
Blue: BTP-8
Green: BTP-8 + dg-Bcan-peptide
CD [mdeg]
d
BTP-7
+ Scramble(dg-Bcan)
+ dg-Bcan-peptide
50μM BTP-7
50μM BTP-7+ 16.7μM scramble
50μM BTP-7+ 33.3μM scramble
50μM BTP-7+ 66.7μM scramble
50μM BTP-7+ 16.7μM BΔg-peptide
50μM BTP-7+ 33.3μM BΔg-peptide
50μM BTP-7+ 66.7μM BΔg-peptide
Ellipticity
HT [V]
Wavelength [nm]

**Fig. S5. Binding of BTP-7 to dg-Bcan and the dg-Bcan-derived peptide**

**a)** Competitive binding assay. Western blot analysis showing no dose-dependent blocking of BTP-7 binding to dg-Bcan occurred in the presence of native Scr-7 peptide (Representative blot shown, n=3) **b)** Circular dichroism (CD) spectra of fluorescein-labeled BTP-7 (50 μM conc.) in the presence of either 50 μM dg-Bcan-peptide (residue 535–548) or the scramble dg-Bcan-peptide (see **Fig. 2b**). Spectra of BTP-7 alone are represented in green, while the spectra of BTP-7 in the presence of dg-Bcan-peptide (or the scramble peptide) are in blue. The spectrum of BTP-7 in the presence of dg-Bcan-peptide is generated by subtracting the spectrum of 'dg-Bcan-peptide alone' from the spectrum of BTP-7 mixed with the dg-Bcan-peptide. **c)** CD spectra of fluorescein-labeled BTP-8 (50 μM conc.) in the presence of either dg-Bcan-peptide or the scramble peptide (as in b)). **d**) CD spectra of BTP-7 in the presence of increasing concentration of the dg-Bcan (or scramble)-peptide, showing a change in the BTP-7 structure in the presence of dg-Bcan-peptide in a dose-dependent manner (0–67 μM). The spectra of BTP-7 remain relatively unchanged in the presence of the scramble peptide. All CD experiments are conducted at 20 ºC. The BTP-7 spectrum peaks at 200 nm, and the high tension (HT) voltage are ensured to be below 600 at that wavelength to avoid detector saturation.

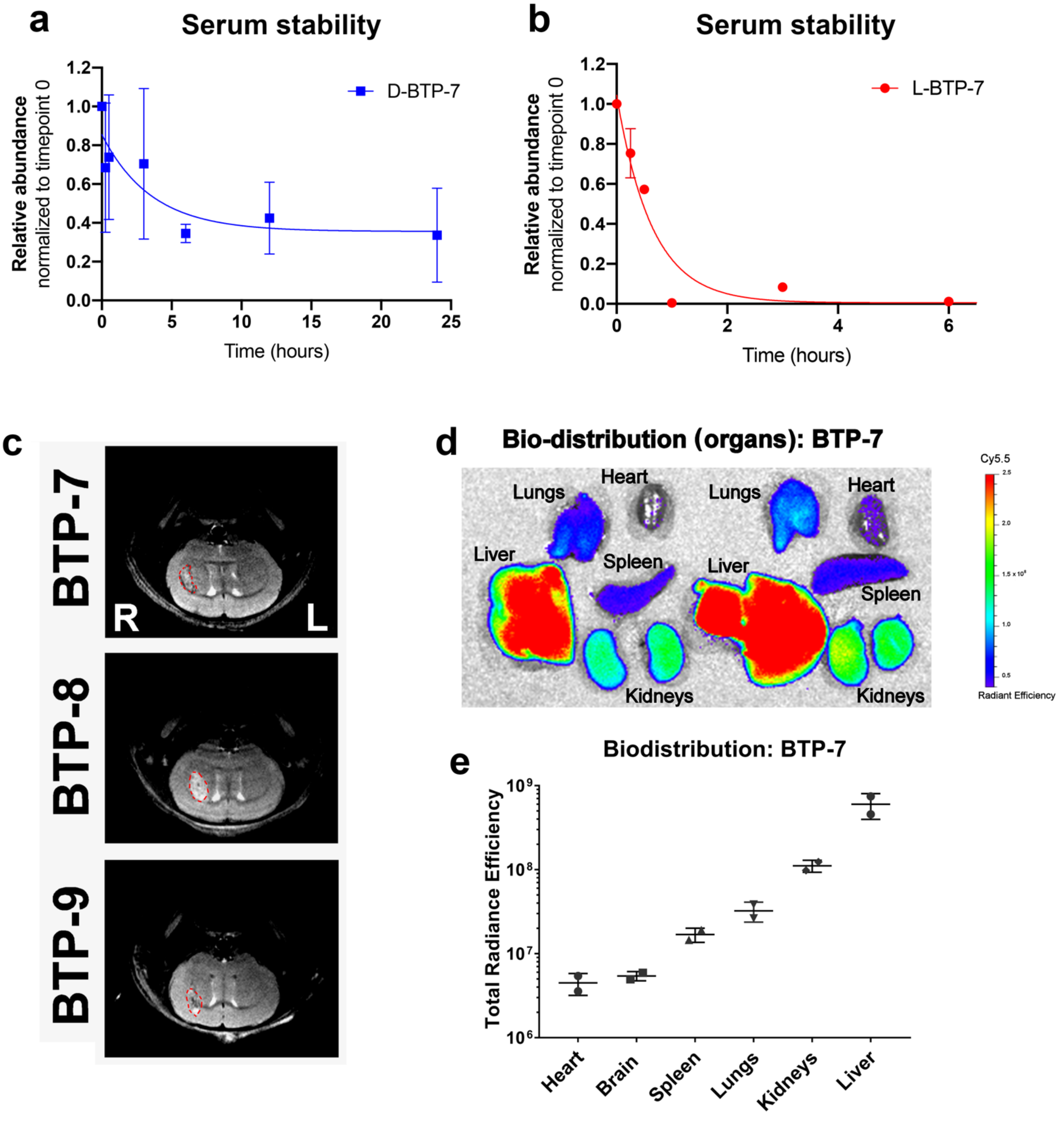


**Fig. S6.** ***In vivo*** **analysis of BTP-7**

**a**,**b)** Serum stability profile of D-BTP-7 and L-BTP-7. Curves were fitted using one phase decay nonlinear regression in Graphpad prism. **c)** Representative coronal MRI images of GBM-6 xenograft tumors (delineated with red dotted lines) prior to injection with Cy5.5 labeled BTP-7, 8 or 9. **d)** Biodistribution analysis of Cy5.5-labeled BTP-7 in mouse heart, lungs, liver, spleen and kidneys at 4 hrs post i.v. injection (100 µL of 100 µM peptide). Images of all organs were obtained by IVIS imaging after excitation at 640 nm (n = 2). **e)** Quantification of the total Cy5.5 radiance efficiency in all mouse organs.

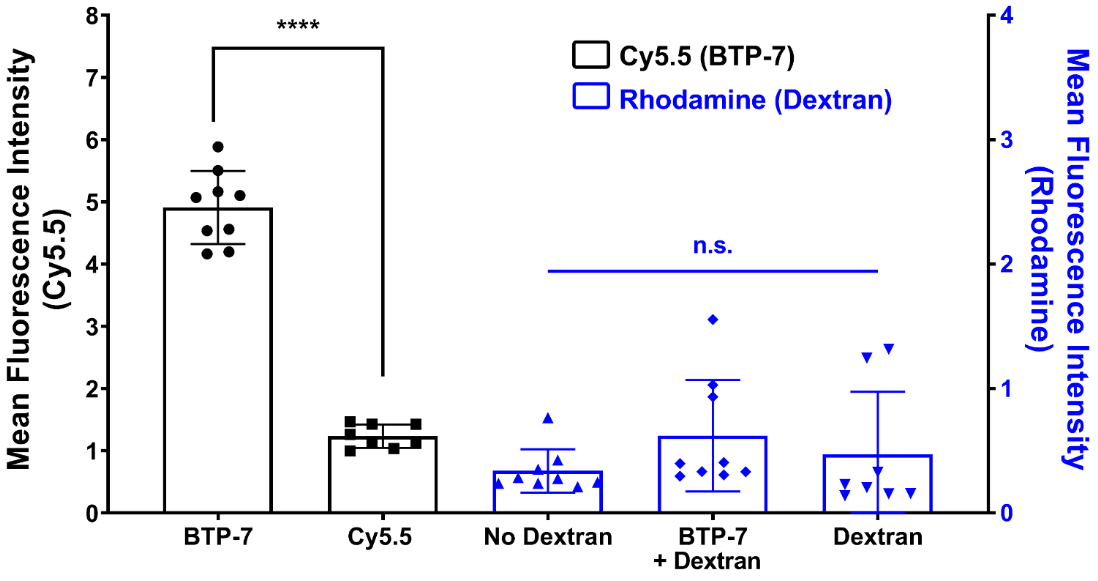


**Fig. S7. BBB permeability of BTP-7 *in vitro***
Quantification of mean fluorescence intensity in the core of BBB organoids after co-incubation with BTP-7-Cy5.5 and 4.4 kDa rhodamine dextran ($c_{dextran}$ = 10 µg/ml, $c_{peptides}$ = 10µM, t=3 hrs). BBB integrity within the organoids was not affected by BTP-7. All statistics were performed using the one-way ANOVA and Tukey's Multiple Comparison test (**** $p < 0.0001$)

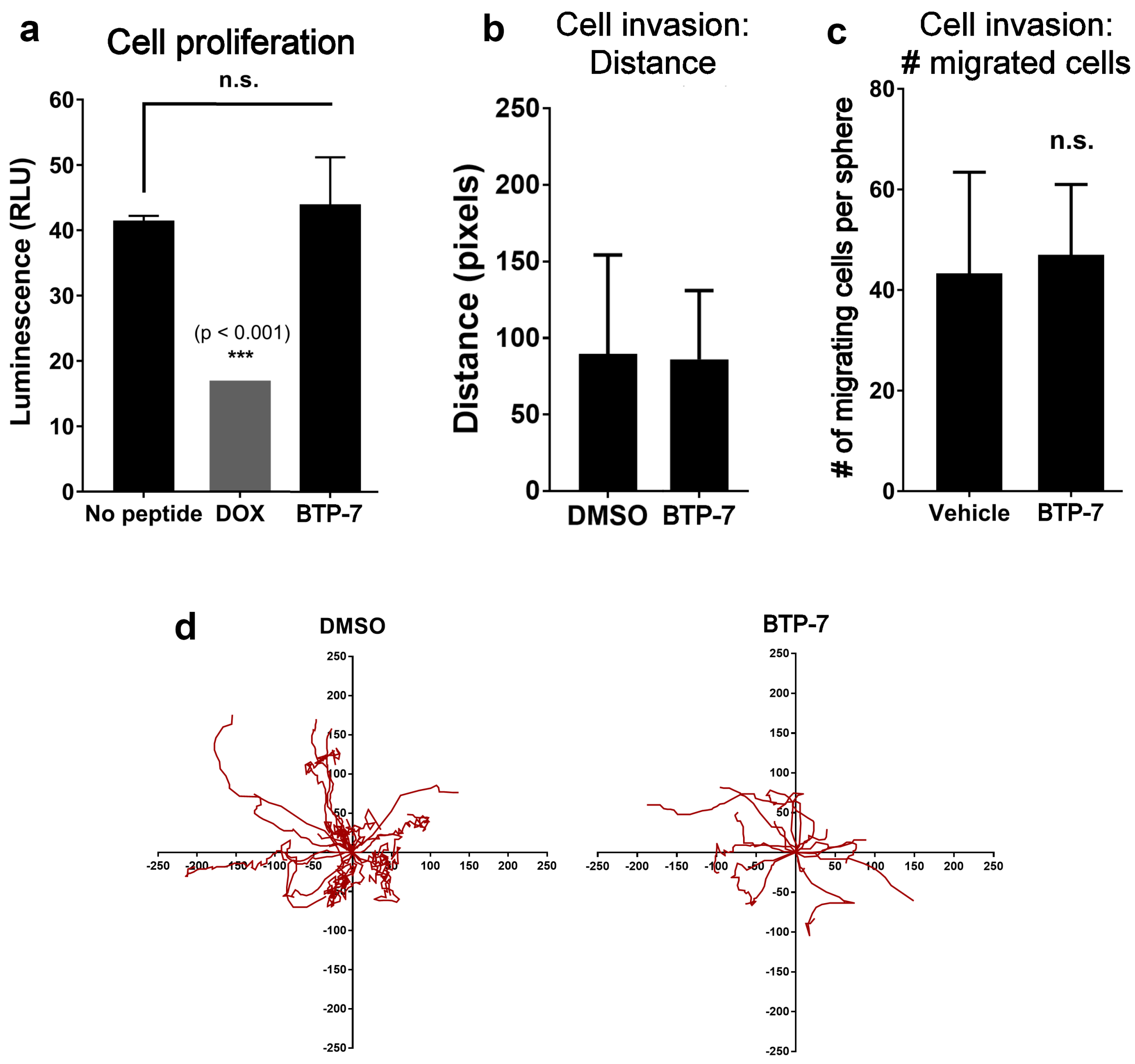


**Fig. S8. Effect of BTP-7 on GBM cell proliferation and invasion**

**a)** Luminescent cell viability (CellTitre Glo) assay of GBM-6 cells (14,000 cells per well) 48 hrs after treatment with BTP-7 (50 µM), vehicle (DMSO) and doxorubicin (DOX, 10 µM) as a positive control. ($n_{well} = 3$). **b, c, d)** Collagen invasion assay of GBM-6 cells treated with BTP-7 (50 µM) or DMSO. GBM-6 neurospheres (n = 3) were embedded in collagen and imaged over 60 hrs. Each cell that has disseminated and migrated from the body of the neurosphere was tracked. The bar graph shows the average distance traveled by each cell, as measured using the ImageJ software. **d)** Cellular tracks of GBM-6 cells (red line) that have migrated from the neurosphere ($n_{sphere} = 3$) over 48 hrs in the presence of BTP-7 (50 µM) or vehicle (DMSO) control.

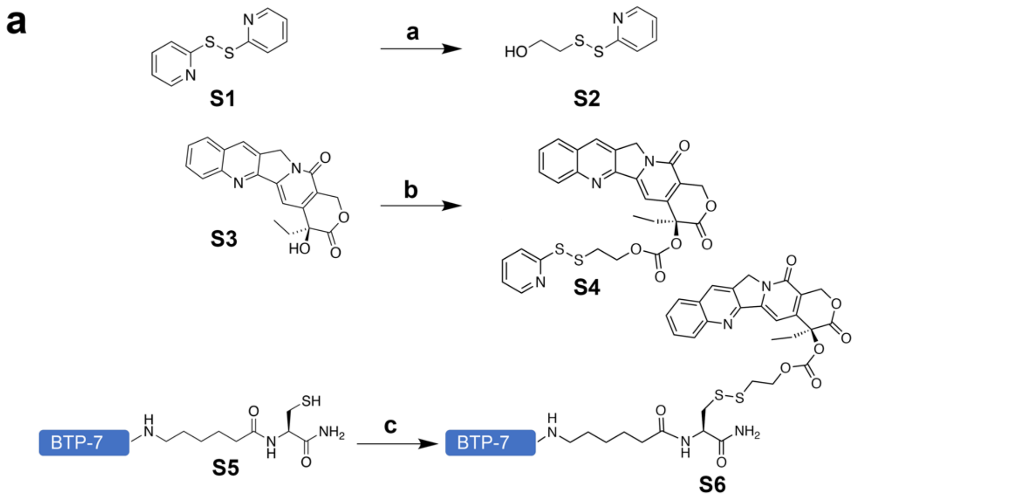


a) **S1** β-mercaptoethanol, DCM, 2h
b) Camptothecin (**S3**), Triphosgene, 4-dimethylaminopyridine, DCM 15min, mixed disulfide **S2**, DCM 1h r.t.
c) **S5,** 0.1M 2-(N-morpholino)ethanesulfonic acid pH 6 in H2O, **S4** in DMSO 1h r.t.

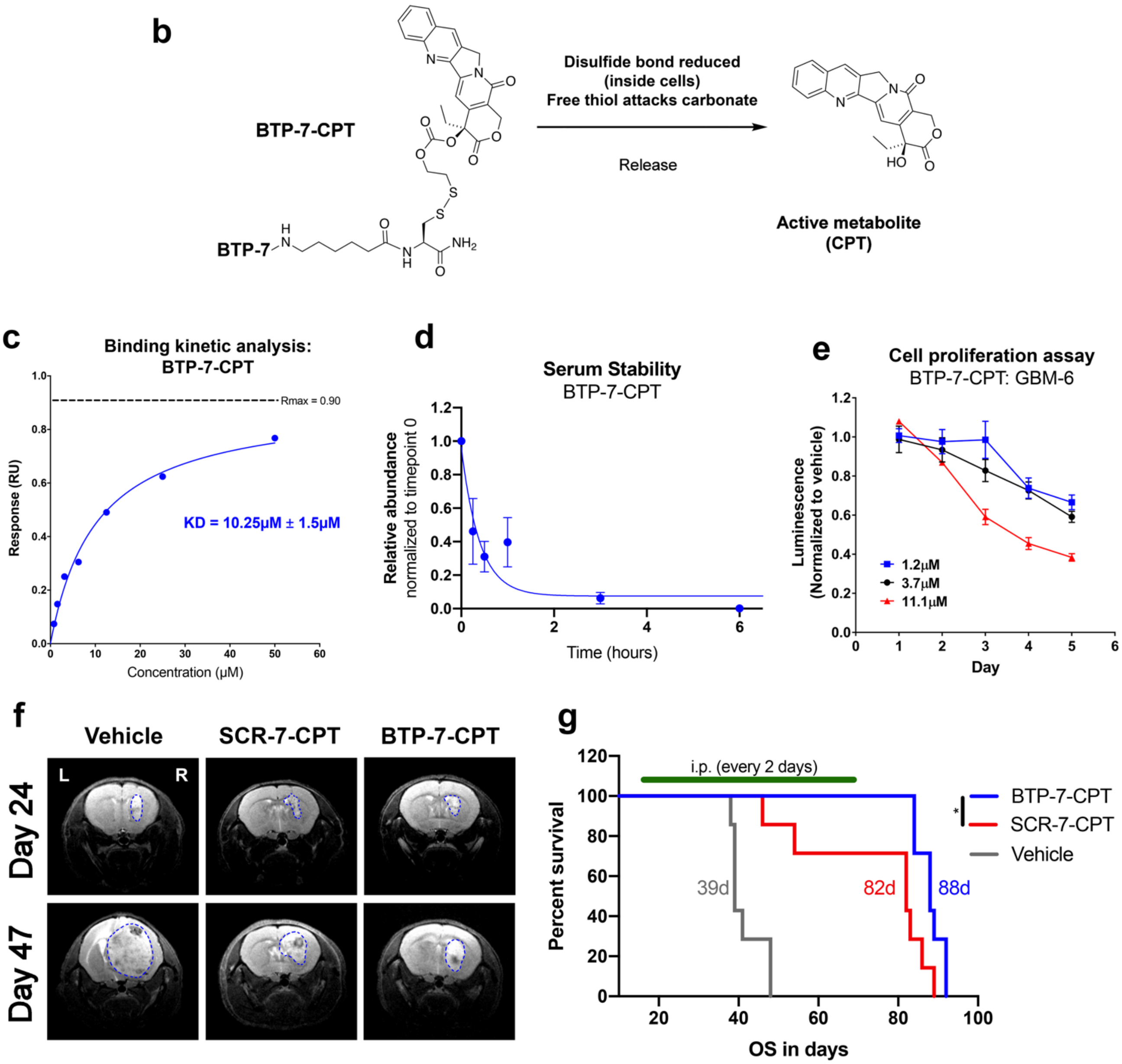

**Fig. S9. Characteristics and efficacy of Camptothecin (CPT) functionalized with BTP-7 in vitro and in vivo**

**a)** Chemical synthesis workflow showing the conjugation of BTP-7 to camptothecin (CPT) (details listed at the end of the SI) and cleaving mechanism upon cell internalization **b)**. **c)** Octet binding kinetic analysis showing the affinity of BTP-7-CPT to dg-Bcan. **d)** Stability profile of BTP-7-CPT in human serum. **e)** Luminescent cell viability (CellTitre Glo) of GBM-6 cells treated with different concentrations of BTP-7-CPT (30,000 cells per well, over 5 days, $n_{well} = 3$). **f)** Representative coronal MRI images of GBM-6 xenograft tumors (delineated with blue dotted lines) established in the right frontal at day 24 post-tumor implantation (before treatment) and at day 47 (after treatment). Both the BTP-7-CPT and Scr-7-CPT groups show reduced tumor burden compared to the control group. **g)** Kaplan-Meier survival plot of second survival study with longer treatment regimen ((i.p. at 10 mg/kg dose every two days for 50 days). A significant difference ($p < 0.05$) is observed between the BTP-7-CPT and Scr-7-CPT group, and ($p < 0.0001$) between the treated groups and vehicle, as determined by the Log-rank (Mantel-Cox) test.

#### Supplementary Notes

**Sequences, synthesis reactions and chromatograms.** Amino acid sequences, as well as LCMS TICs or HPLC chromatograms and mass spectra of all proteins and peptides used in this study are depicted below.

**Peptide: BTP-2-FITC**
Sequence: wrkaftGy-Ahx-K(FITC)

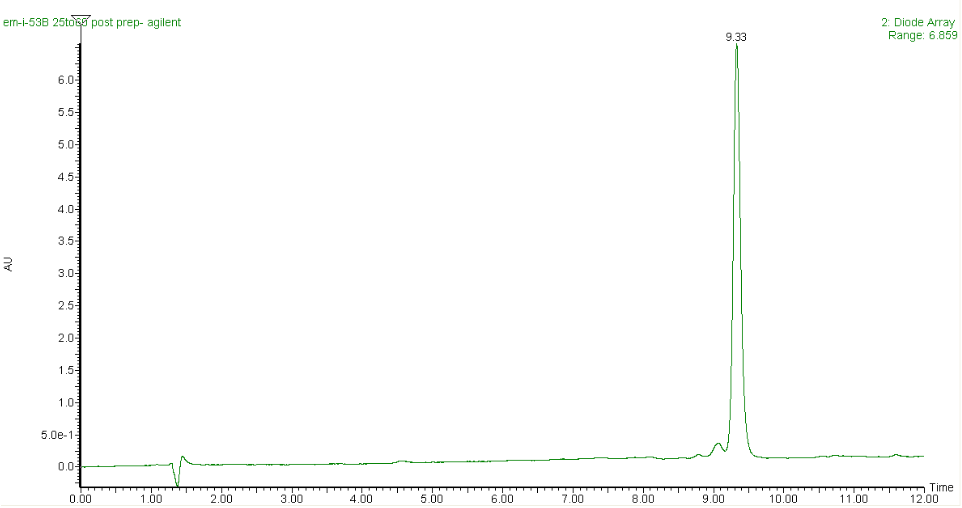


ESI-MS: *m/z* $[M+2H]^{2+}$ Expected 829.4; Observed 829.7

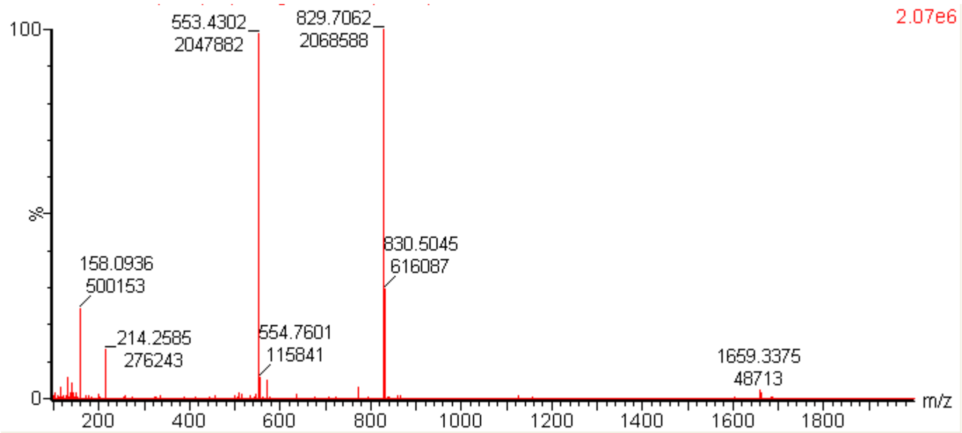

**Peptide: BTP-3-FITC**

Sequence: rrrhdanp-Ahx-K(FITC)

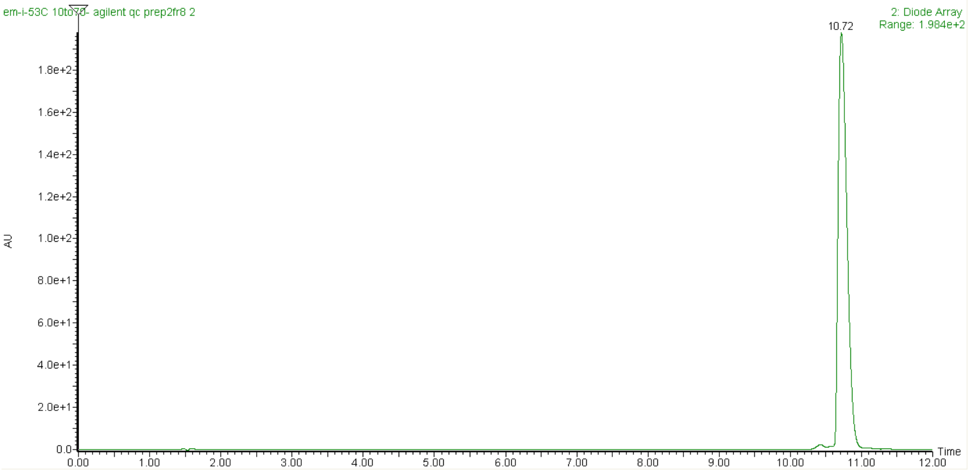


ESI-MS: *m/z* $[M+3H]^{3+}$ Expected 550.9; Observed 551.0

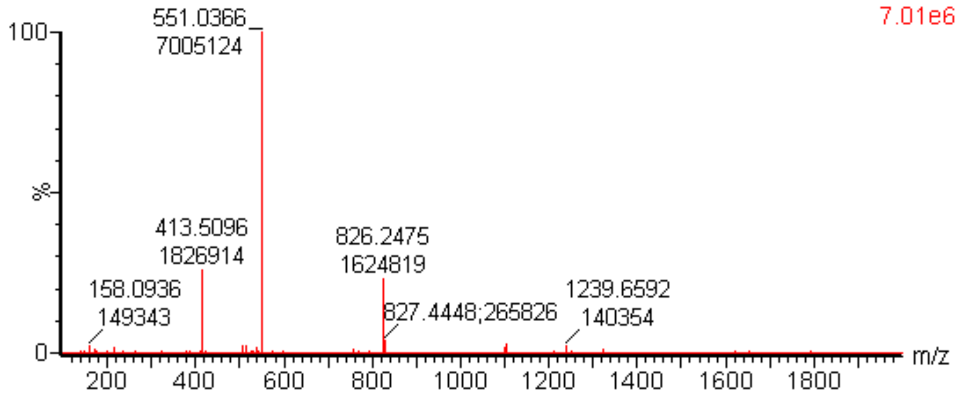

**Peptide: BTP-5-FITC**
Sequence: nkhvfrhw-Ahx-K(FITC)

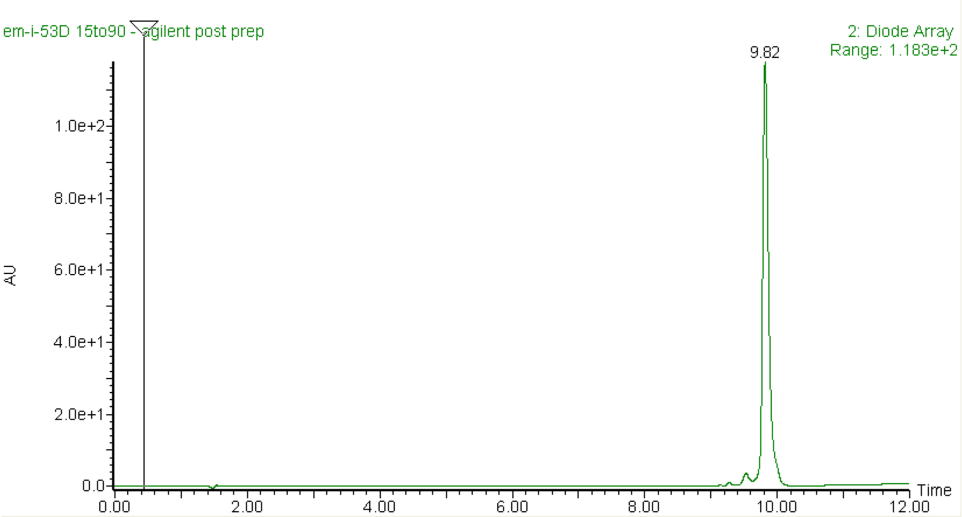


ESI-MS: *m/z* $[M+3H]^{3+}$ Expected 584.9; Observed 585.2

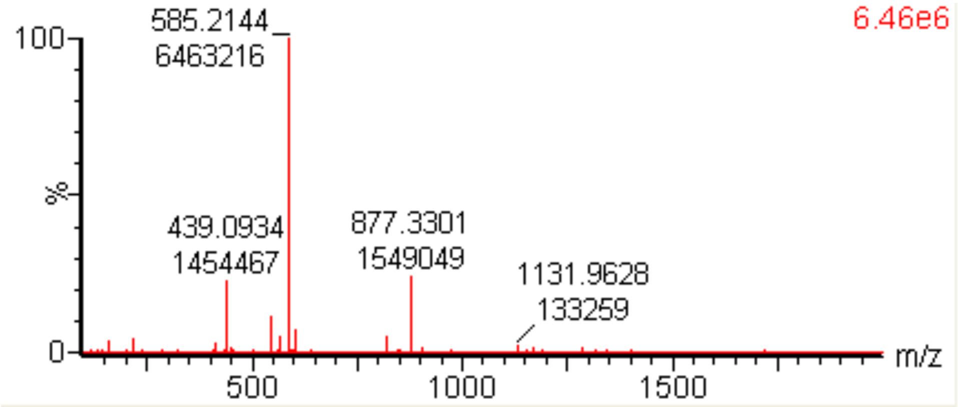

**Peptide: BTP-7-FITC**
Sequence: tkwGhvnk-Ahx-K(FITC)

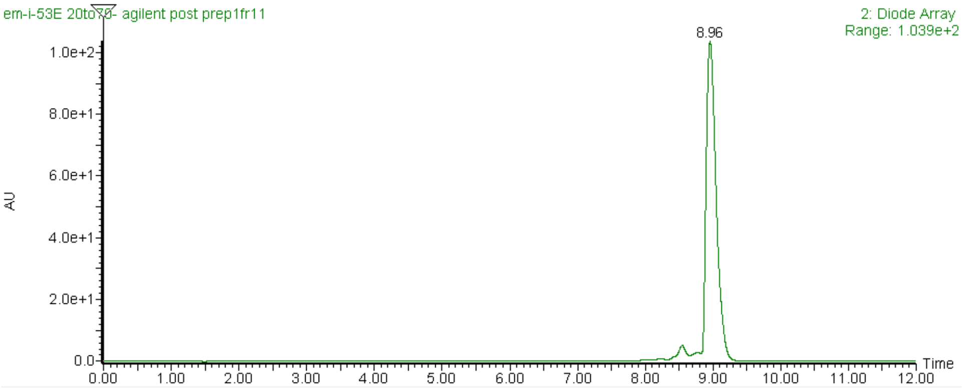


ESI-MS: *m/z* $[M+3H]^{3+}$ Expected 533.6; Observed 533.8

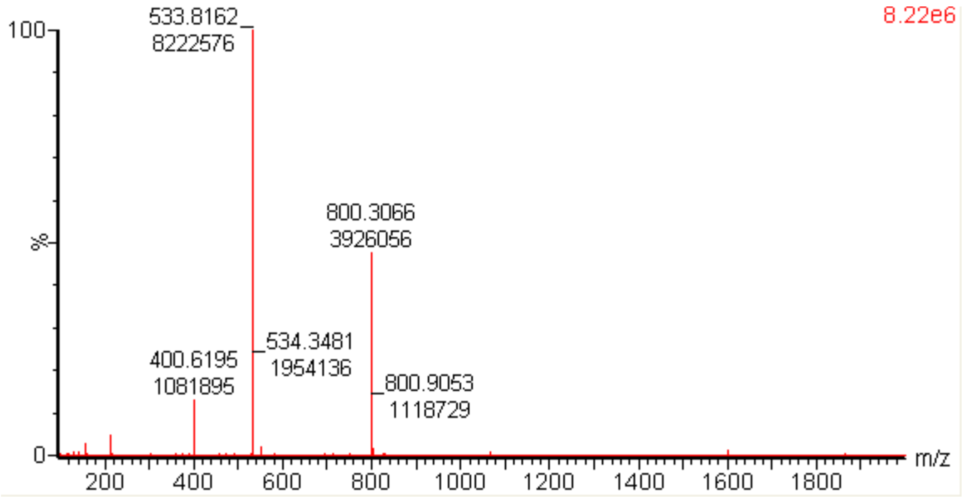

**Peptide: BTP-8-FITC**

Sequence: tirklvrh-Ahx-K(FITC)

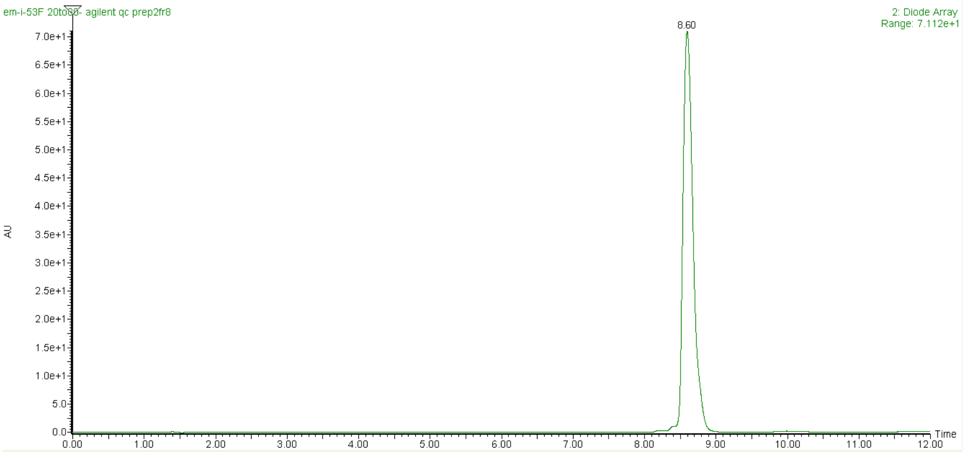


ESI-MS: *m/z* $[M+3H]^{3+}$ Expected 551.3; Observed 551.5

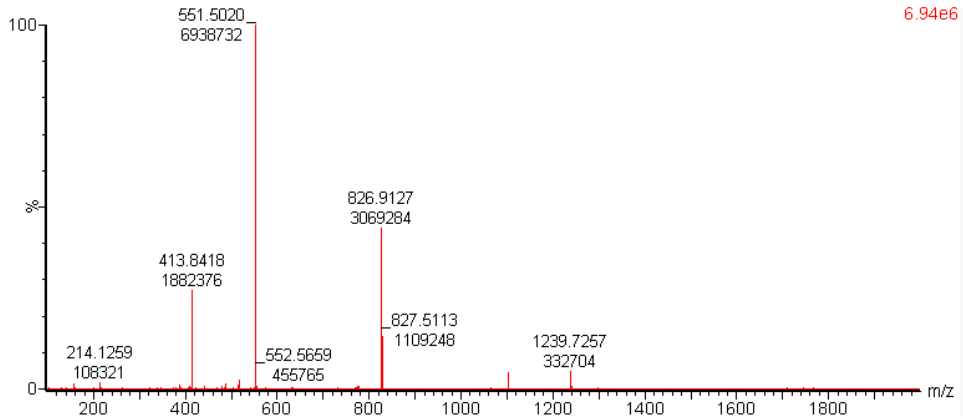

**Peptide: BTP-9-FITC**

Sequence: adrrqrai-Ahx-K(FITC)

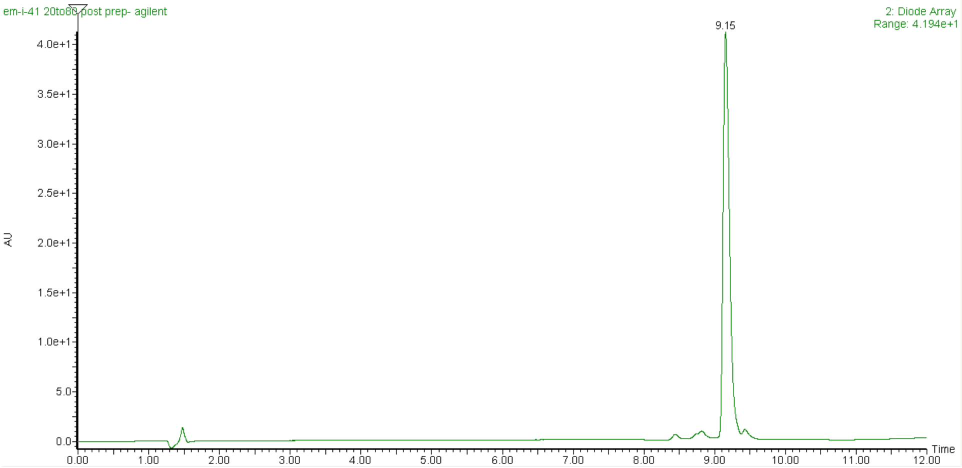


ESI-MS: *m/z* $[M+3H]^{3+}$ Expected 538.9; Observed 539.1

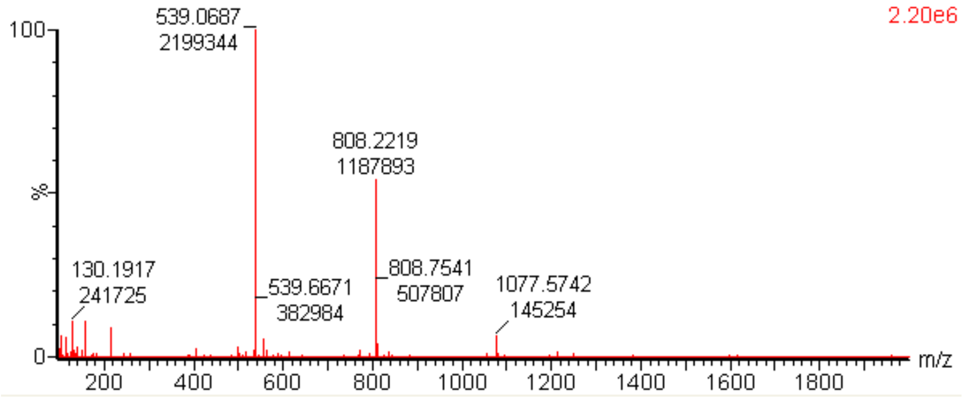

**Peptide: BTP-10-FITC**
Sequence: shwavnrf-Ahx-K(FITC)

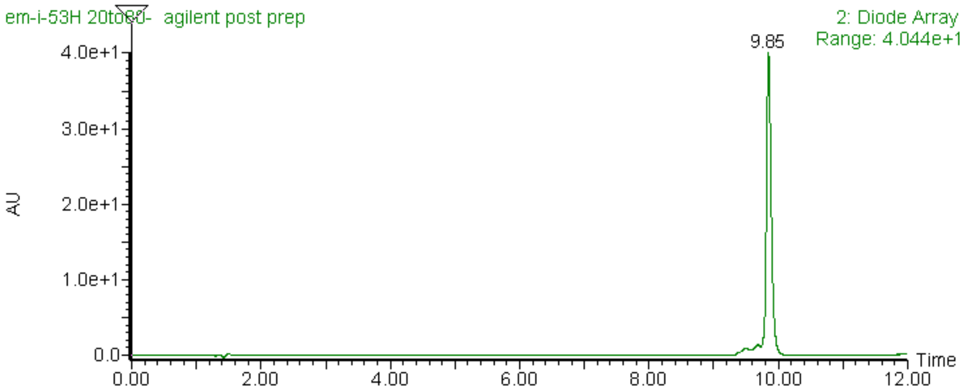


ESI-MS: *m/z* $[M+3H]^{3+}$ Expected 549.3; Observed 549.4

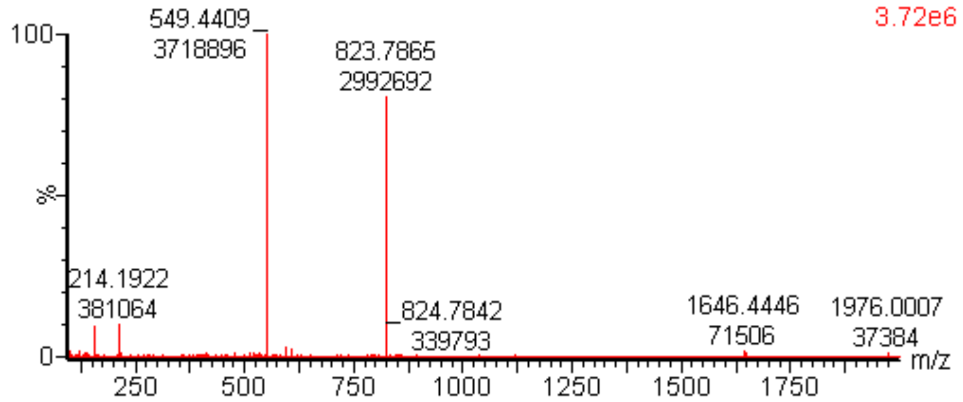

**Peptide: BTP-2-Cy5.5**
Sequence: wrkaftGy-Ahx-G(Cy5.5)

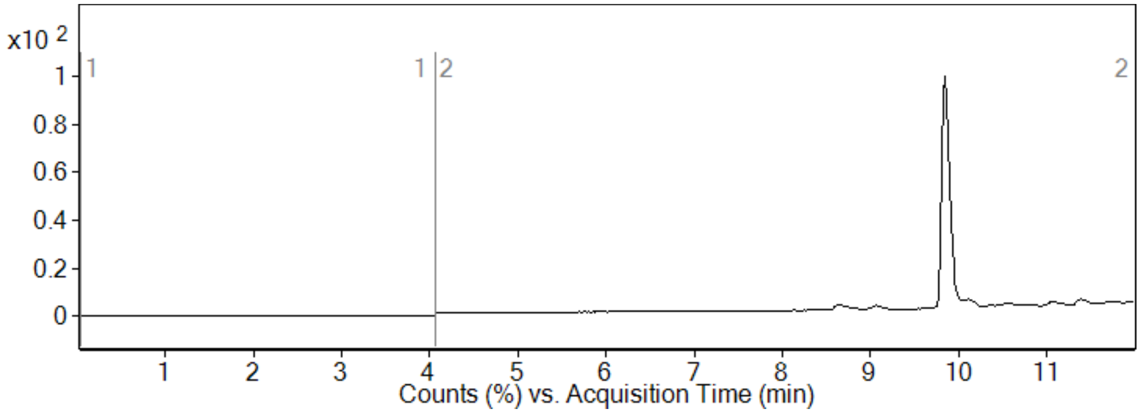


ESI-MS: *m/z* $[M+3H]^{3+}$ Expected 634.3; Observed 634.4

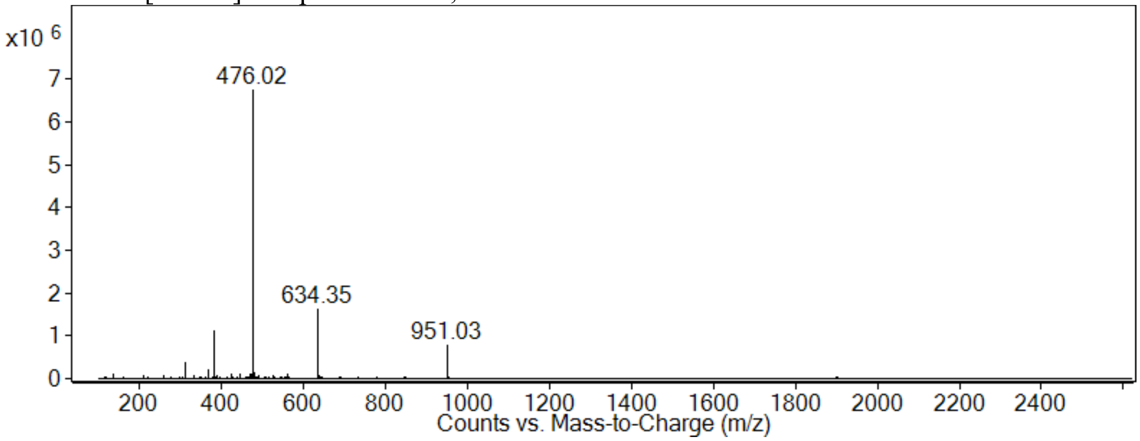

**Peptide: BTP-7-Cy5.5**
Sequence: tkwGhvnk-Ahx-G(Cy5.5)

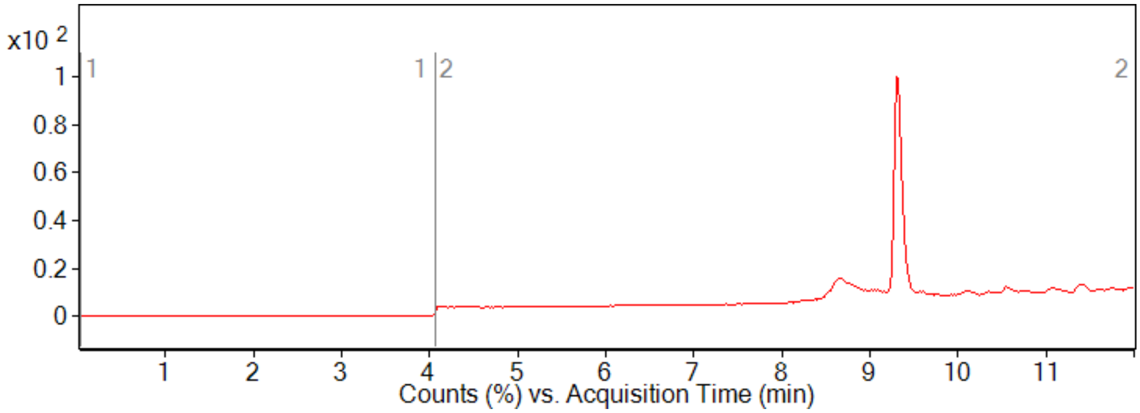


ESI-MS: *m/z* $[M+3H]^{3+}$ Expected 614.6; Observed 614.7

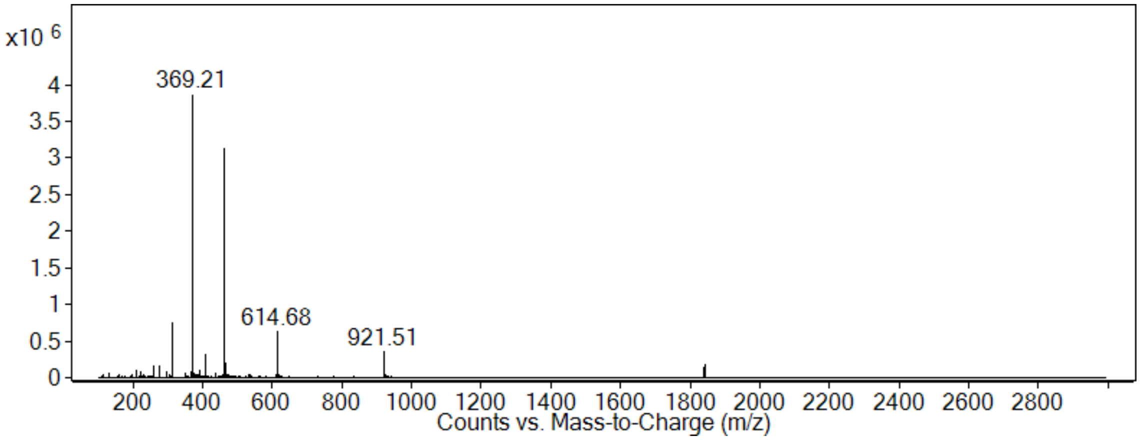

**Peptide: BTP-8-Cy5.5**
Sequence: tirklvrh-Ahx-G(Cy5.5)

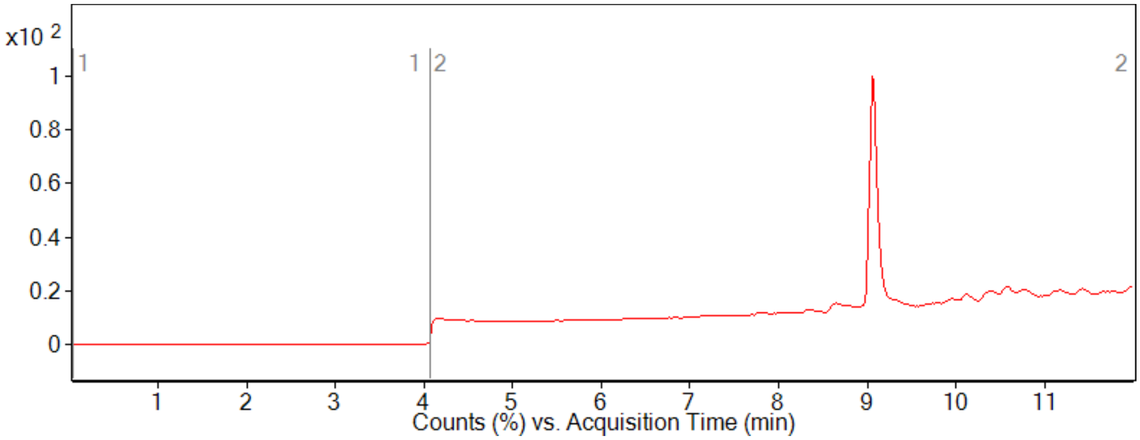


ESI-MS: *m/z* $[M+3H]^{3+}$ Expected 632.3; Observed 632.4

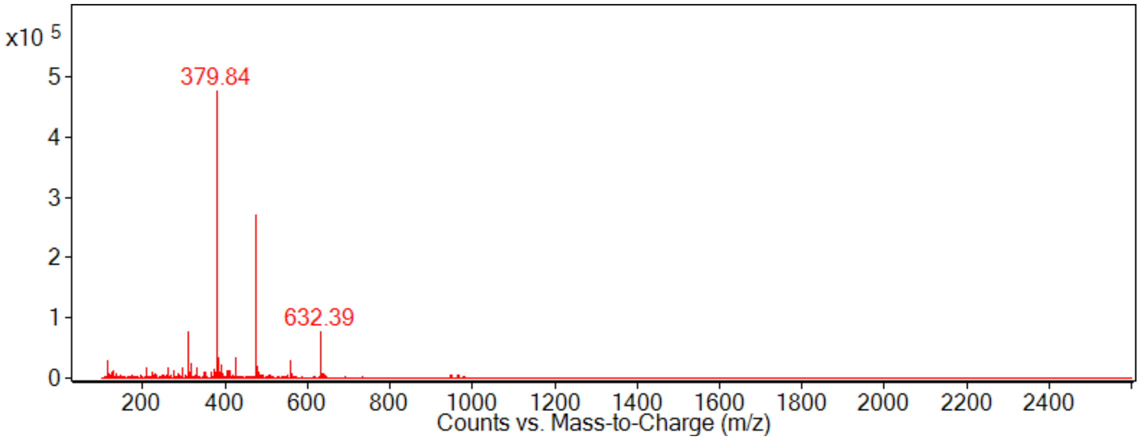

**Peptide: BTP-9-Cy5.5**
Sequence: adrrqrai-Ahx-G(Cy5.5)

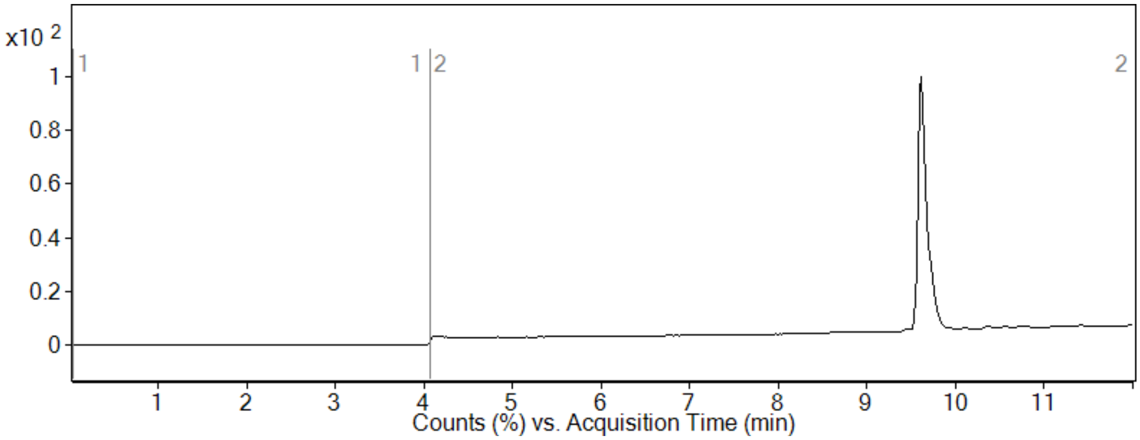


ESI-MS: *m/z* $[M+3H]^{3+}$ Expected 620.0; Observed 620.0

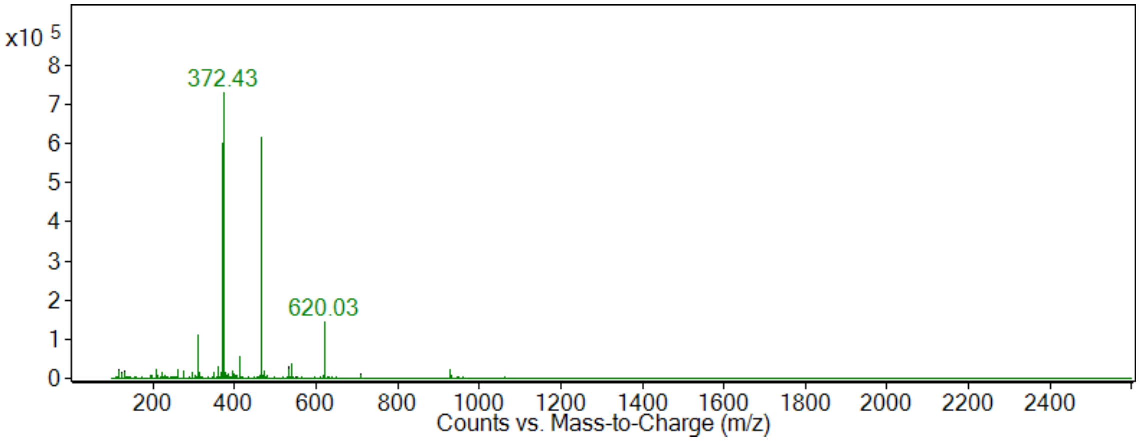

**Peptide: BTP-10-Cy5.5**
Sequence: shwavnrf-Ahx-G(Cy5.5)

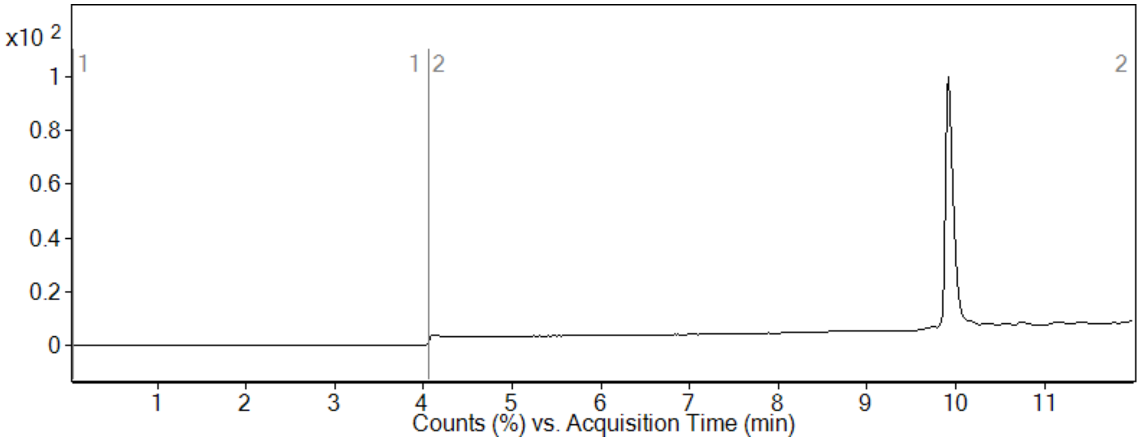


ESI-MS: *m/z* $[M+3H]^{3+}$ Expected 630.3; Observed 630.4

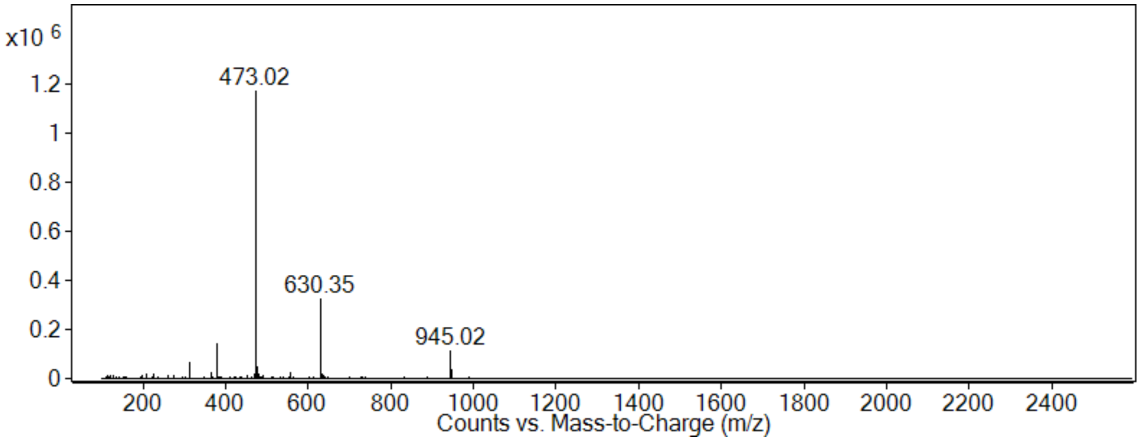

**Peptide: Scr-7-Cy5.5**

Sequence: hnGkwkvt-Ahx-Azidolysine(Cy5.5)
Mass expected: 1855.0 Da
Mass observed: 1855.4 Da

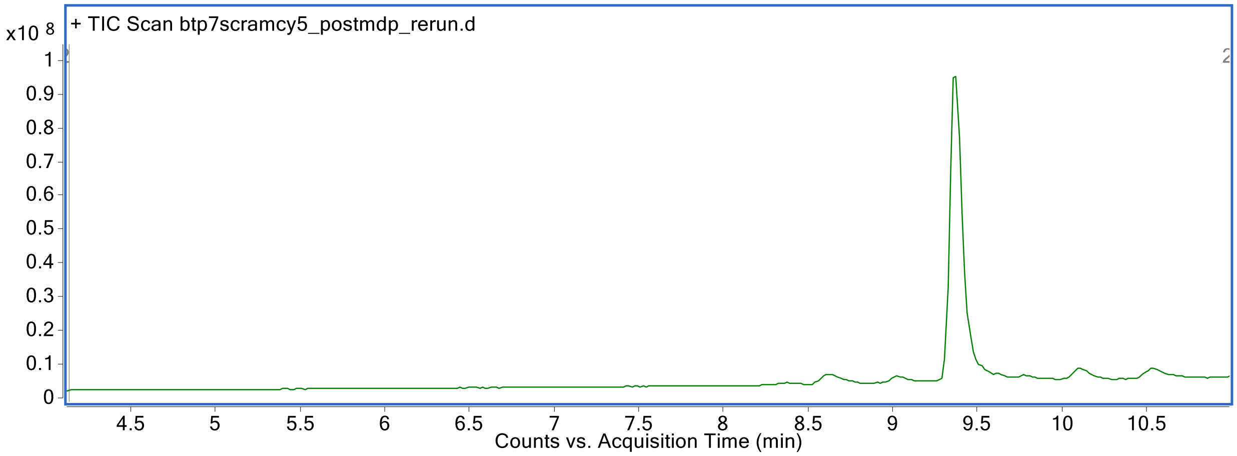


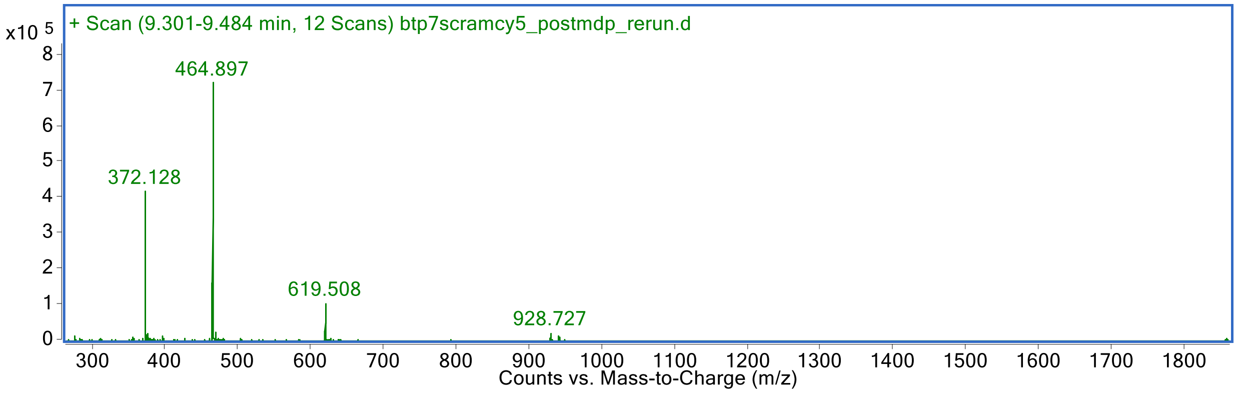

**Peptide: BTP-7 (precursor before Cy5.5 conjugation)**

Sequence: tkwGhvnk-Ahx-propargylglycine
Mass expected: 1176.6 Da
Mass observed: 1176.7 Da

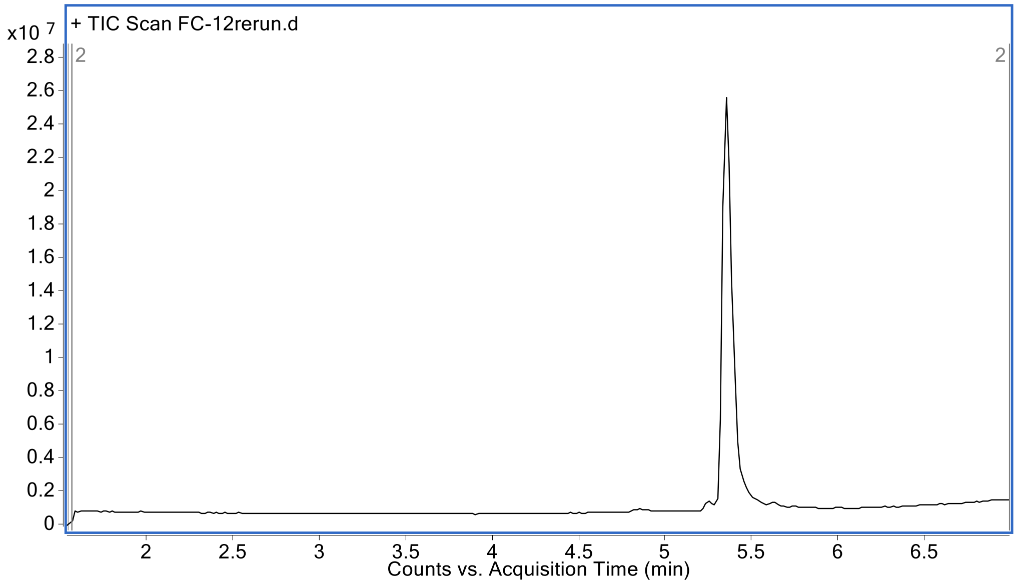


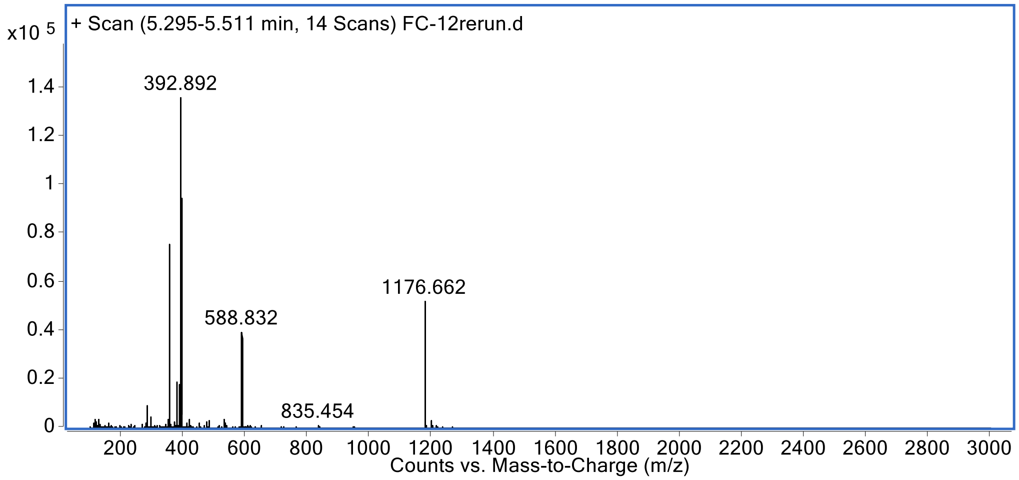

**Peptide: BTP-7-Ahx**

Sequence: tkwGhvnk-Ahx
Mass expected: 1080.6 Da
Mass observed: 1080.6 Da

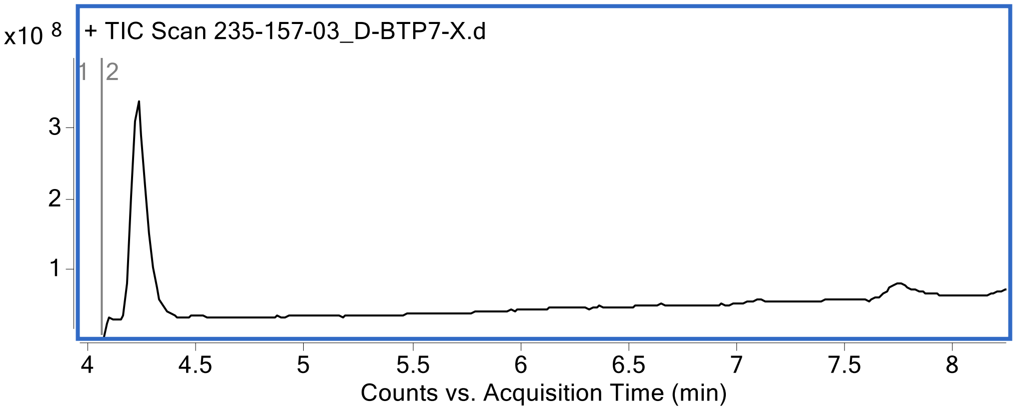


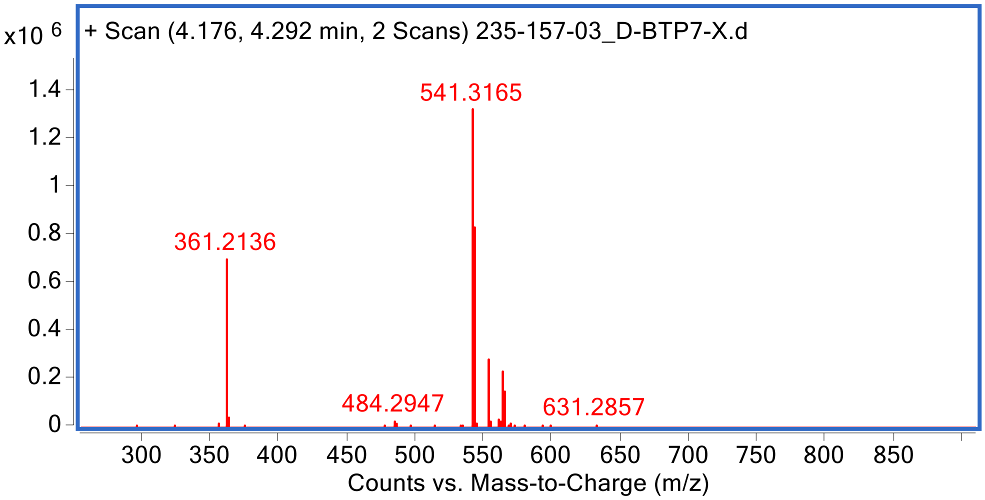

**Peptide: L-BTP-7-Ahx**

Sequence: TKWGHVNK-Ahx
Mass expected: 1080.6 Da
Mass observed: 1080.6 Da

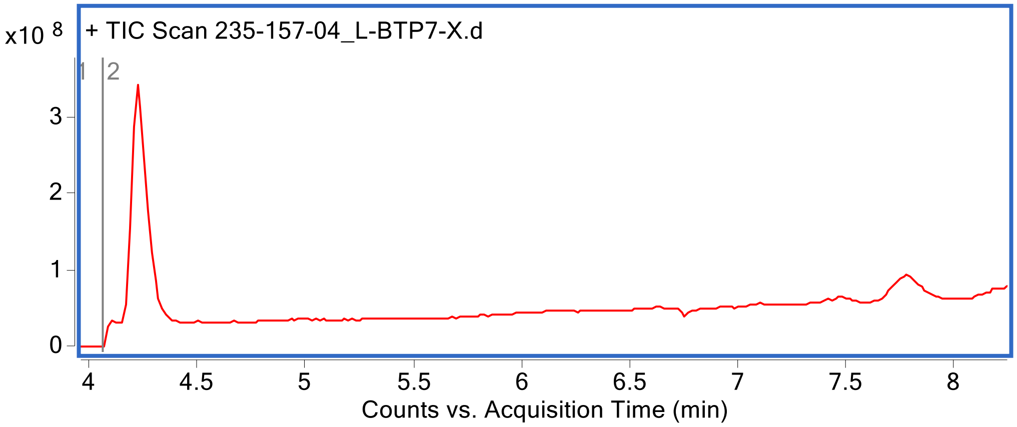


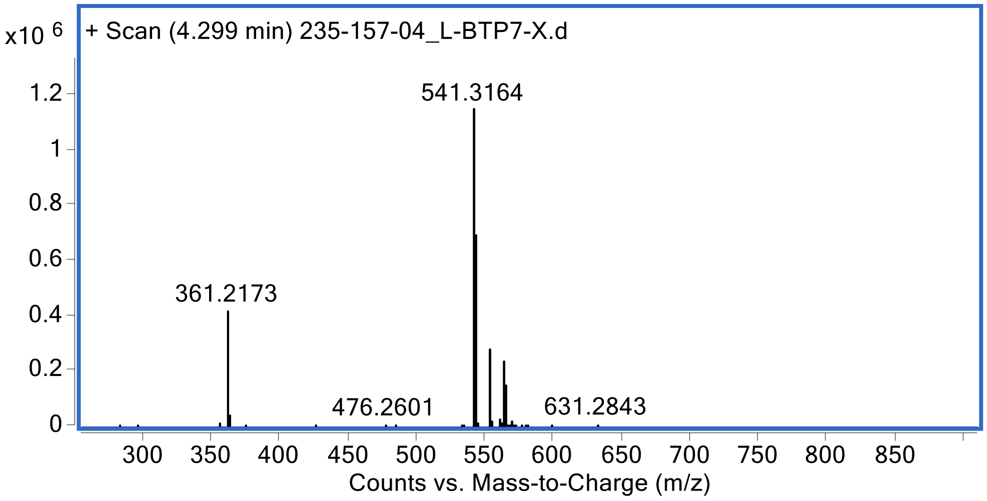

**Peptide: BTP-7-biotin-crosslinker**

Sequence: tkwGhvnk-Ahx-K(diazirine)-Ahx-K(biotin)
Mass expected: **1786.0188** Da
Mass observed: **1786.0158** Da

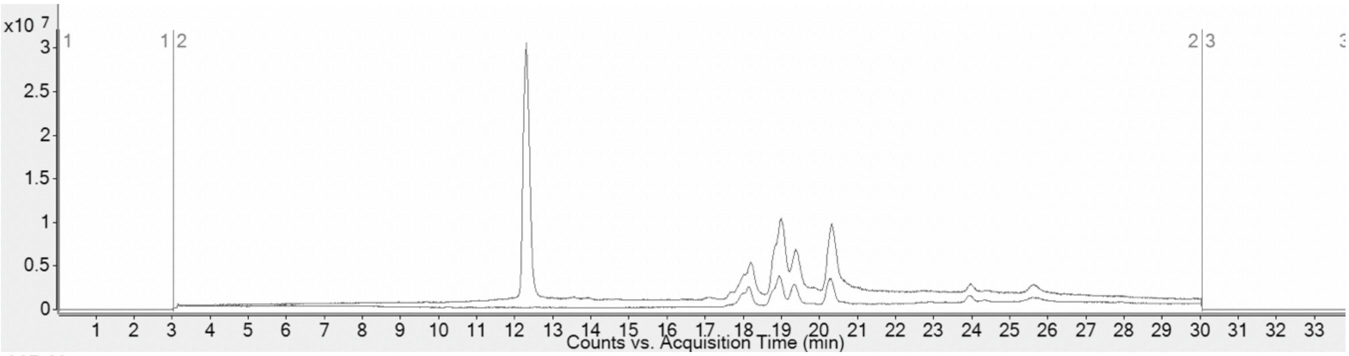


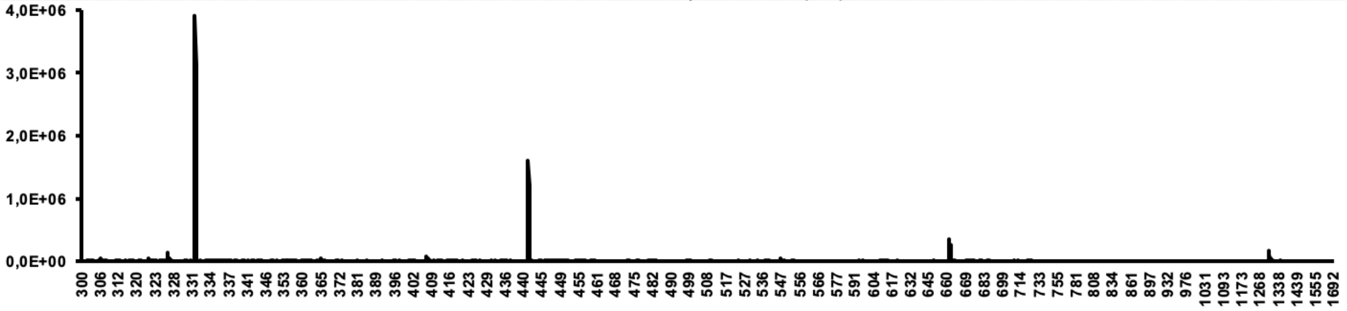


LCMS (Jupiter C4 column, 1-61% MeCN, overlayed with blank)
BTP-7-diazirine
Chemical Formula (uncharged peptide): $C_{83}H_{135}N_{25}O_{17}S$
Exact mass: 1786.0188
Molecular weight: 1787.2110
Molecular weight of TFA salt: 2243.3038

**MALDI mass spectrometry of magnetic screening particles**

**Peptide: dg-Bcan**

| Predicted Fragmentation Pattern | | | | | | |
|---|---|---|---|---|---|---|
| Seq | # | b: Δ Error | b | y | y: Δ Error | 1 |
| V | 1 | 0.669 | 455.232 | --- | --- | 14 |
| H | 2 | 0.287 | 592.291 | 1431.723 | 3.316 | 13 |
| G | 3 | 1.233 | 649.312 | 1294.664 | 3.347 | 12 |
| P | 4 | 2.342 | 746.365 | 1237.642 | 3.682 | 11 |
| P | 5 | --- | 843.418 | 1140.59 | 2.523 | 10 |
| T | 6 | 17.964 | 944.465 | 1043.537 | 2.669 | 9 |
| E | 7 | 3.135 | 1073.508 | 942.489 | 2.188 | 8 |
| T | 8 | 3.221 | 1174.556 | 813.446 | 1.398 | 7 |
| L | 9 | 4.203 | 1287.64 | 712.399 | 1.183 | 6 |
| P | 10 | --- | 1384.692 | 599.315 | -0.397 | 5 |
| T | 11 | --- | 1485.74 | 502.262 | -0.05 | 4 |
| P | 12 | --- | 1582.793 | 401.214 | -0.266 | 3 |
| R | 13 | --- | 1738.894 | 304.162 | 0.348 | 2 |
| E | 14 | --- | --- | 148.06 | --- | 1 |

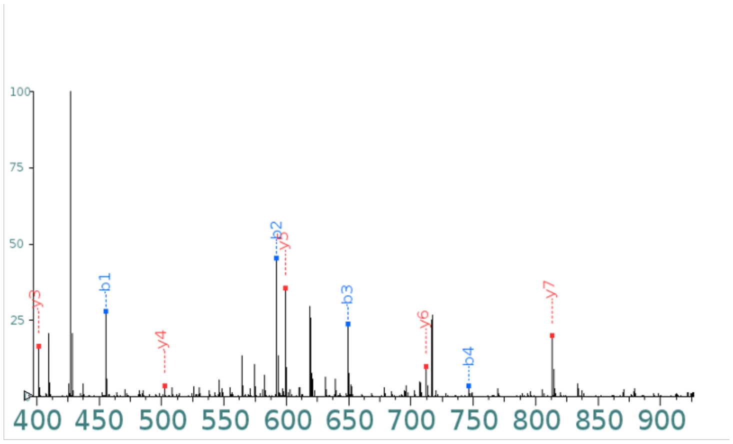


| Predicted Fragmentation Pattern | | | | | | |
|---|---|---|---|---|---|---|
| Seq | # | b: Δ Error | b | y | y: Δ Error | 1 |
| V | 1 | 1.005 | 455.232 | --- | --- | 14 |
| H | 2 | 0.596 | 592.291 | 1431.723 | 2.634 | 13 |
| G | 3 | 1.327 | 649.312 | 1294.664 | 2.97 | 12 |
| P | 4 | 1.851 | 746.365 | 1237.642 | 3.485 | 11 |
| P | 5 | --- | 843.418 | 1140.59 | 2.095 | 10 |
| T | 6 | 1.873 | 944.465 | 1043.537 | 1.616 | 9 |
| E | 7 | 3.022 | 1073.508 | 942.489 | 1.735 | 8 |
| T | 8 | 2.493 | 1174.556 | 813.446 | 0.573 | 7 |
| L | 9 | 4.488 | 1287.64 | 712.399 | 0.925 | 6 |
| P | 10 | --- | 1384.692 | 599.315 | -0.498 | 5 |
| T | 11 | 4.479 | 1485.74 | 502.262 | -0.111 | 4 |
| P | 12 | --- | 1582.793 | 401.214 | -0.19 | 3 |
| R | 13 | --- | 1738.894 | 304.162 | -2.561 | 2 |
| E | 14 | --- | --- | 148.06 | --- | 1 |

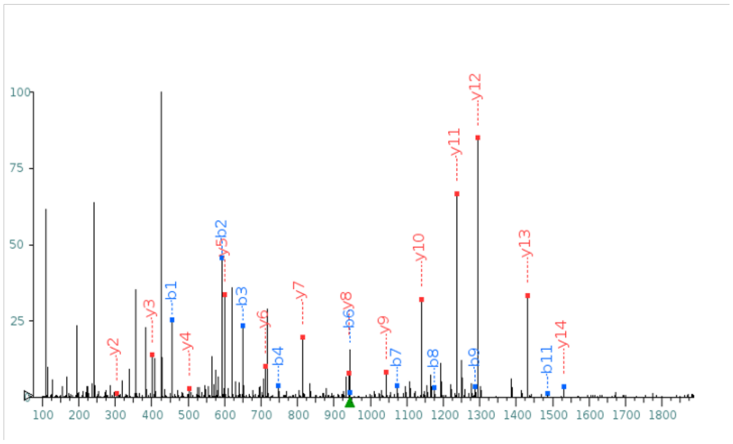

**Peptide: Scramble-dg-Bcan**

| Predicted Fragmentation Pattern | | | | | | |
|---|---|---|---|---|---|---|
| Seq | # | b: Δ Error | b | y | y: Δ Error | 1 |
| T | 1 | -0.171 | 457.211 | --- | --- | 14 |
| R | 2 | -0.187 | 613.312 | 1429.743 | --- | 13 |
| E | 3 | 1.667 | 742.355 | 1273.642 | --- | 12 |
| L | 4 | 1.996 | 855.439 | 1144.6 | --- | 11 |
| T | 5 | 1.391 | 956.486 | 1031.516 | --- | 10 |
| P | 6 | --- | 1053.539 | 930.468 | -3.15 | 9 |
| G | 7 | 1.795 | 1110.561 | 833.415 | --- | 8 |
| V | 8 | 1.805 | 1209.629 | 776.394 | --- | 7 |
| E | 9 | 2.914 | 1338.672 | 677.325 | --- | 6 |
| T | 10 | 2.661 | 1439.719 | 548.283 | 0.541 | 5 |
| H | 11 | 1.686 | 1576.778 | 447.235 | -1.361 | 4 |
| P | 12 | --- | 1673.831 | 310.176 | -2.315 | 3 |
| P | 13 | --- | 1770.884 | 213.123 | -1.656 | 2 |
| P | 14 | --- | --- | 116.071 | -1.734 | 1 |

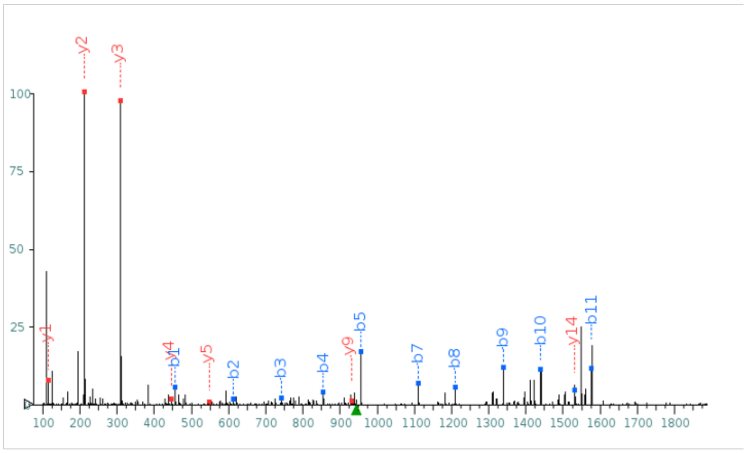


| Predicted Fragmentation Pattern | | | | | | |
|---|---|---|---|---|---|---|
| Seq | # | b: Δ Error | b | y | y: Δ Error | 1 |
| T | 1 | -0.905 | 457.211 | --- | --- | 14 |
| R | 2 | 0.111 | 613.312 | 1429.743 | --- | 13 |
| E | 3 | -1.457 | 742.355 | 1273.642 | --- | 12 |
| L | 4 | 0.426 | 855.439 | 1144.6 | --- | 11 |
| T | 5 | 0.881 | 956.486 | 1031.516 | --- | 10 |
| P | 6 | --- | 1053.539 | 930.468 | -0.526 | 9 |
| G | 7 | 0.916 | 1110.561 | 833.415 | --- | 8 |
| V | 8 | -0.012 | 1209.629 | 776.394 | --- | 7 |
| E | 9 | 2.276 | 1338.672 | 677.325 | --- | 6 |
| T | 10 | 1.558 | 1439.719 | 548.283 | -3.801 | 5 |
| H | 11 | 1.144 | 1576.778 | 447.235 | -2.18 | 4 |
| P | 12 | --- | 1673.831 | 310.176 | -2.708 | 3 |
| P | 13 | --- | 1770.884 | 213.123 | -2.085 | 2 |
| P | 14 | --- | --- | 116.071 | -2.129 | 1 |

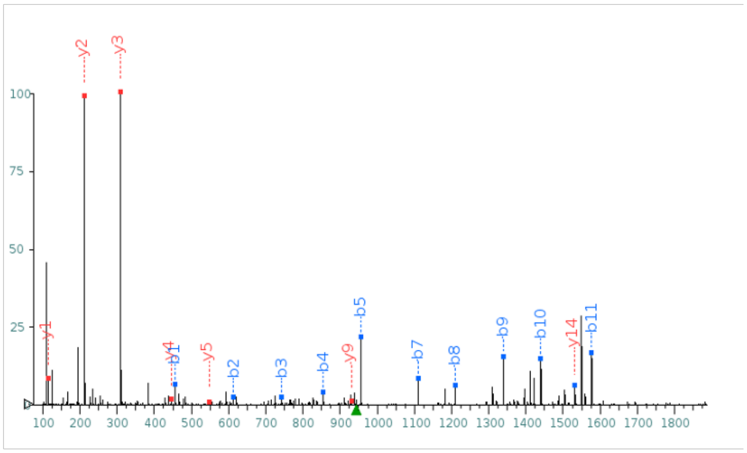

**Synthesis of the Campthotecin-linker and conjugation to BTP-7**

**2-(pyridin-2-yldisulfaneyl)ethan-1-ol (S2)[1]**

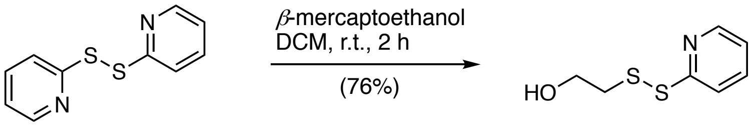


**S1** **S2**

2-Mercaptoethanol (500 µL, 7.10 mmol, 1 eq.) and 2,2'-dipyridyl disulfide (**S1**, 4.70 g, 21.4 mmol, 3 eq.) were dissolved in DCM (20 mL) and stirred at room temperature for 3 hrs. Afterwards, the reaction mixture was concentrated under reduced pressure and flash column chromatography [hexanes/EtOAc, 10:1 to 2:1] afforded mixed disulfide **S2** (1.01 g, 5.39 mmol, 76%) as a yellow solid.

**R*f*** = 0.45 [hexanes/ethyl acetate, 1:1].

**$^1$H NMR**(500 MHz, DMSO-*d*$_6$) δ 8.50 – 8.42 (m, 1H), 7.88 – 7.75 (m, 2H), 7.24 (ddd, *J* = 6.6, 4.8, 2.1 Hz, 1H), 4.99 (t, *J* = 5.5 Hz, 1H), 3.62 (q, *J* = 6.1 Hz, 2H), 2.92 (t, *J* = 6.3 Hz, 2H) ppm.

**$^{13}$C NMR** (500 MHz, DMSO-*d*$_6$) δ 159.5, 149.5, 137.8, 121.1, 119.3, 59.1, 41.2 ppm.

**FTIR** (thin film): $\tilde{\nu}$ = 3288 (br), 2920 (w), 2863 (w), 1574 (m), 1415 (s), 1043 (m), 754 (s) cm$^{-1}$.

**HRMS** (ESI): calcd. for $C_7H_{10}NOS_2^+$: 188.0198 [M+H]$^+$
found: 188.0269 [M+H]$^+$.

**2-pyridinyldithioethyl carbonate Camptothecin (S4)[1]**

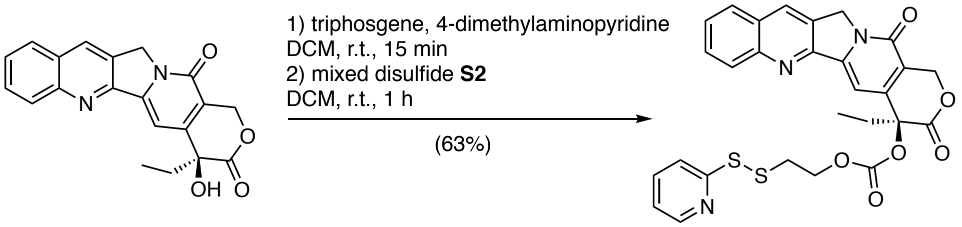


**S3** **S4**

Camptothecin (**S3,** 250 mg, 0.718 mmol, 1 eq.), triphosgene (82.6 mg, 0.278 mmol, 0.387 eq.) and 4-dimethylaminopyridine (459 mg, 3.75 mmol, 5.21 eq.) were combined in dry DCM (10 mL), and after 15 min, mixed disulfide **S2** was added (148 mg, 0.790 mmol, 1.10 eq.) and the reaction was stirred at room temperature for 4 hrs. Flash column chromatography [DCM:Acetone, 20:1 to 4:1] afforded **S4** (254 mg, 0.452 mmol, 63%).

$\mathbf{R}_f$ = 0.27 [hexanes/ethyl acetate, 1:3].

$^{1}$**H NMR** (500 MHz, DMSO-$d_6$) δ 8.68 (s, 1H), 8.39 (ddd, $J$ = 4.8, 1.9, 0.9 Hz, 1H), 8.18 – 8.09 (m, 2H), 7.85 (ddd, $J$ = 8.5, 6.8, 1.5 Hz, 1H), 7.81 – 7.71 (m, 1H), 7.71 – 7.63 (m, 2H), 7.15 (ddd, $J$ = 7.4, 4.8, 1.1 Hz, 1H), 7.09 (s, 1H), 5.52 (d, $J$ = 2.0 Hz, 2H), 5.29 (s, 2H), 4.33 (t, $J$ = 6.0 Hz, 2H), 3.20 – 3.08 (m, 2H), 2.25 – 2.13 (m, 2H), 0.92 (t, $J$ = 7.4 Hz, 3H) ppm.

$^{13}$**C NMR** (500 MHz, DMSO-$d_6$) δ 167.1, 158.6, 156.5, 152.7, 152.2, 149.6, 147.9, 146.3, 144.7, 137.7, 131.6, 130.4, 129.8, 129.0, 128.5, 128.0, 127.8, 121.3, 119.4, 119.2, 94.4, 77.9, 66.5 66.8, 50.3, 36.8, 30.3, 7.6 ppm**.**

**FTIR** (thin film): $\tilde{\nu}$ = 2992.83 (w), 1748.22 (m), 1667.56 (m), 1748.22 (m), 1667.56 (m),1253.34 (m), 666.08 (s) $cm^{-1}$.

| | | |
|---|---|---|
| **HRMS** (ESI): | calcd. for $C_{28}H_{23}N_3O_6S_2^+$: | 562.1101 $[M+H]^+$; |
| | found: | 562.1104 $[M+H]^+$. |

**BTP-7-Camptothecin (S6)**

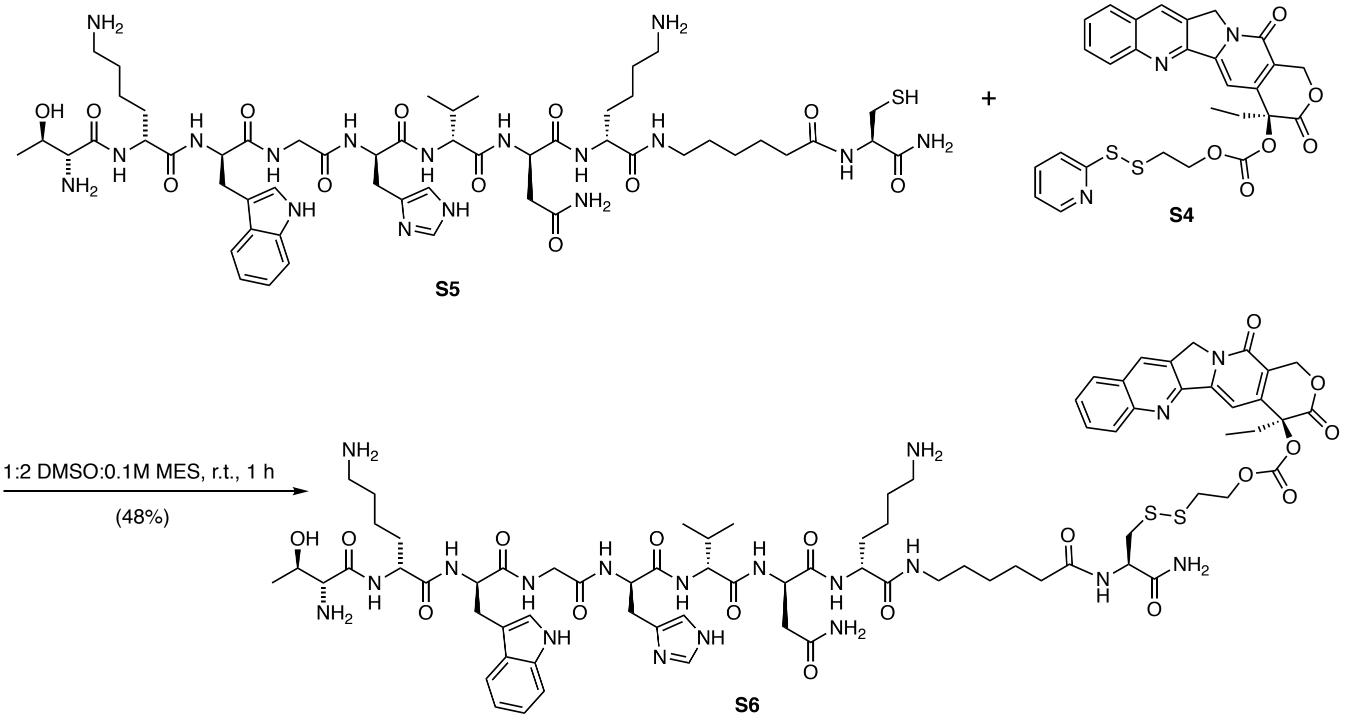


The pyridyldithiol arm of **S4** allows for conjugation to free thiols via disulfide exchange, enabling **S4** to be attached to BTP-7 with a C-terminal cysteine **S5**. To perform this conjugation, the peptide (**S5**, 50 mg, 32 μmol, 1.9 eq.) in 2-($N$-morpholino)ethanesulfonic acid (MES, 0.1 M, pH 6, 14 mL) was combined with **S4**

(9.5 mg, 17 µmol, 1 eq.) in DMSO (7 mL) and stirred at room temperature for 1 hr. The reaction was quenched with trifluoroacetic acid (0.20 mL, 2.6 mmol, 150 eq.). The camptothecin-BTP-7 (**S6**) conjugate was purified by RP-high-performance liquid chromatography (RP-HPLC) in 100 mM ammonium acetate buffer (pH = 5) and an acetonitrile gradient (15–30% over 75 min) (15 mg, 8.1 µmol, 48%). Purity of each RP-HPLC fraction was confirmed by LC-MS.

**HRMS** (LC-MS): calcd. for $C_{76}H_{103}N_{19}O_{18}S_2^{+}$: 817.8661 $[M+2H]^{2+}$
found: 817.8666 $[M+2H]^{2+}$.

**Scrambled BTP-7-Camptothecin (S8)**

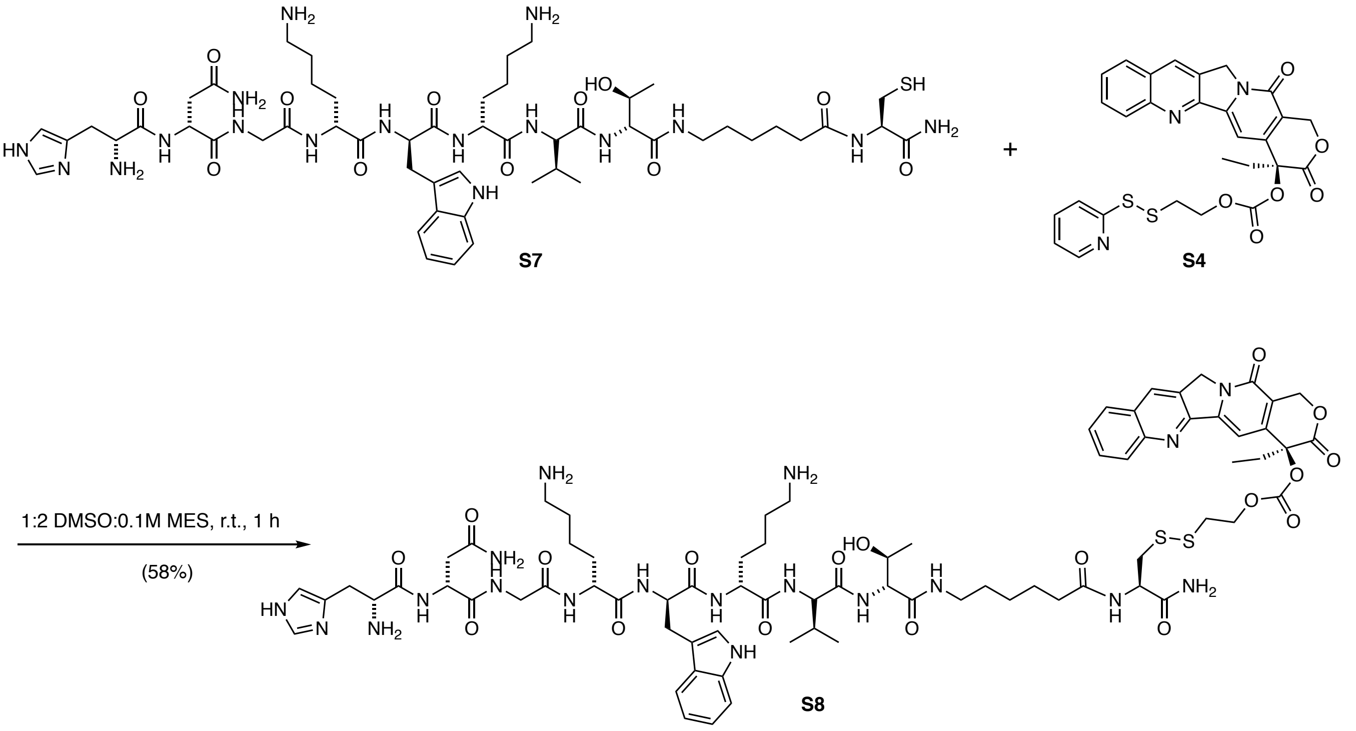


The conjugation and purification protocols given for **S6** were used to conjugate the scrambled BTP-7 peptide (**S7**) to camptothecin to synthesize the camptothecin-scramble (**S8**) (18 mg, 9.7 µmol, 58%).

**HRMS** (LCMS): calcd. for $C_{76}H_{103}N_{19}O_{18}S_2^{+}$: 817.8661 $[M+2H]^{2+}$
Found: 817.8669 $[M+2H]^{2+}$.

### 2-(pyridin-2-yldisulfaneyl)ethan-1-ol (S2)

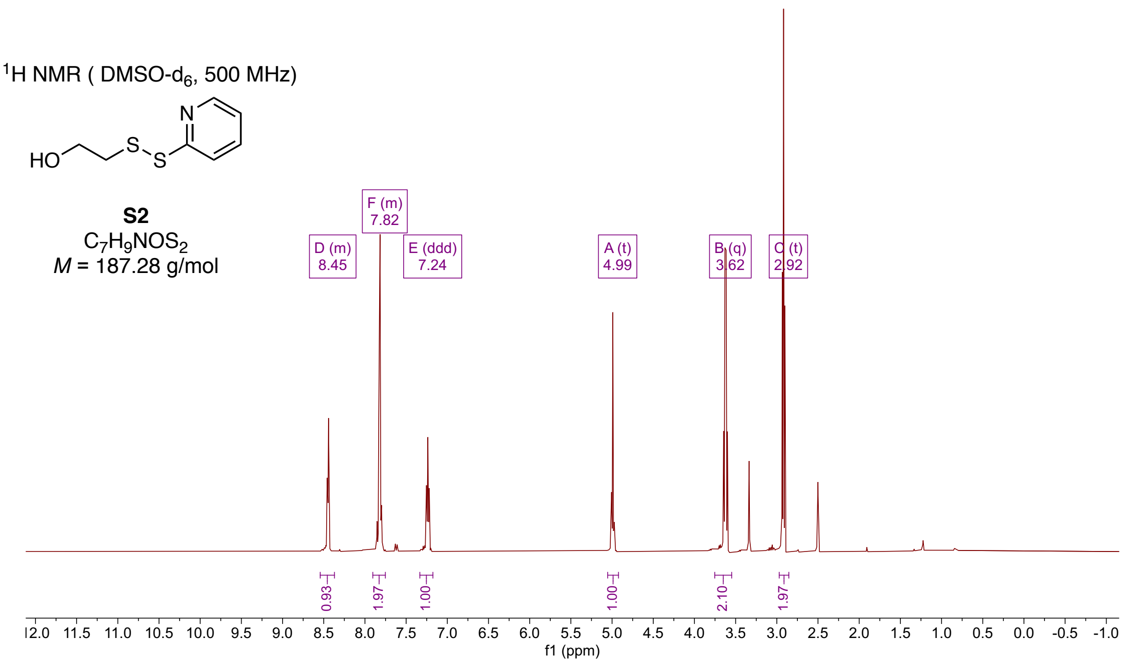


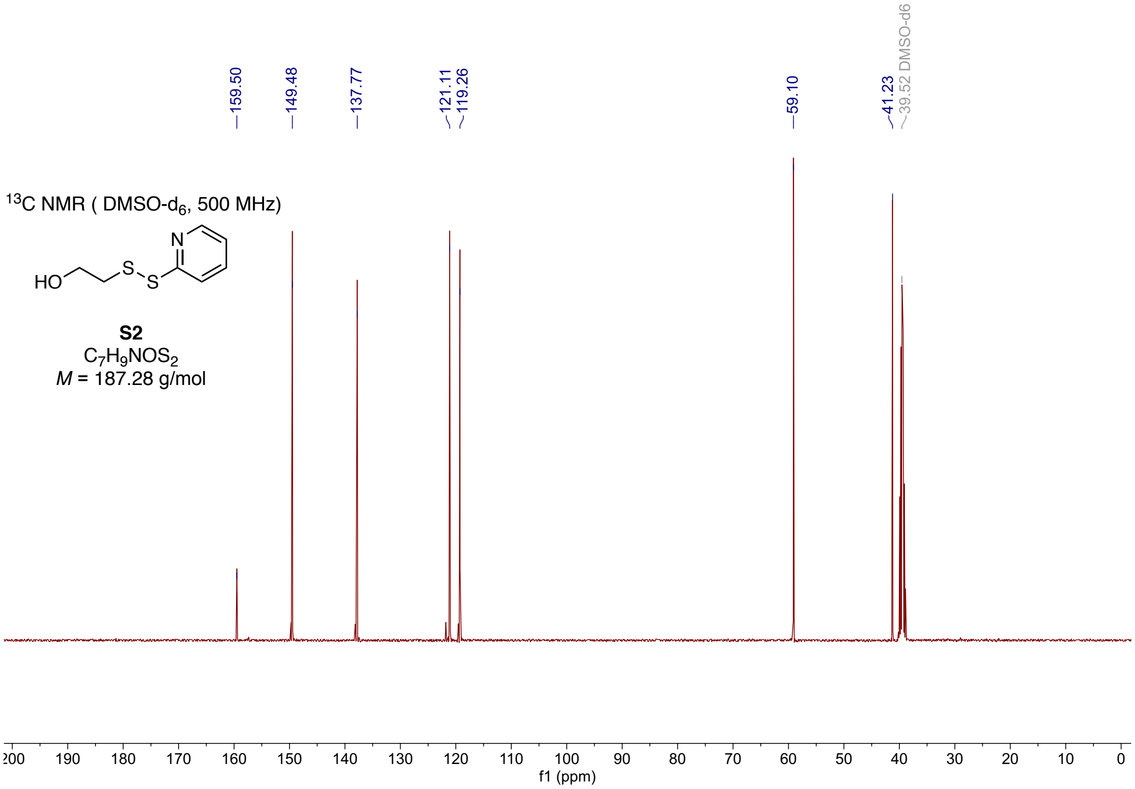

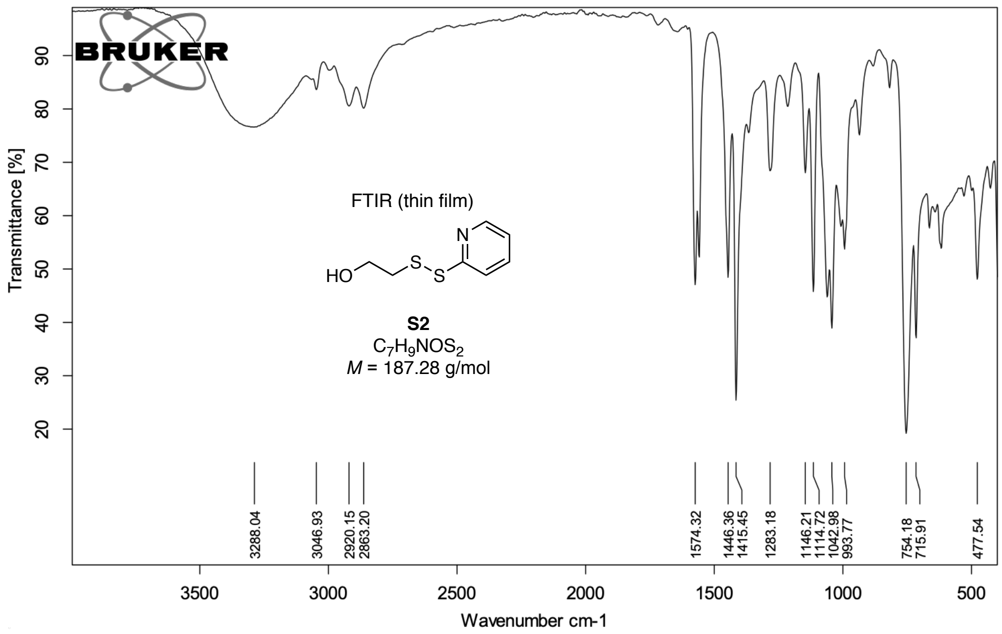
BRUKER
Transmittance [%]
FTIR (thin film)
HO
S
S
N
S2
$C_7H_9NOS_2$
$M$ = 187.28 g/mol
3288.04
3046.93
2920.15
2863.20
1574.32
1446.36
1415.45
1283.18
1146.21
1114.72
1042.98
993.77
754.18
715.91
477.54
Wavenumber cm-1

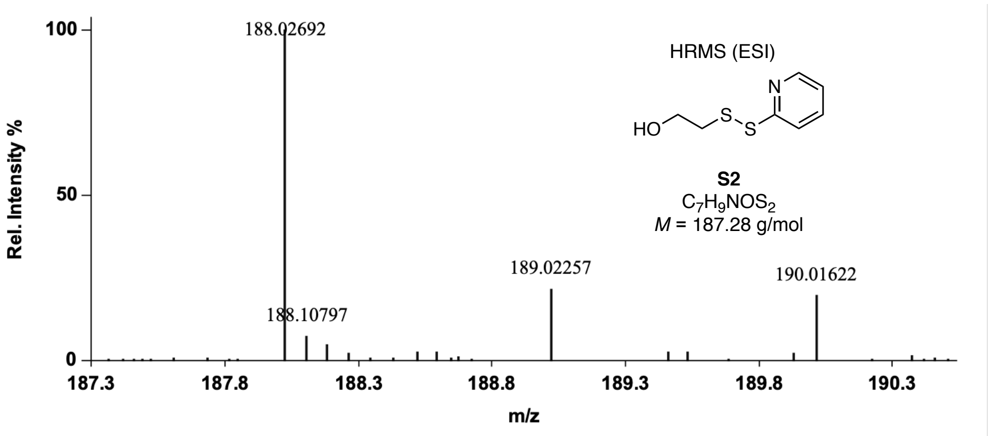
Rel. Intensity %
188.02692
HRMS (ESI)
HO
S
S
N
S2
$C_7H_9NOS_2$
$M$ = 187.28 g/mol
189.02257
190.01622
188.10797
m/z

### 2-pyridinyldithioethyl carbonate Camptothecin (S4)

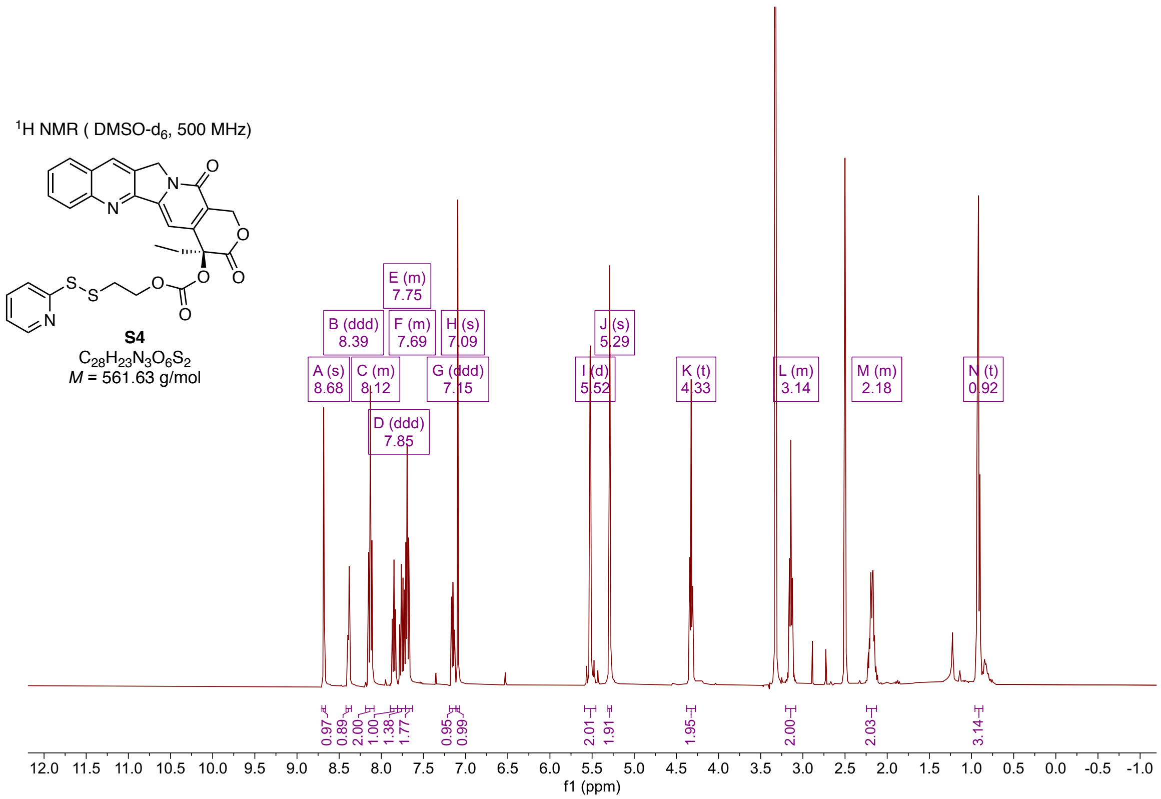


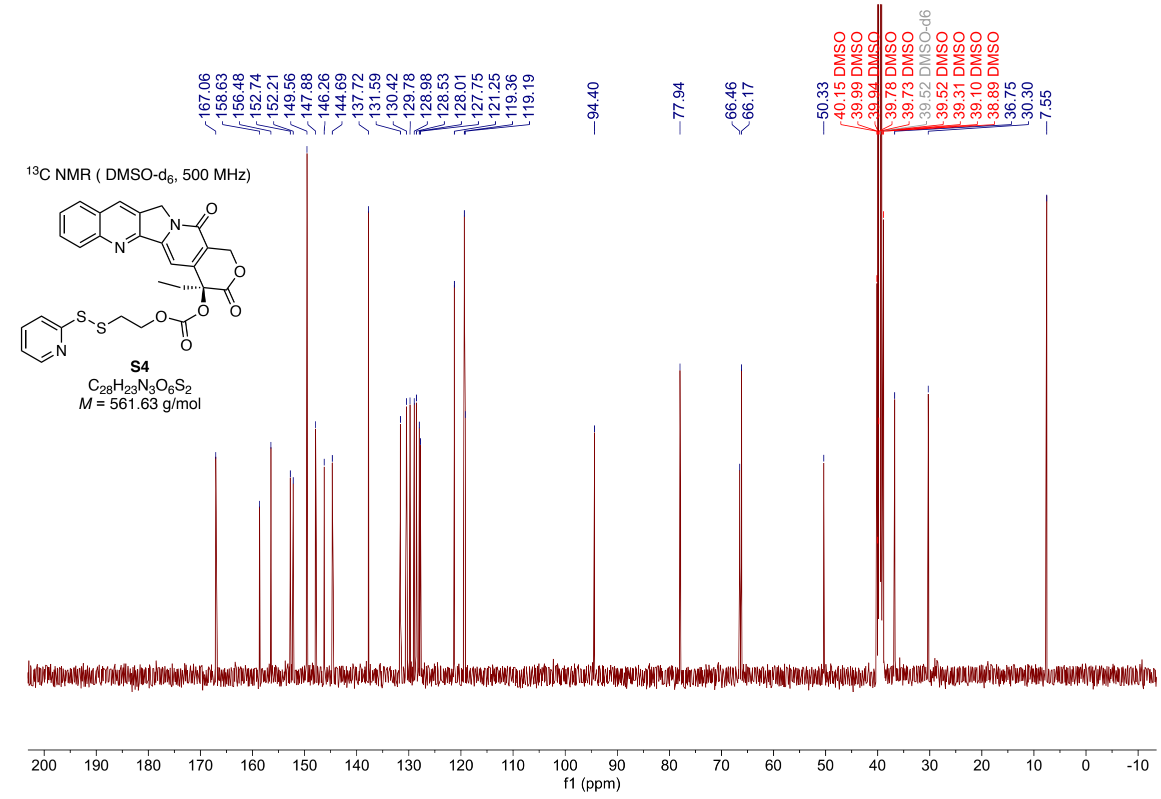

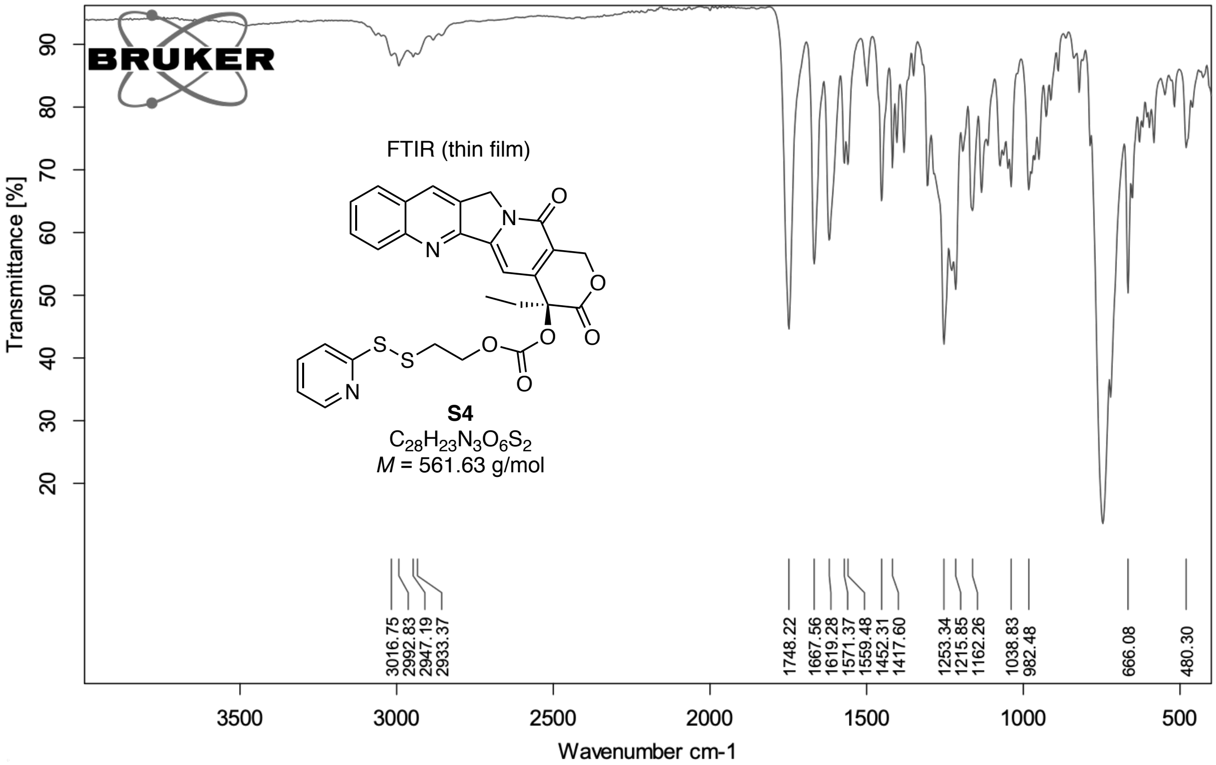


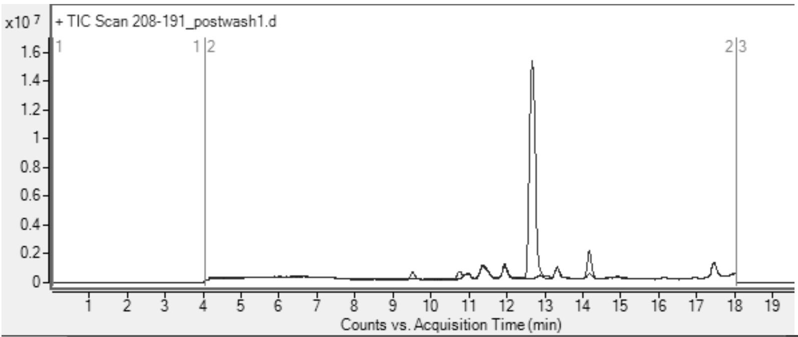


LCMS ($C_4$, 1-91% MeCN, overlay with blank chromatogram)

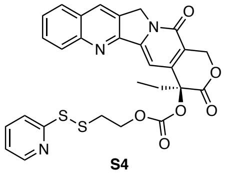

**S4**

$C_{28}H_{23}N_3O_6S_2$

$M$ = 561.63 g/mol

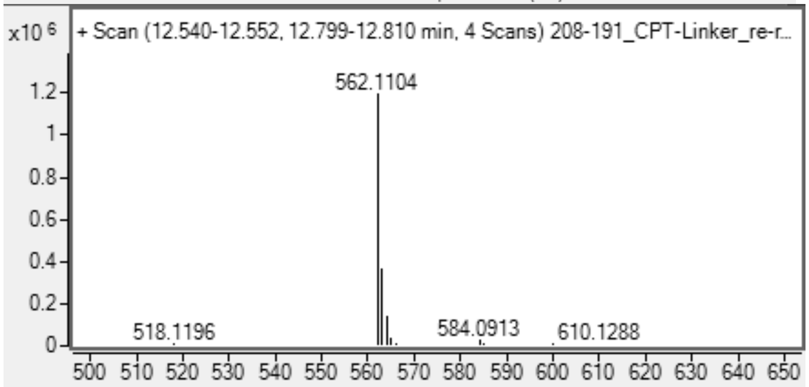


HRMS (ESI)

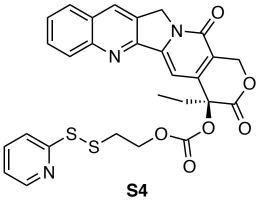

**S4**

$C_{28}H_{23}N_3O_6S_2$

$M$ = 561.63 g/mol

#### BTP-7-Campthotecin (S6)

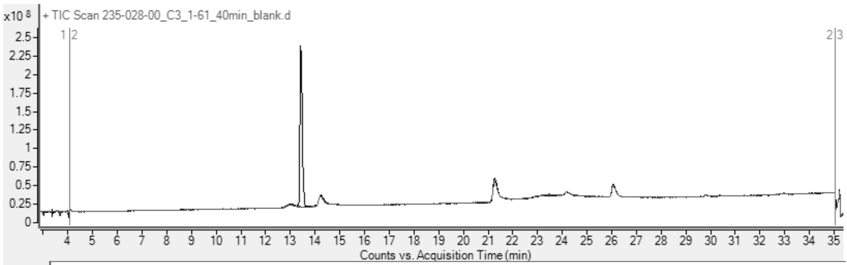


LCMS (C4, 1-61% MeCN, overlayed with blank)
BTP7-Campthotecin
**S6**
$C_{76}H_{103}N_{19}O_{18}S_2$
*M* = 1634.90 g/mol

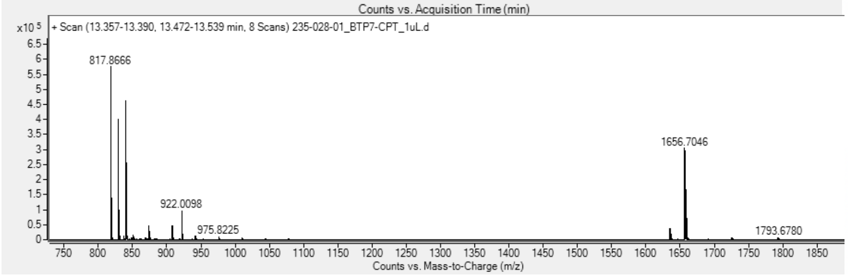


HRMS (ESI)
BTP7-Campthotecin
**S6**
$C_{76}H_{103}N_{19}O_{18}S_2$
*M* = 1634.90 g/mol

#### Scrambled BTP-7-Campthotecin (S8)

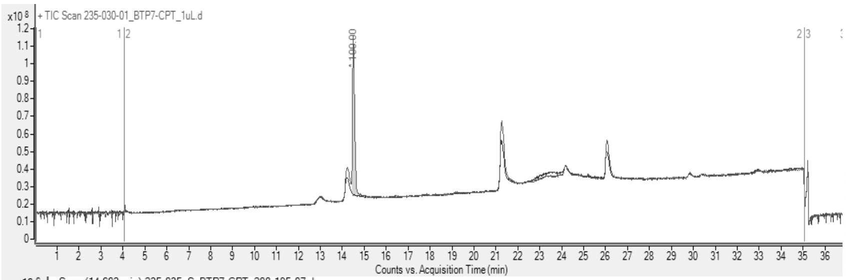


LCMS (C4, 1-61% MeCN, overlayed with blank)
Scrambled BTP7-Campthotecin
**S8**
$C_{76}H_{103}N_{19}O_{18}S_2$
*M* = 1634.90 g/mol

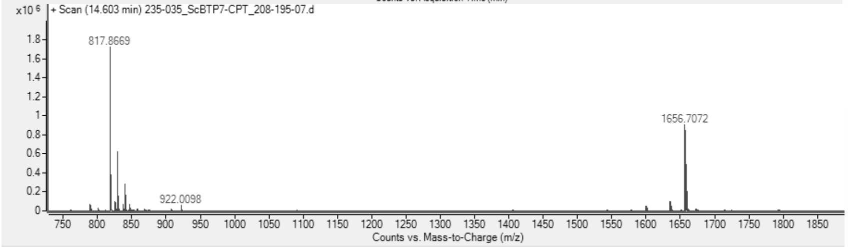


HRMS (ESI)
Scrambled BTP7-Campthotecin
**S8**
$C_{76}H_{103}N_{19}O_{18}S_2$
*M* = 1634.90 g/mol

**Octet association and dissociation curves**

FITC-BTP-7

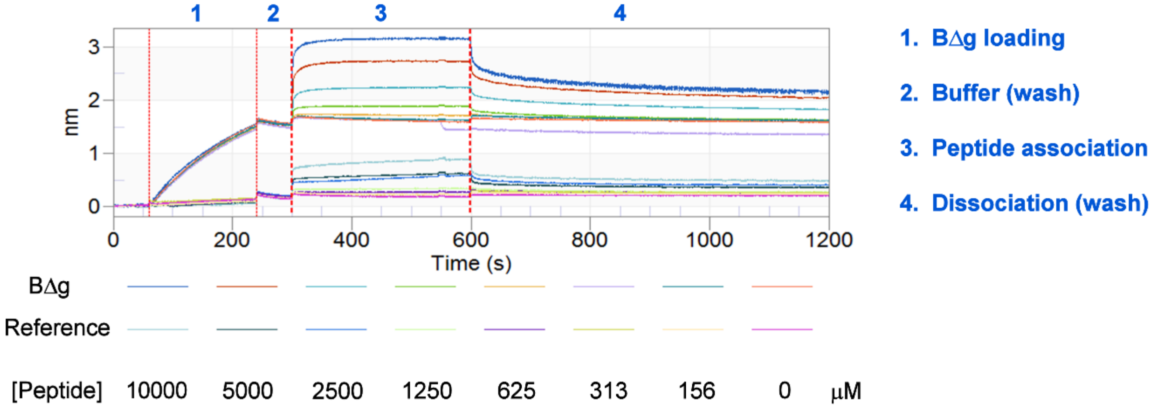


Cy5.5-BTP-7

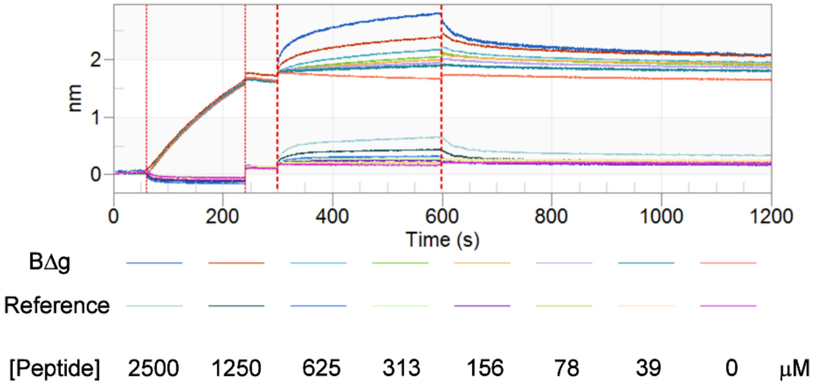


BTP-7-CPT

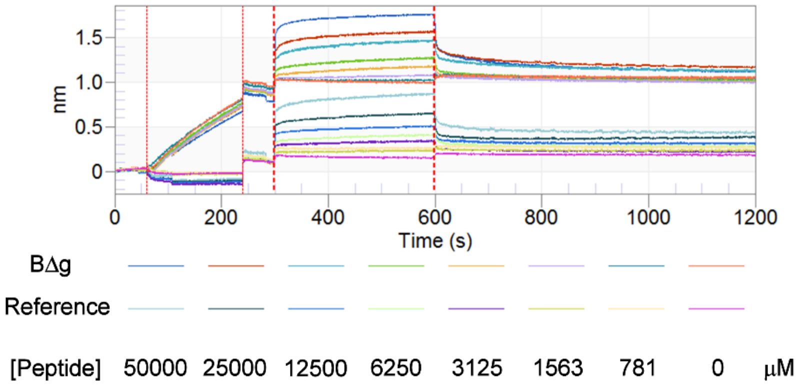
